# Comparative chemical characterisation of chitosans and their impact on growth, faecal consistency and microbiota composition in weaned piglets

**DOI:** 10.64898/2026.03.26.714014

**Authors:** Simona Di Blasio, Anouschka Middelkoop, Francesc Molist, Stefan Cord-Landwehr, Mattia Pirolo, Alhussein Abdelrahman Elrayah, Luca Guardabassi, Liam Good, Ludovic Pelligand

**Author notes:** Corresponding author: Simona Di Blasio (E-mail address).

## Abstract

Managing post-weaning diarrhoea (PWD) in piglets is difficult due to limits on antibiotics and zinc. Chitosan is emerging as a potential feed additive. We analysed a chito-oligosaccharide hydrochloride (COS-HCl), a low molecular weight (LMW) chitosan, and a medium molecular weight (MMW) chitosan, and assessed their effects on growth, faecal consistency, microbiota, and potential interference with enterotoxigenic Escherichia coli (ETEC).

The three chitosans were characterised using ¹H-NMR, SEC-RI-MS, and SEC-RI-MALLS. COS-HCl had an Mw of 0.824 kDa; LMW and MMW showed Mw ranges of 14.4 kDa (0.3-30 kDa) and 116 kDa (15-600 kDa). Degrees of acetylation were 9.5%, 6.5%, and 15%.

Two 42-day field studies evaluated average daily gain (ADG), faecal consistency, and microbiota. In the first trial, COS-HCl at 0.025–0.1% did not significantly affect ADG (-33 to - 12 g/d). In the second, LMW and MMW at 0.01% did not significantly change ADG (-7 and +3 g/d). Faecal consistency, ETEC shedding, and microbiota composition were similar to controls. An enzymatic HPLC-MS method enabled quantification of MMW chitosan in premix.

Our results highlight the importance of advanced chitosan characterisation for precision nutrition and suggest that a threshold dosemay be needed to benefit growth and gut health in PWD management.

**Graphical Abstract:** 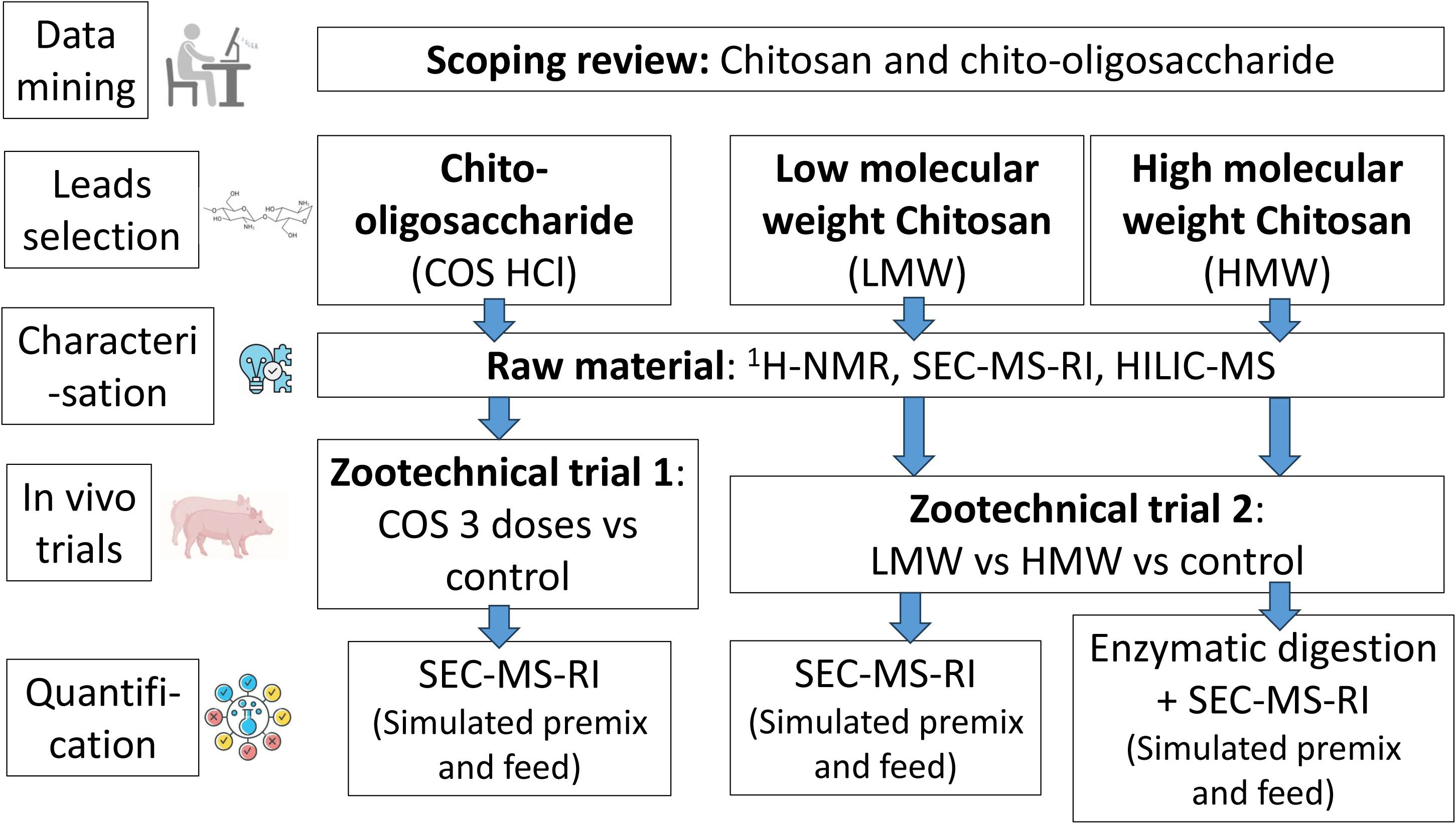

## 1. Introduction

Antimicrobial resistance in livestock negatively affects animal health and food security, and zoonotic transmission is an important public health treat. Treatment options for pigs are increasingly limited. In particular, in pigs, enterotoxigenic *Escherichia coli* (ETEC) is a primary cause of severe bacterial post-weaning diarrhoea (PWD). PWD typically affects piglets during the first two weeks after weaning. A significant factor in PWD is the separation of the piglets from the mother and an abrupt dietary change from sow’s milk to solid feed, leading to decreased feed intake and stress (Le Dividich & Sève, 2000). Associated symptoms include intestinal wall damage, immune system dysfunction, gastrointestinal problems, gut microbiota dysbiosis, and reduced growth (Gresse et al., 2017).

The disease is often treated with highest-priority and critically important antibiotics, such as colistin, which is increasingly discouraged since 2016 to reduce the risk of antimicrobial resistance to this last-resort antimicrobial in people. The use of zinc oxide, which was extensively used to control PWD in intensive animal production, has been phased out since summer 2022 (European Commission, 2017). Alternative products, such as feed additives, modified diets, improve management of the animals and new veterinary medical products are urgently needed.

Chitosan, a natural biopolymer derived from deacetylation of chitin, is a linear polysaccharide composed of β-1-4 linked D-glucosamine and N-acetyl-D-glucosamine units. Despite sharing a unique CAS-number (9012-76-4), chitosan derivatives are chemically diverse, with different degrees of de-acetylation and molecular weights ranging from high molecular weight chitosan (HMWC, >700 kDa), medium molecular weight chitosan (MMWC, 150–700 kDa), and low molecular weight chitosan (LMWC, less than 150 kDa). Furthermore, depolymerisation of chitosan results in chito-oligosaccharide (COS) formed of 2 and 20 units of glucosamine with molecular weights below 4 kDa.

Chitosan and chito-oligosaccharides (COSs) used as feed additives are reported to improve production and performance in post-weaning piglets (Swiatkiewicz et al., 2015a). While a specific mechanism of action responsible is unknown, their effects on gut flora stabilisation, improving intestinal morphology, nutrient digestibility and lowering inflammation have been widely reported (Edo et al., 2025).

As a feed additive, chitosan has several additional attractive features. Feed additives fall under EU Regulation (REGULATION (EC) No 1831/2003 Official Journal of the European Union, 2003), which requires i) characterisation and validation of quantification methods to demonstrate content homogeneity and batch stability, ii) assessment of efficacy in 3 zootechnical trials and assessment of safety of the feed additive for iii) the target species, iv) the consumer and v) the environment. Feed concentration of chitosan and chitosan oligosaccharides up to 5 g/kg (0.5%), administrated for 42 days, are well tolerated by pigs with no effect on blood parameters (Swiatkiewicz et al., 2015b). The possible presence of chitosan and chitosan oligosaccharides in pig meat is likely to be safe for the consumer as intestinal absorption is limited to smaller oligomers of COS (chitobiose and chitotriose) (Chen et al., 2005). Moreover, chitosans are commonly present in food and beverages (exoskeletons of crustacea, fungi (Kirk, 2008), wine and beer (Peñas et al., 2015)). Finally, chitosans are used as dietary supplements for body weight reduction in people (Trivedi et al., 2016) (Fatahi et al., 2022).

We hypothesised a beneficial effect of chitosan oligosaccharides and chitosan as zootechnical feed additives in the context of PWD. The objective was to characterise them analytically and evaluate their impact on growth performance, faecal consistency and microbiota composition, and ETEC shedding. Here we report the selection of the three chitosans for comparative testing, challenges faced in developing analytical methods to characterize, detect, and quantify their composition in premix and feed and the outcome of two *in vivo* trials assessing zootechnical performance. These analytical results are encouraging ahead of a PWD challenge study, which will be reported separately.

## 2. Materials and methods

### 2.1. Scoping review for rationale chitosan selection

Published research was semi-systematically assessed, collecting information about the chitosan molecular weight, deacetylation degree, feed inclusion levels and zootechnical effects on weaned piglets. Search terms (“Chitosan AND piglets” and “COS AND piglets”) were expanded to include, “pig,” “chito-oligosaccharide,” and “NOT nanoparticles” were applied to 4 libraries: Pubmed, Science Direct, Abbvie ONE Library and RVC SCOUT. Abstracts of 2,126 references were screened, and full-text articles reporting growth performance data (ADG, FCR, ADFI, BWG) versus a control diet were included in a random-effects scoping review for ADG. Plots were created using GraphPad Prism v10.3.2.

### 2.2. Procurement

Based on the semi-systematic review results and EFSA feed quality requirements, promising chitosans, including chito-oligosaccharide hydrochloride (COS-HCl), low molecular weight (LMW) and medium molecular weight chitosans (MMW), with different molecular weights (expressed in kDaltons) and degree of polymerisation were sourced as GMP-grade. Chito-oligosaccharide hydrochloride (COS-HCl, <3kDa, DP: 2–10) and medium molecular weight chitosan (MMW, Mw 116 kDa) were purchased from Aoxin (Jiangsu Aoxin Biotechnology Co., Ltd, Shanghai, China). Low molecular weight (LMW, Mw 14.4 kDa) was purchased from Korui (Jiaxing Korui Biotech Co. Ltd, Zhejiang, China)).

### 2.3. Analytical characterisation of leads

#### Proton Nuclear Magnetic Resonance (^1^H-NMR): all chitosans

The degree of acetylation (DA) was determined using proton NMR (^1^H-NMR) (see supplementary method M1) *Chito-oligosaccharide Hydrochloride characterisation: Size exclusion chromatography, refractive index detection, mass spectroscopy (SEC-RI-MS)* Molecular Weight (Mw), Number Average Molecular Weight (Mn), and degree of polymerisation (DP) were determined by Size Exclusion Chromatography (SEC) coupled with Refractive Index (RI) and Mass spectrometry (MS) detection (SEC-RI-MS) (see supplementary method M2).

#### Chito-oligosaccharide Hydrochloride: Characterisation of the composition

Hydrophilic interaction liquid chromatography-evaporative light scattering detector-Mass Spectrometry (HILIC-ELSD-MS system) was firstly employed to qualitatively reveal small oligomers and the most prevalent in COS HCl. (see supplementary method M3), but the actual oligomeric composition of COS HCl was determined by Size Exclusion Chromatography (SEC) coupled with Refractive Index (RI) and Mass spectrometry (MS) detection (SEC-RI-MS) (see supplementary method M2).

#### LMW and MMW chitosan: Characterisation of the composition

Weight averaged Molecular Weight (Mw), degree of polymerisation (DP) and Dispersity (D) were determined by *High-performance* Size Exclusion Chromatography (HP-SEC) coupled with Refractive Index (RI) and Multi Angle Laser Light Scattering (MALLS) detectors (see supplementary method M4).

### 2.4. In vivo zootechnical studies

#### 2.4.1. Ethical and regulatory approvals

Two 42-days zootechnical studies were conducted at Schothorst Feed Research BV (SFR, Lelystad, The Netherlands) following recommendations of Commission 2007/526/CE. Both experimental protocols (VOF-61 and VOB-62) were approved by the Animal Care and Use Committee of SFR and by RVC Clinical Research Ethical Review Board (CRERB). Since the feed additive was not registered in the EU, a dietary exposure assessment for COS and chitosan was conducted to evaluate the possible risk to meat consumers. The Medicines Evaluation Board (Utrecht, The Netherlands) approved that piglets surviving the trial could be commercially reared for meat production (slaughter > 4 month after end of trial).

#### 2.4.2. Animals and housing

Clinically healthy Tempo x TN70 piglets (Large White x Norsvin Landrace) were selected based on their sex and BW at weaning (30 days). The first trial, trial included 192 pigs (9.4 ± 0.03 kg) and the second trial 288 piglets (7.6 ± 0.03 kg).

At weaning, the piglets were moved into experimental pens with partially slatted floors, without bedding. Animals had ad libitum access to feed (feeder with 3 feeding places) and water (nipple drinker). Enrichment was provided (cotton rope, metal chain with plastic bar and floor-based rubber toy (Easyfix Luna, MS Schippers, The Netherlands). In the first trial piglets were housed in groups of four (1:1 male:female ratio) in 2.00 x 1.13 m pens. In the second trial piglets were housed in groups of eight (1:1 male:female ratio) in 2.60 x 1.60 m pens. Rooms were windowless and climate-controlled, starting at 29°C on the day of weaning, decreasing to 22°C by day 42. Artificial lighting was provided for 11.5 hours (6:30 AM to 6:00 PM). Veterinary interventions and medications were recorded. Piglets were excluded from the study if they required relocation to a hospital pen or if they reached a humane end-point, which triggered humane euthanasia.

#### 2.4.3. Feed

From day 0 to 14 post-weaning, the piglets were fed a Weaner I diet. On day 14 post-weaning, all piglets were abruptly switched to a Weaner II diet, which was used until day 42.

The diets were formulated to be iso-nutritive, meeting or exceeding the nutrient requirements recommended by the Dutch Centraal Veevoederbureau (CVB) for weaned piglets. They consisted of wheat, maize, barley, and soybean meal, with a medium-high crude protein level of 190 g and 170 g per kg of dry matter for Weaner I and II, respectively, and 11.0 SID lysine/kg dry matter (see Supplementary Table S1 and S2). Chitosans or COS were added into the treatment diets, under the form of a maize premix, which replaced part of the maize content. The diets were pelleted at 70°C (3 mm diameter pellets) at a specialised feed mill.

All experimental diets were analysed for the Weende components (also known as “proximate analysis”) at SFR, with moisture determined gravimetrically after oven-drying at 80°C vacuum to a constant weight (NEN-ISO 6496:1999), crude protein content measured according to the Dumas principle (NEN-EN-ISO 16634-2:2016), crude fibre determined by the filter technique (NEN-EN-ISO 6865:2001), crude fat content determined after hydrolysis with hydrochloric acid under heating (NEN-ISO 6492:1999), and ash was measured gravimetrically after washing the sample for 3 h at 550°C (NEN-ISO 5984:2003).

#### 2.4.4. Sample size calculations

For sample size calculations, Average Daily Gain (AGD) and Feed Conversion Ratio (FCR) were considered as zootechnical efficacy endpoints (EFSA FEEDAP Panel, 2018), using data from (Zhou et al., 2012), where the effect of supplementing the diet with COS at the inclusion of 0.1 and 0.2% was tested on weaner pig performance at 42 post-weaning. Between day 0-42 post-weaning, ADG was 41 g/day higher (standard error of the mean 12.4) and gain:feed was 0.037 g/g (SEM 0.01) higher in the treated group compared to the control group. Following EFSA power calculation guidelines and using GenStat 21^st^ ed., we needed 12 and 9 replication pens for the ADG and gain:feed endpoints, respectively to achieve 80% power and significance level of 0.05 computed for. See Supplementary Table S3.

#### 2.4.5. First trial experimental design

The first trial was a COS dose response with 3 inclusion levels 0.025%, 0.05% and 0.1% COS as well as a negative control diet (NC) (Table1)

Piglets were block randomised into replicates based on body weight and sex. Each of the 12 replicates consisted of 4 piglets (male:female ratio of 1:1), yielding 48 piglets/treatment followed for 42 days post-weaning. The experiment was conducted in two batches over 2 weaning rounds with 3 weeks in between. The first batch consisted of 5 replicates per treatment and the second batch consisted of 7 replicates per treatment.

#### 2.4.6. Second trial experimental design

The second trial compared two chitosans (LMW and MMW) at similar inclusion level (0.01%) against a negative control (Table 2)

**Table 1.** First SFR trial dietary treatments.

| Trt | Description | Product | Inclusion Weaner I, % | Inclusion Weaner II, % |
| --- | --- | --- | --- | --- |
| 1 | Negative control (NC) | - | 0 | 0 |
| 2 | Low dose COS | COS | 0.025 | 0.025 |
| 3 | Medium dose COS | COS | 0.05 | 0.05 |
| 4 | High dose COS | COS | 0.1 | 0.1 |

**Table 2.** Second SFR trial dietary treatments.

| Trt | Description | Product Batch nr. | Inclusion Weaner I, % | Inclusion Weaner II, % |
| --- | --- | --- | --- | --- |
| 1 | Negative control (NC) | - | 0 | 0 |
| 2 | LMW chitosan | LMW chitosan 2110123000901 | 0.01 | 0.01 |
| 3 | MMW chitosan | MMW chitosan E0820210704 | 0.01 | 0.01 |

**Table 3.** Overview of the measurements.

| Day | Body | Feed | Faecal | Other observations | Daily |
| --- | --- | --- | --- | --- | --- |
| post-weaning | weight per piglet | weight back per pen | sampling per pen |  | observations |
| 0 | X | X |  | Faecal consistency | Clinical signs, |
| 7 |  |  | X | score three times a week during day 0-14 post-weaning and twice a week between day 14-42 post-weaning on pen level | antibiotic treatments, mortality/culls, any unusual incidents or adverse events. |
| 14 | X | X | X |  |  |
| 42 | X | X |  |  |  |

**Table 4.**
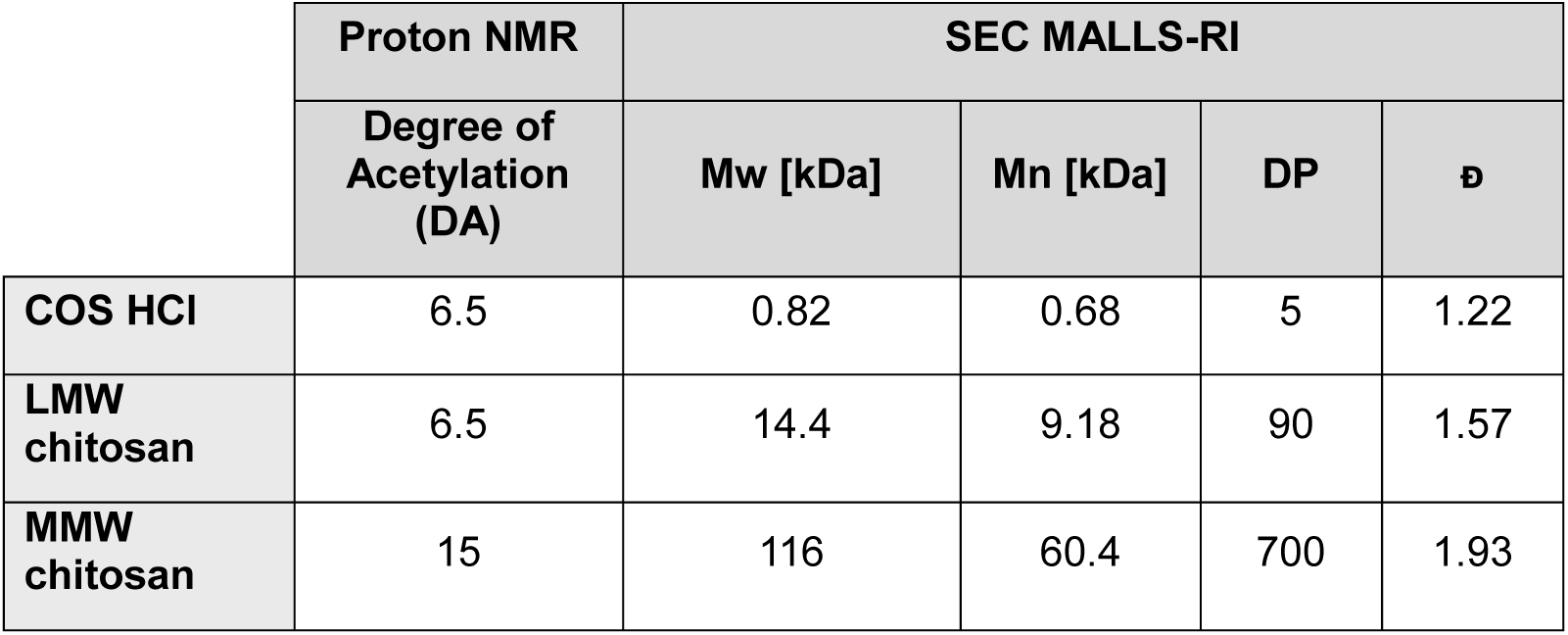
Degree of acetylation (DA), Molecular Weight (Mw), degree of polymerisation (DP), Dispersity (ᴆ) values for MMW and LMW chitosan were provided by Münster University.

Piglets were block randomised into replicates based on body weight and sex. Each of the 12 replicates consisted of 8 piglets (male:female ratio of 1:1), with 96 piglets/treatment followed for 42 days post-weaning. The first batch included 8 replicates per treatment while the second batch included the remaining 4 replicates per treatment.

#### 2.4.7. Zootechnical measurements

Individual pig body weights were recorded at weaning, day 14 and day 42 post-weaning. Average daily gain (ADG, g/pig/d), feed conversion ratio (FCR) and Daily feed intake (ADFI, g/pig/d) were calculated on a pen basis for each experimental phase (i.e. day 0-14 and day 14-42 post-weaning) and for the total weaner (i.e. day 0-42 post-weaning). Feed allowance and refusals were recorded for each pen.

Pen faecal consistency score (FS) was measured three times per week between day 0-14 post-weaning and twice weekly between day 14-42 post-weaning on an 8-point scale (Supplementary Table S4, 0 = severe water thin diarrhoea, 6 = normal faecal consistency, 8= hard, dry and lumpy faeces). FS was averaged for each experimental phase (i.e. day 0-14 and day 14-42 post-weaning) and for the total weaner (i.e. day 0-42 post-weaning)

Faeces were sampled on pen level at 7- and 14-days post-weaning. Using a gloved hand (new glove for each pen), the slats in the corner of the pen where the piglets defecate were wiped (Weber et al., 2017) to collect pooled faecal samples, including diarrheic samples. Samples were pooled in sterile cups, homogenized with a sterile tool and 2x 1 gram of faeces was stored in 2mL cryotubes at -80°C until shipment on dry ice to the University of Copenhagen (UCPH) for microbiological analysis

#### 2.4.8. F4-ETEC quantification

Quantitative PCR (qPCR) amplifying on the *faeG* gene (F4ac) was used to quantify F4-ETEC in 1:10 dilution of faecal samples on the LightCycler 96 System (Roche Life Science, Copenhagen, Denmark), as previously described (Larsen et al., 2023). Copy number in each sample was calculated using the equation: copy number = [10^(−1/S)]^(I−Ct), where S is the slope of the log-linear part of the standard curve, I the intercept of the standard curve, and Ct is the cycle threshold of the sample. Copies were normalized for gram of faeces and the limit of detection was set to 36 Ct, equal to ∼100 F4ac copies/reaction. The mean *faeG* copy numbers between groups were compared with the Wilcoxon Rank Sum test.

#### 2.4.9. Faecal metataxonomic sequencing

Total DNA from faecal samples collected at day 7 and 14 post-weaning was extracted using the QIAamp UCP Pathogen MiniKit (QIAGEN, Copenhagen, Denmark), with the addition of a bead-beating step using the Pathogen Lysis tube S (QIAGEN, Copenhagen, Denmark), and the inclusion of two blank extraction controls. The V3-V4 region of the 16S rRNA gene was amplified using the Quick-16S NGS Library Prep Kit (Zymo Research, CA USA), as previously described (Pirolo et al., 2023). Negative and positive control (ZymoBIOMICS DNase/RNase Free Water and ZymoBIOMICS Microbial Community DNA Standard, respectively) were included in library preparation. Sequencing was performed on an Illumina MiSeq platform (2 × 300 bp paired end reads) using the MiSeq Reagent Kit v3 (600 cycles; Illumina), according to manufacturer’s instructions. The 16S rRNA sequencing data have been submitted to the NCBI Sequence Read Archive (SRA) under BioProject PRJNA1260772.

### 2.5. Quantification of LMW and MMW chitosan or chitosan HCl in premix and feed

To evaluate stability and homogeneity of chitosan or COS in diet, representative samples from each premix and feed were sent to Münster University for chitosan or COS quantification. The analytical methods used to characterize the samples are described below.

#### 2.5.1. SEC-RI-MS (COS HCl, COS premix and COS feed analysis)

The matrix effect of maize and feed on the detection and quantification of COS was evaluated using method M2 with the following modifications. Simulated samples were prepared in 150 mM ammonium acetate buffer (pH 4.2): COS alone (final concentration 1 mg/mL); premix (COS 1 mg/mL and maize 4 mg/mL) and COS feed sample (COS 1 mg/mL with maize 4 mg/mL and feed 10 mg/mL). Additionally different COS concentrations (0.025 – 5 mg/mL) were mixed with a constant amount of maize (4 mg/mL) and maize (4 mg/mL) and feed (10 mg/mL) to further evaluate the matrix effect. All samples were filtered through 0.2 µm modified nylon filters (VWR, Radnor, DE, USA) and injected as described previously.

#### 2.5.2. Enzymatic method/ UHPLC-ESI-MS (Ultra-High Performance Liquid Chromatography-Electrospray Ionization-Mass Spectrometry) - detection and quantification of chitosan breakdown products)

##### Enzymatic method 1

The chitosan content and its degree of acetylation in feed samples were quantified based on the method by Urs et al., 2023, with modifications.

The method involves enzymatic depolymerisation coupled to a Mass Spectrometric Analysis, provides information about the content of glucosamine and N-acetyl glucosamine units, and degree of acetylation (DA) of the chitosan in the sample. 1% of each chitosan was mixed to maize. Subsequently the mixture underwent enzymatic digestion with chitinases and chitosanases before being analysed by UHPLC-MS.

In detail, the sample was homogenized using steel balls, followed by chemical *N*-acetylation with deuterated acetic anhydride in Sodium bicarbonate buffer. This reaction proceeded under room temperature mixing, followed by heating at 100°C for complete acetylation. After centrifugation and washing, the pellet was resuspended in ammonium acetate buffer. Enzymatic digestion was conducted using *Trichoderma viride* chitinase (Sigma-Aldrich, St-Louis MO, USA), ChiB (*Serratia marcescens* chitinase ChiB), and CSN174 (*Streptomyces* sp. chitosanase CSN-174) at 37°C for three days. The digested samples were freeze-dried, resuspended in water, filtered, and mixed with an internal standard before UHPLC-MS analysis (see supplementary method M5).

##### Enzymatic method 2

Samples (5 mg/mL) were incubated with ChiB and CSN174 at 37°C for 16 hours, then centrifuged to remove insoluble material. The supernatant was filtered, N-acetylated with deuterated acetic anhydride in Sodium bicarbonate buffer, freeze-dried, and digested with *Trichoderma viride* chitinase for three days. Filtered samples were mixed with an internal standard and analysed via UHPLC-MS (see supplementary method M6).

#### 2.5.3. Ultrahigh performance liquid chromatography–electrospray ionization mass spectrometry (UHPLC-ESI-MS)

UHPLC-ESI-MS was used to determine the average Degree of Acetylation by quantitatively determining the amount of GlcNAc and GlcN units present in the chitosan sample (Hamer et al., 2015). All measurements were carried out using a Dionex Ultimate 3000RS UHPLC system (Thermo Scientific, Milford, MA, USA) coupled to an amaZon speed ESI-MS^n^ detector (Bruker Daltonik, Bremen, Germany). The reaction products were separated by a VanGuard precolumn (1.7 μm, 2.1 × 5 mm) and an Acquity UHPLC BEH amide column (1.7 μm, 2.1 × 150 mm; Waters Corp., Milford, MA, USA) through Hydrophilic Liquid Interaction Chromatography (HILIC) with solvent A (80% ACN and 20% H_2_O) and solvent B (20% ACN and 80% H_2_O), both containing 10 mM NH_4_HCO_2_ and 0.1% (v/v) HCOOH. Starting with 100% solvent A (0.0–3.0 min), solvent B gradient was increased from 0 % to 85 % (3.0–4.5 min), followed by 85% solvent B (4.5–5.5 min), then decreasing from 85% to 0% solvent B (5.5–5.7 min), finally reaching 100% of solvent A (5.7–6.7 min). The measurements were performed at a column oven temperature of 35 ^◦^C, 0.4 mL/min flow rate, and a scan range of m/z 50–2000. One μL of the sample premixed with a known amount of internal standard R*1 was injected and the resulting data were analysed via peak areas using Data Analysis v4.1 software (Bruker Daltonik, Bremen, Germany) from (Urs et al., 2023)

### 2.6. Statistical analyses

#### 2.6.1. In vivo animal zootechnical trials

The study used a randomized complete block design, with the pen as the experimental unit for statistical purposes to determine efficacy on zootechnical performance (BW, ADG, ADFI, FCR, FS). Blocking was based on room and batch.

After completing the in-life phase of the experiment, the raw data were entered into a database and verified without any changes.

The experimental data were analysed using one-way ANOVA via the General Linear Model function in GenStat^®^ Version 24 for Windows™ (VSN International Ltd, Hemel Hempstead, UK). The analyses were run with and without excluding outliers. Significant differences are declared at *P*≤0.05, with near significant trends as 0.05<*P*≤0.10. (Near) Significant fixed effects were further analysed by Least Significant Differences (LSD, Fisher’s LSD method) to compare treatment means. Multiple comparisons between treatment groups were tested with Tukey adjustment. Data are presented as absolute means. The *P*-value and standard error of the mean (SEM) are reported per response parameter. A polynomial regression analysis (POL) was also performed in GenStat^®^ Version 24 to investigate the relationship between dose and response. Linear and quadratic terms for the dose were included in the model.

Descriptive statistics of the data are given in supplementary Tables S8–S13.

Data from pigs that died or were culled were excluded after their removal, but their performance was considered until their removal.

Observations were marked as outliers if the residual value exceeded 2.5 times the standard error of the residuals, and missing values were estimated using GenStat®. Outliers were identified and marked in the datasets, and analyses were conducted both with and without these outliers. If any one of the response parameters (ADG, ADFI or FCR) was identified as an outlier, all three parameters for that unit and period were marked as missing (for data excluding outliers see Supplementary Tables).

#### 2.6.2. 16S rRNA data analysis

16S rRNA sequencing data were processed in R v4.2.1 using the DADA2 v1.14.1 pipeline (Benjamin J Callahan, 2016) and FIGARO v3.0 (Sasada et al., 2020) for the selection of optimal filtering and trimming parameters. After exclusion of potential contaminants using control samples using decontam v.1.12.0 (Davis et al., 2018) the resulting amplicon sequence variant (ASV) were taxonomically assigned using the Silva database v.138.1 for DADA2 (Quast et al., 2013) and only sequences assigned to Bacteria were retained. The final phyloseq object was constructed using phyloseq v1.30.0 (McMurdie and Holmes, 2013), reads were transformed using the Cumulative sum scaling (CSS) method and ASVs with ≥ 10 reads in ≥ 2 samples were retained for subsequent analysis. Alpha-diversity (Chao1, Shannon and Simpson indexes) and beta-diversity (Bray–Curtis dissimilarity metric) indexes were calculated using R package vegan. Comparison of alpha-diversity indexes was performed using the Wilcoxon Rank Sum test and p-values were corrected for multiple comparisons using Holm’s correction. Beta-diversity was visualized using a Principal Coordinates Analysis (PCoA) plot, and differences in beta-diversity were estimated by permutational multivariate analysis of variance (PERMANOVA) using the Adonis function.

## 3. Results

### 3.1. Semi systematic review, forest plot and chitosan selection

Twenty-one peer-reviewed *in vivo* studies were identified between April 2007 to September 2020 (Walsh, Sweeney, Bahar, Flynn, et al., 2013) (Wan et al., 2017) (Zhao et al., 2017) (Liu et al., 2008) (Liu et al., 2010) (Xiong et al., 2015) (Yin, 2008) (Yang et al., 2012) (Zhou et al., 2012) (Chen et al., 2009) (Yan and Kim, 2011) (Egan et al., 2015) (Xu et al., 2018) (Han et al., 2007) (Hu et al., 2018) (Duan et al., 2020) (Oliveira et al., 2017) (Wang et al., 2009) (Walsh, Sweeney, Bahar, O’Doherty et al. 2013) (Xu et al., 2013). A forest plot summarised the point estimate and confidence interval of ADG difference (chitosan or COS versus NC) for each comparison within study (Figure S1). Dietary inclusion levels ranged between 0.003 and 0.5%, molecular weight ranged between 1 and 232 kilo Daltons.

Twelve studies or sub-studies demonstrated significant increase in “average daily gain” (ADG) for chitosan on COS compared to negative control (Wan 2017, Yang 2012, Yin 2028, Zhou 2012, Xu 2018, Han 2007a, Hu 2018 Oliveira 2017, Xu 2013), two significant decreases (Egan 2015, Walsh 2013a) and results were equivocal for other studies.

By grouping the products based on their molecular weight and efficacy, it emerged that the low molecular weight chitosan used by Yin et al (2008) scored the highest ADG value (62 g/d), followed by the high molecular weight chitosan used by Xu et al. (2018), ADG 56 g/d. Among the chito-oligosaccharides present in the list, COS used by Wan et al. 2017 and COS used by Zhou et al 2012 scored the highest ADG values, 41 and 52 g/d respectively.

Considering these results, we decided to initially test a GMP food-grade water soluble COS hydrochloride, which had the potential to be better characterised than heterogeneous chitosan with high molecular weight. The initial trial sought to determine the optimal dose by testing three different COS HCl mid-range inclusion levels: 0.025%, 0.05%, and 0.1%, against basal diet. Subsequently, since the first trial did not show zootechnical improvements (see 1^st^ trial results – Table 5) a low molecular weight (LMW) chitosan and a medium molecular weight (MMW) chitosan were chosen to be tested in a second efficacy trial. The low molecular weight chitosan was a feed grade product which was already used in *in vivo* studies at 0.005% and 0.01% inclusion level (Hu 2018 and Zhang 2020) and commercialised as a feed additive for livestock in China by Korui (Jiaxing Korui Biotech Co. Ltd). The MMW chitosan was purchased by Aoxin (Jiangsu Aoxin Biotechnology Co., Ltd) as food grade and GMP certified product.

**Table 5.**
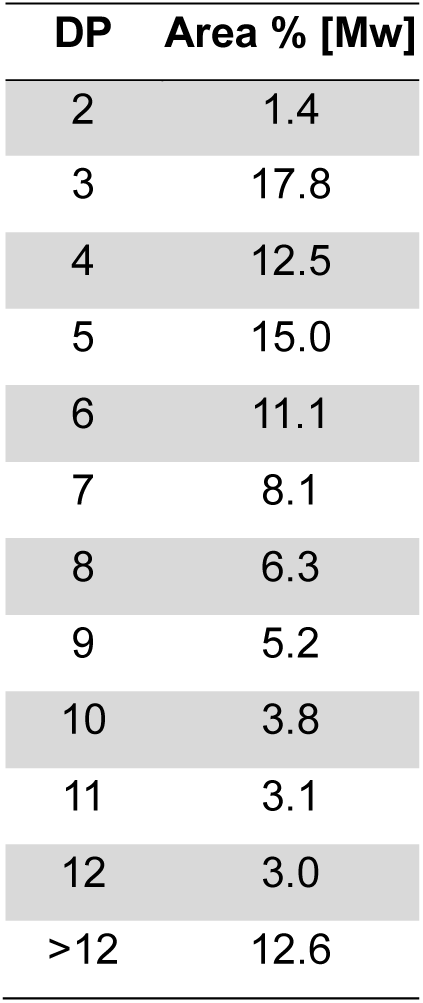
Relative peak areas and corresponding MW of the different oligomers of our sample of COS-HCl.

| DP | Area % [Mw] |
| --- | --- |
| 2 | 1.4 |
| 3 | 17.8 |
| 4 | 12.5 |
| 5 | 15.0 |
| 6 | 11.1 |
| 7 | 8.1 |
| 8 | 6.3 |
| 9 | 5.2 |
| 10 | 3.8 |
| 11 | 3.1 |
| 12 | 3.0 |
| >12 | 12.6 |

**Table 6.**
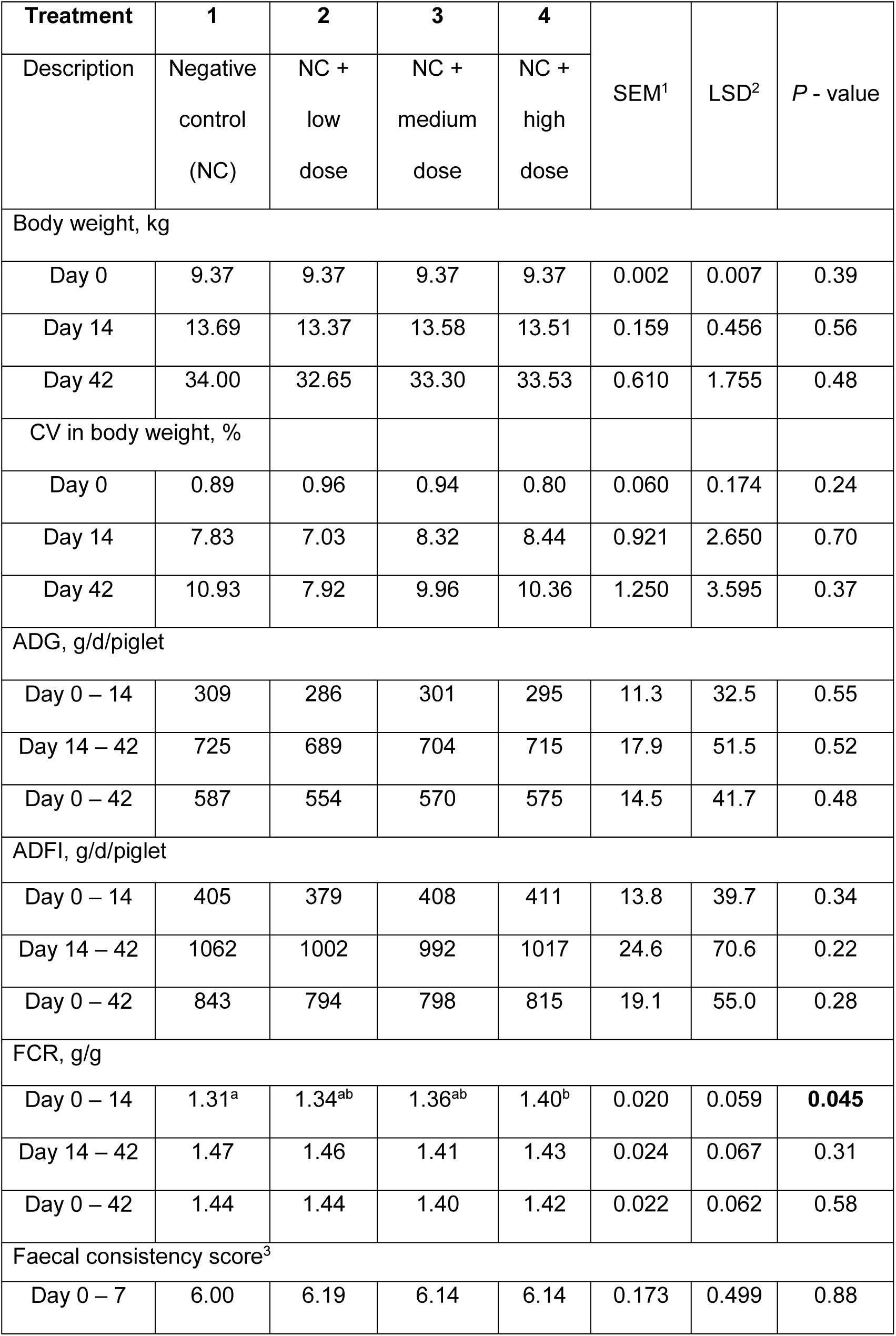
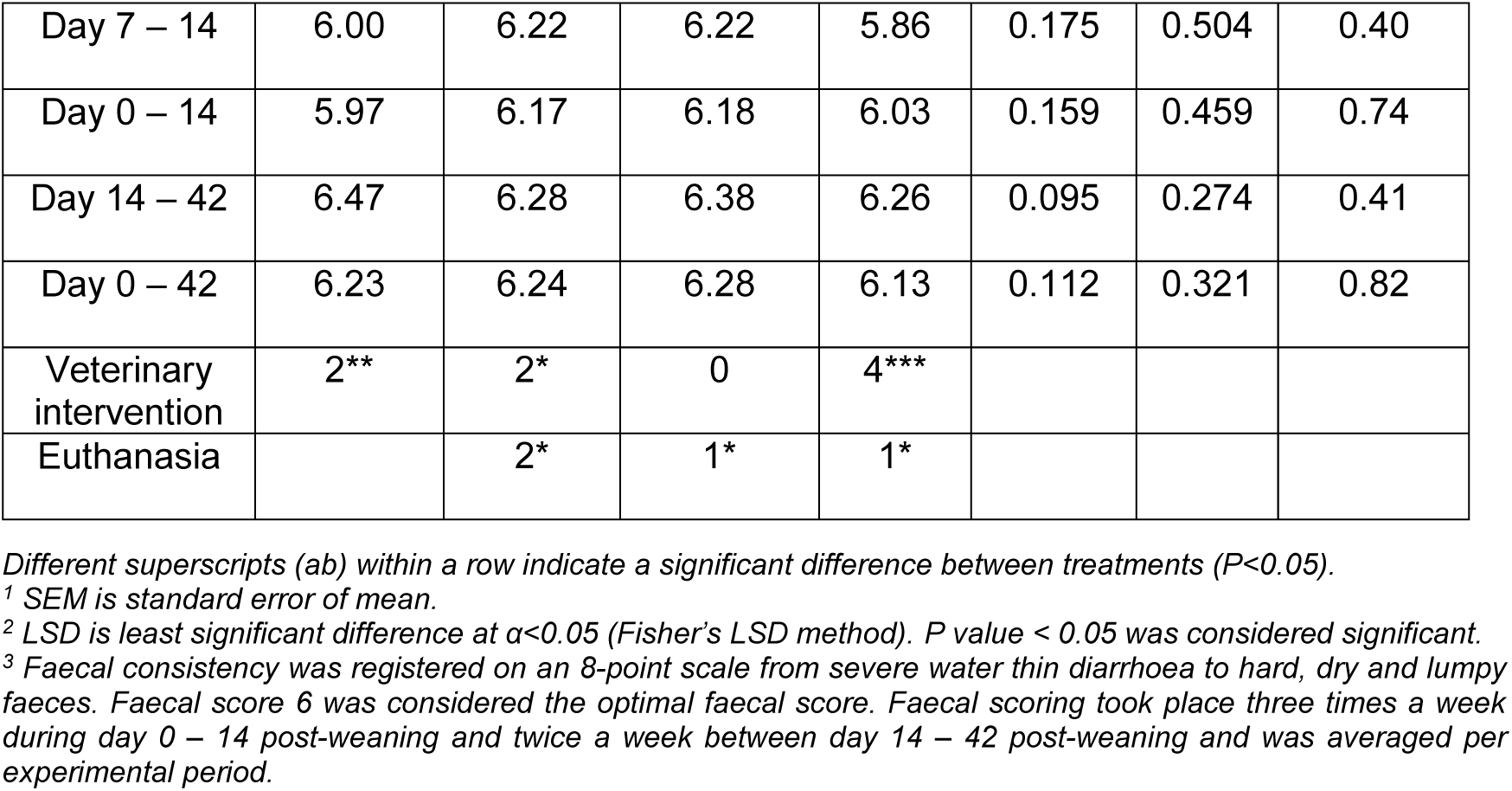
Effect of experimental diet on piglet performance between day 0-42 post-weaning (including outliers), veterinary interventions)

### 2.1. Characterisation results

#### 2.1.1. Chitosan hydrochloride

COS HCl occurs as a mixture of oligomers; the manufacturer (Aoxin) specifies Average Molecular Weight (MW) of the oligomers <3000 Daltons (Da), a degree of polymerisation (DP) between 2 - 10 and deacetylation degree ≥ 90%.

The degree of acetylation (DA) was determined by ^1^H-NMR spectroscopy (Supplementary Figure 1), whereas degree of polymerisation (DP), Number Average Molecular Weight (Mn), (Mw), and polymer composition of COS HCl was determined by Size Exclusion Chromatography (SEC) coupled with Refractive Index (RI) and Mass spectrometry (MS) detection (SEC-RI-MS) (see supplementary method M2).

To characterise the composition of COS HCl, HILIC-ELSD-MS system was firstly employed. The technique showed the small oligomers present in COS HCl (Figure 1), it gives an idea of the prevalent oligomers, but it was not a quantitative analysis.

**Figure 1.**
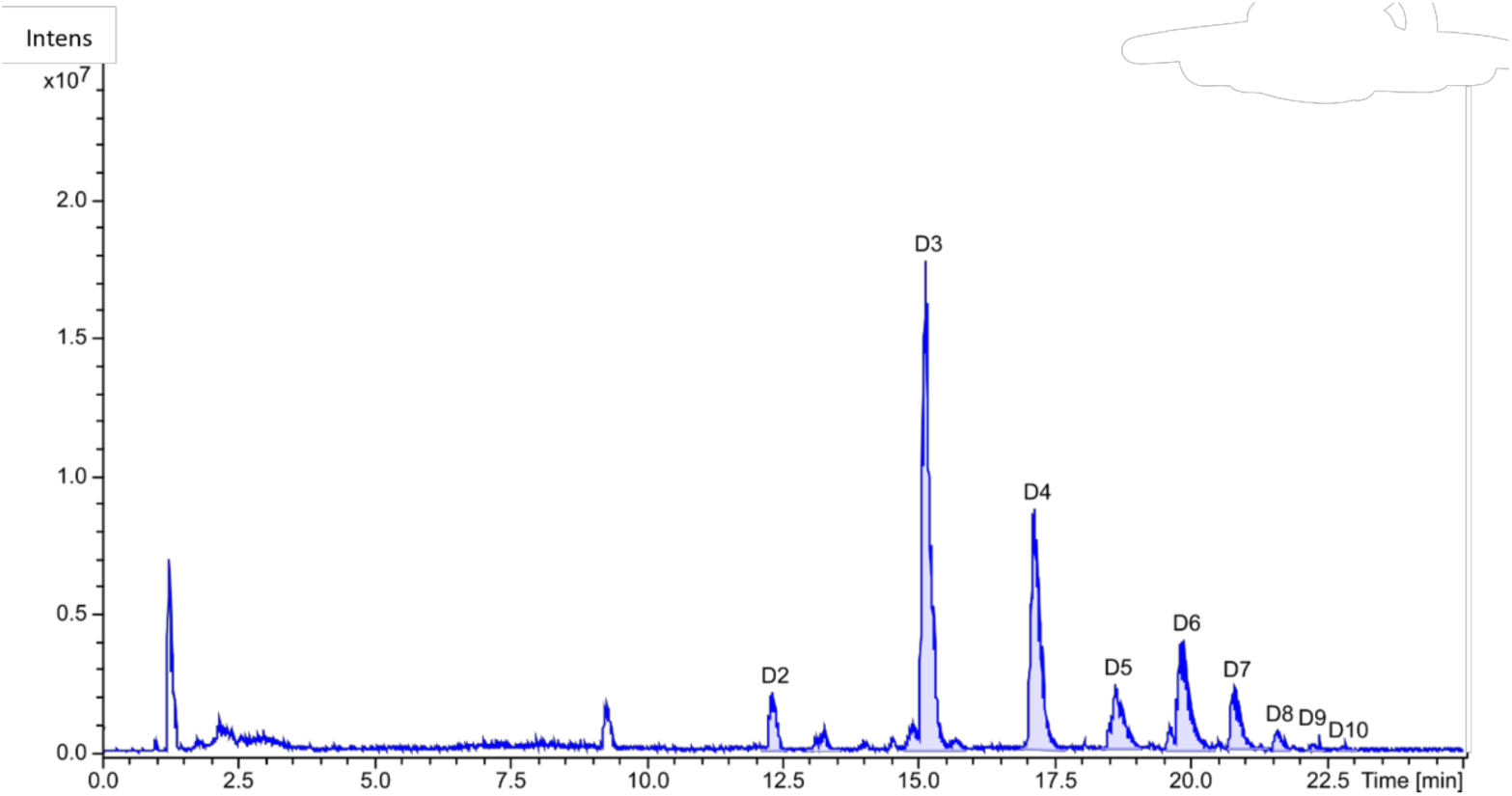
Base peak chromatogram of chito-oligosaccharide hydrochloride (COS HCl) analysed by hydrophilic interaction liquid chromatography coupled with mass spectrometry (HILIC-MS). Peak D corresponds to D-glucosamine.

To accurately detect small and larger chitosan-oligomers, a newly developed method was used (Hellmann et al, 2024), consisting of a SEC column in an Ultra High-Pressure Liquid Chromatography (UHPLC) System coupled to a refractive index detector and a mass spectrometer.

The setup allowed detection of large oligomers and even polymers which are not detectable by MS in a quantitative way by using the RI signal. This is the method of choice for characterisation of the COS spectrum, enabling determination of the relative abundance of each oligomer from the RI detector signals (mainly de-acetylated as the degree of deacetylation > 90%).

The chromatogram obtained from the refractive index detector and relative peak areas of the different oligomers (Figure 2 and Table 5) showed that COS HCl mainly consisted of short oligomers, with DP3 (chitotriose, 17.8%) and DP5 (chitopentaose, 15%) being the main components. The percentage of each DP was calculated using the area of each peak. See Suppl. Materials Figure S3 for more information about the relative percentages of each polymer and oligomer.

**Figure 2.**
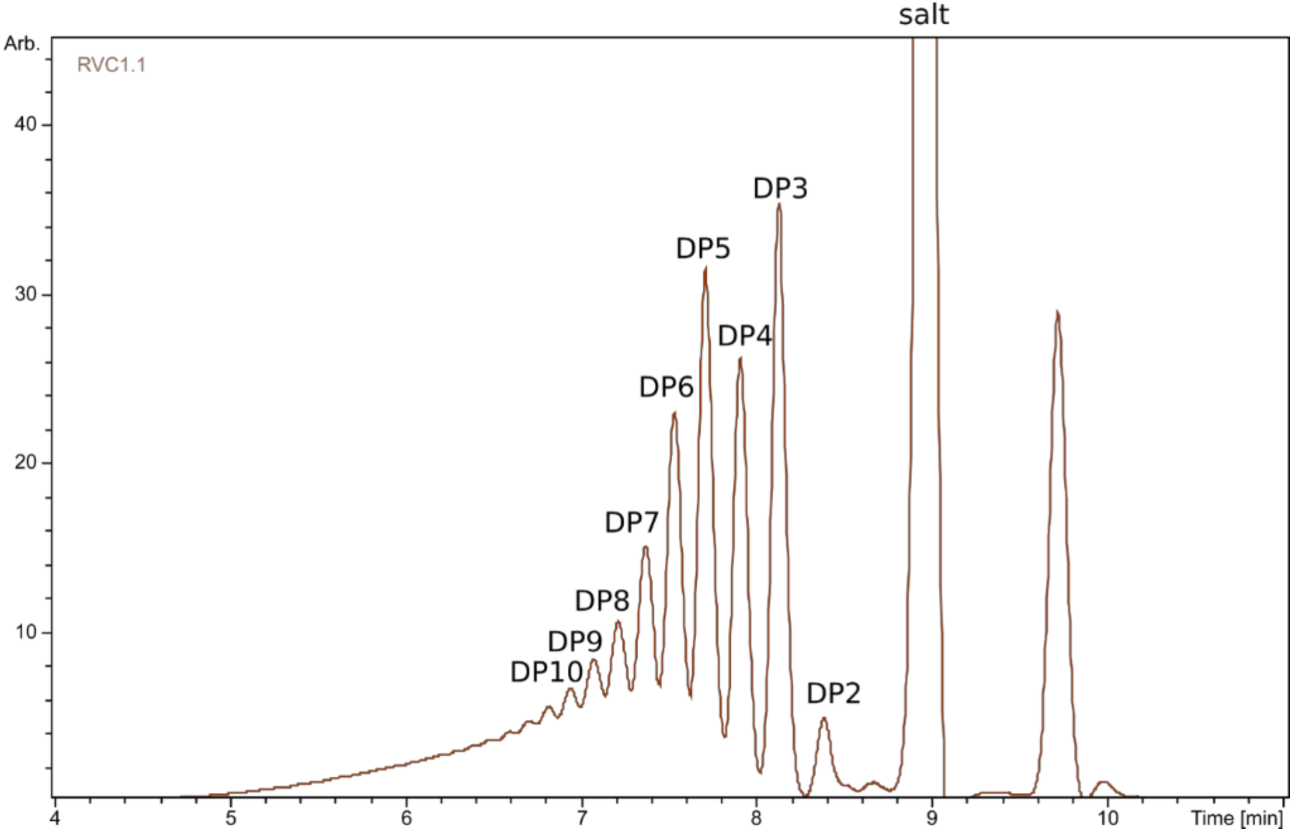
Refractive index chromatogram of chito-oligosaccharide hydrochloride (COS HCl) obtained via size-exclusion chromatography (SEC), illustrating the distribution of oligomers. In SEC, larger oligomers elute earlier than smaller ones due to their reduced interaction with the stationary phase.

**Figure 3.**
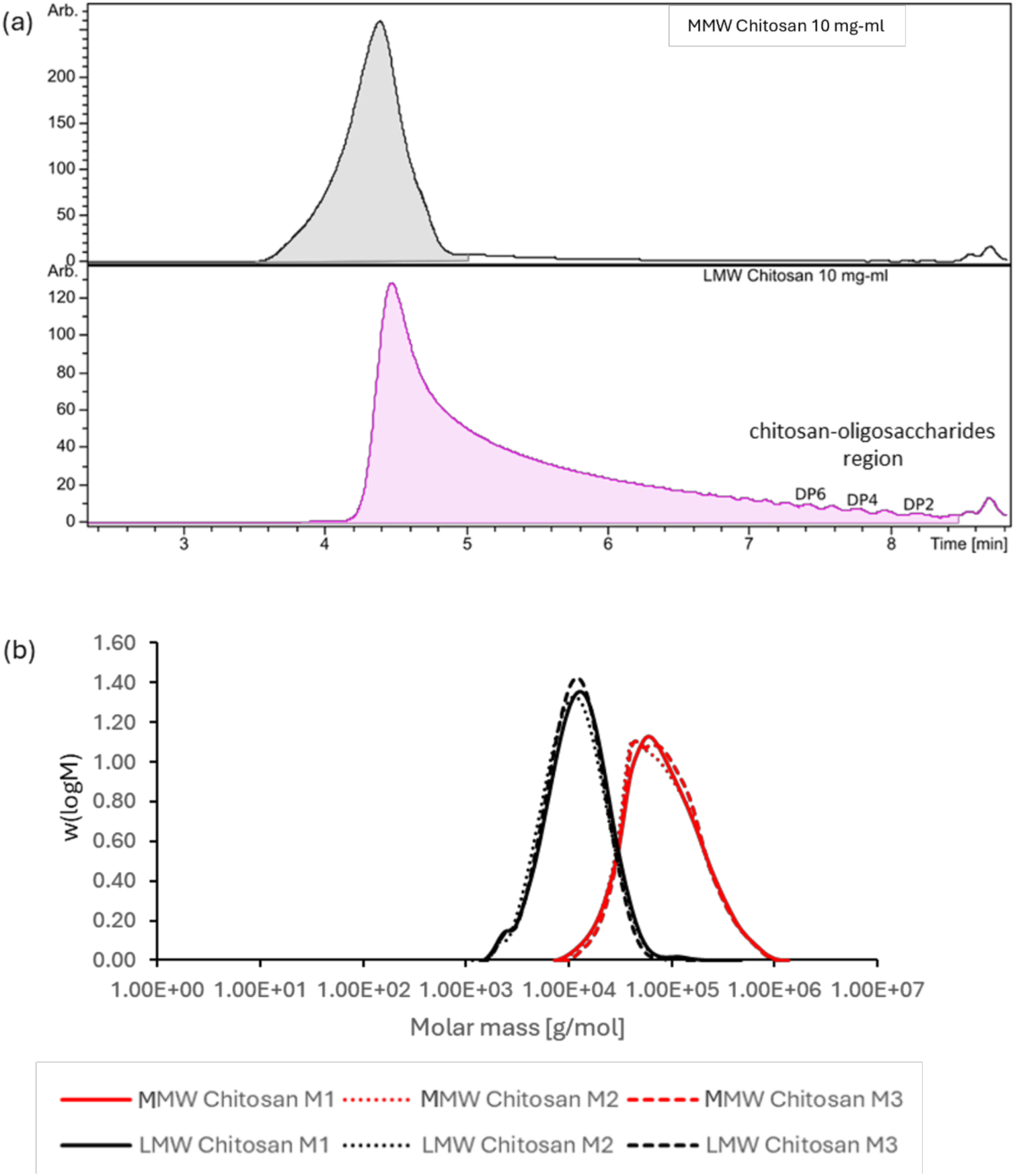
(a) Size-exclusion chromatography with refractive index and multi-angle light scattering detection (SEC-RI-MALS) chromatograms for medium molecular weight (MMW, top) and low molecular weight (LMW, bottom) chitosan. (b) Molar mass distribution profiles of medium molecular weight (MMW) and low molecular weight (LMW) chitosan, illustrating differences in polymer size between the two samples.

#### 2.1.2. Low Molecular weight and Medium Molecular weight chitosan

MMW chitosan was produced from shrimp and crab shells as described above, excluding the enzymatic step. LMW chitosan was produced as micropowder by irradiating crustacean chitosan with γ-radiations and subsequent jet-pulverisation. This technology was developed at the Shanghai Institute of Applied Physics, Chinese Academy of Sciences (Wu et al., 2005). Weight averaged Molecular Weight (Mw), degree of polymerisation (DP) and Dispersity (Đ) were determined by Size Exclusion Chromatography (SEC) coupled with Refractive Index (RI) and Multi Angle Laser Light Scattering (MALLS) detector. Degree of acetylation (DA) was determined by ^1^H-NMR spectroscopy Table 4.

MMW had an average molecular weight of about 116 kDa and contained chitosans in a range of about 15 - 600 kDa. RI chromatogram showed only one polymer peak (see Figure 4). Oligomers were not present in this sample.

**Figure 4.**
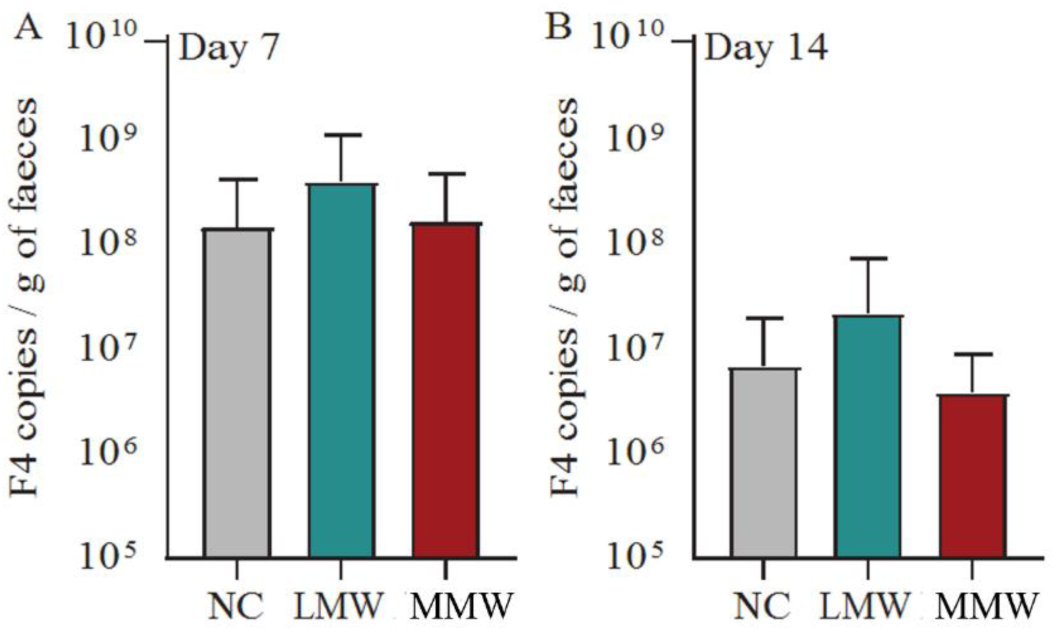
F4-ETEC faecal shedding at day 7 (A) and 14 (B) post-weaning during the second in vivo study. F4 copies per reaction were normalized by grams of faecal material. MMW, medium-molecular weight chitosan; LMW, low-molecular weight chitosan; NC, negative control.

**Figure 5.**
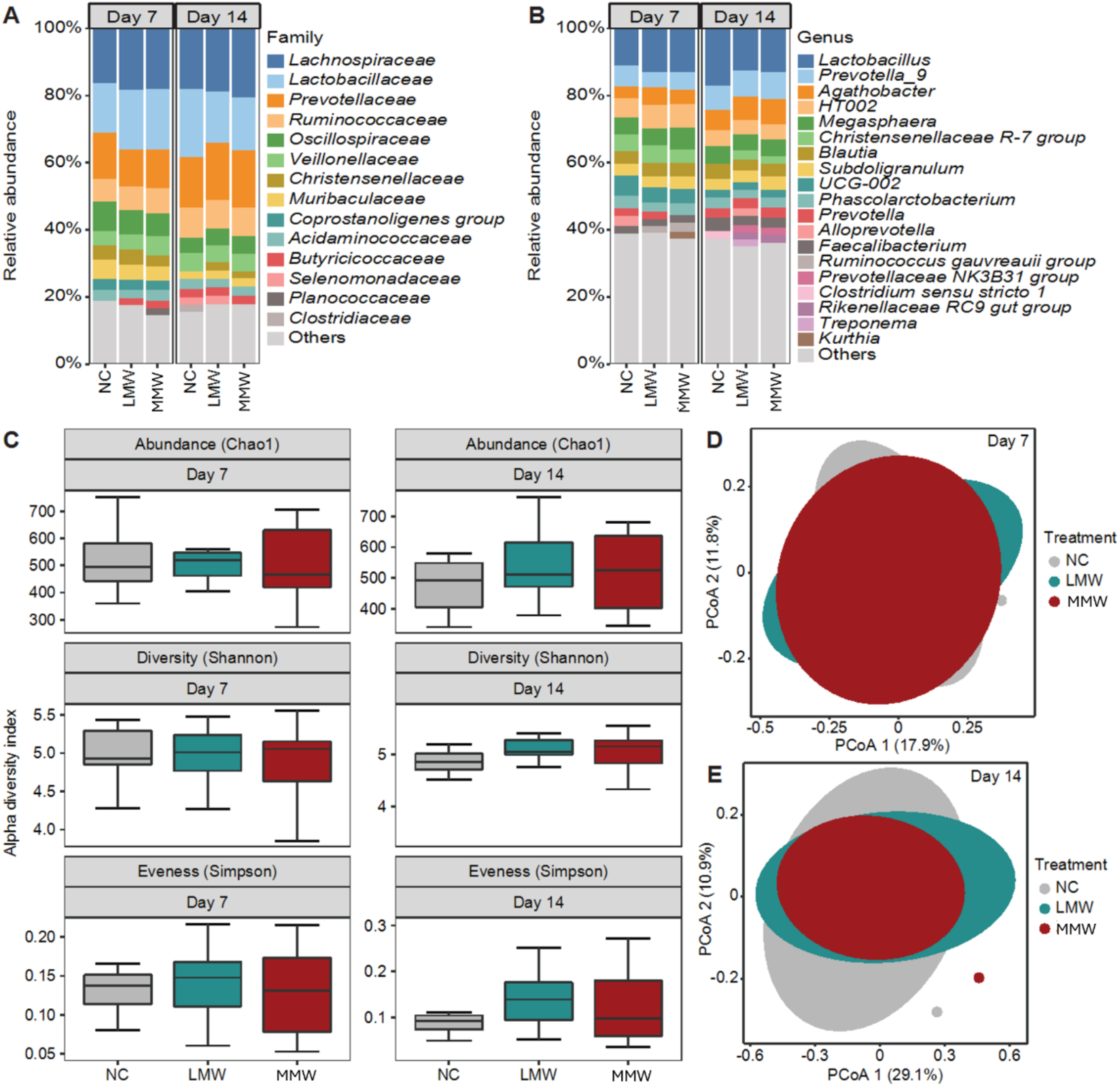
Impact of medium- or low-molecular weight chitosan on the gut microbiota of piglets at day 7 and 14 post-weaning. Relative composition of bacteria at family (A) and genus (B) level. Only taxa with a relative abundance >2% are displayed. Box-plot of α-diversity calculated with Chao1, Shannon, and Simpson indexes (C). Two-dimensional principal coordinates analysis (PCoA) plot based on the BrayCurtis dissimilarity matrix of samples collected at day 7 (D) and 14 (E) post-weaning.

In contrast, the LMW chitosan sample displayed an average molecular weight of only 14.4 kDa and covered a mass range of about 0.3 kDa to 30 kDa. The SEC-RI-MS measurement revealed the presence of chitosan oligomers down to chitosan dimers (chitobiose), in addition to the polymeric portion.

Overall, the molecular weight of the polymeric part of the LMW chitosan was smaller and less dispersed than MMW chitosan,. is confirmed by the molar mass distribution (see Figure 3.c).

##### In vivo zootechnical studies results

#### 2.1.3. 1^st^ trial – COS HCl

The first trial was a dose finding study, where three COS HCl inclusion levels (doses) were tested, 0.025, 0.05 and 0.1%, with comparison to the basal diets (0%). The piglets were followed for 42 days post-weaning.

##### Piglet health

Four piglets died from meningitis between day 0 – 42 post-weaning (2.1% mortality; Supplementary Table S5, with one being euthanized in agreement with pre-defined humane endpoint. Six piglets were medically treated with injectable antibiotics in this period, i.e. 3.1% of the piglets (Supplementary Table S6). Medical interventions consisted of treatments for arthritis (3 out of 8), meningitis (3 out of 8), pneumonia (1 out of 8) and eye infection (1 out of 8).

##### Piglet performance and gut effect

The effect of dietary treatment on pig performance is summarised in Tables 6.

The average BW of the weaned piglets at day 0, 14 and 42 post-weaning was 9.4 ± 0.03 kg, 13.5 ± 0.09 kg and 33.2 ± 0.28 kg. Overall ADG was 571 ± 12.8 g/piglet/day, overall ADFI 812 ± 12.7 g/piglet/day, and overall FCR 1.42 ± 0.015 between day 0-42 post-weaning. The FS was on average 6.22 ± 0.058 for the total experimental period.

No significant effects of dietary treatments on piglet performance were observed, i.e. BW, CV in BW, ADG, ADFI, and FS, in any of the experimental periods, i.e. day 0-14, day 14-42 and day 0-42 post-weaning.

The lowest dose of COS (0.025%) was associated with the lowest values for BW, ADG, and ADFI. Although there were no statistically significant differences between the 0.025% COS group and the negative control, COS 0.025 % treatment showed a negative effect in performance. Supplementation with the high dose of COS increased the FCR between day 0-14 post-weaning compared to the negative control diet (*P*<0.05), with intermediate FCR scores for the low and medium dose groups. The effect was not observed between day 14-42 post-weaning or in the overall experimental period (day 0-42 post-weaning). Results excluding outliers are presented in Supplementary Table S9 and they indicate an increase in FCR between day 0-14 post-weaning as well, but no significant differences were observed using post-hoc pairwise comparisons of least square means with Tukey’s adjustment.

A visible drop in faecal score was observed on day 4 post-weaning, which quickly recovered to the optimal score of 6 by day 7 post-weaning (Figure S2). After three weeks post-weaning, the faecal score increased by 0.5 to 1 point.

Polynomial regression analysis indicated a linear dose-response relationship (P < 0.01) between COS concentration and FCR from day 0–14, with a reduction in feed efficiency with increasing doses of polymer in the diet of weaned piglets. No significant dose-response effects were observed for the other performance parameters and faecal consistency. The results including and excluding outliers are summarised in the Supplementary Tables S15 and S16.

#### 2.1.4. 2nd trial – LMW and MMW test products

The second trial aimed to test efficacy, comparing low molecular weight and medium molecular weight chitosan at 0.01% inclusion, monitored for 42 days.

##### Piglet health

Thirteen piglets were medically treated with injectable antibiotics between day 0-42 post-weaning, i.e. 4.5% of the piglets. An overview of the antibiotic treatments is given in the Supplementary Tables S10 and S11. Medical interventions consisted of treatments for arthritis (6 out of 13), meningitis (4 out of 13), pneumonia (2 out of 13), and general weakness (1 out of 13). The number of piglets treated did not differ between experimental treatments (P = 0.30). Antibiotic treatments across the different disease classes (arthritis, meningitis, pneumonia, and general weakness) did not differ between experimental treatments (P = 0.19). Five piglets died between day 0-42 post-weaning (1.7% mortality) and one piglet was moved to a hospital pen (Supplementary Table S12). Mortality did not differ between experimental treatments (P = 1.00).

##### Piglet performance and gut effect

Average body weight of the weaned piglets was 7.65 ± 0.039 kg at day 0 (average weaning age 29.6 ± 0.08), 11.3 ± 0.09 kg at day 14, and 29.8 ± 0.24 kg at day 42 post-weaning. Overall ADG was 571 ± 12.8 g/piglet, overall ADFI 701 ± 13.8 g/piglet, and overall FCR 1.32 ± 0.013 between day 0-42 post-weaning.

No significant effects of dietary treatments were seen on piglet performance, i.e. BW, CV in BW, ADG, ADFI, FCR, and FS, in any of the experimental periods, i.e. day 0-14, day 14-42 and day 0-42 post-weaning (Table 7). Results excluding outliers (Supplementary Table 14) also indicate no significant effects of low and medium molecular weight chitosan on piglet performance.

**Table 7.**
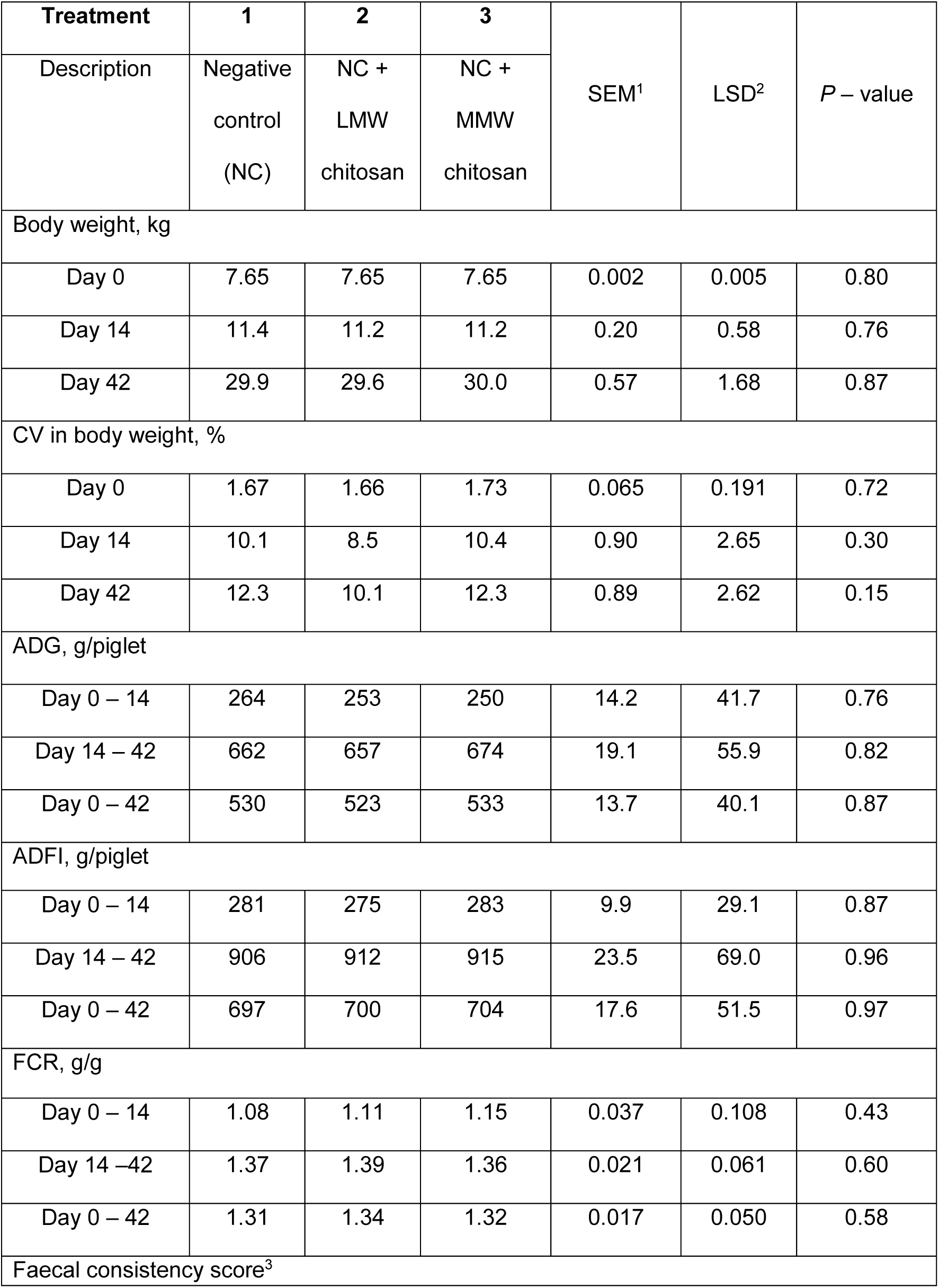
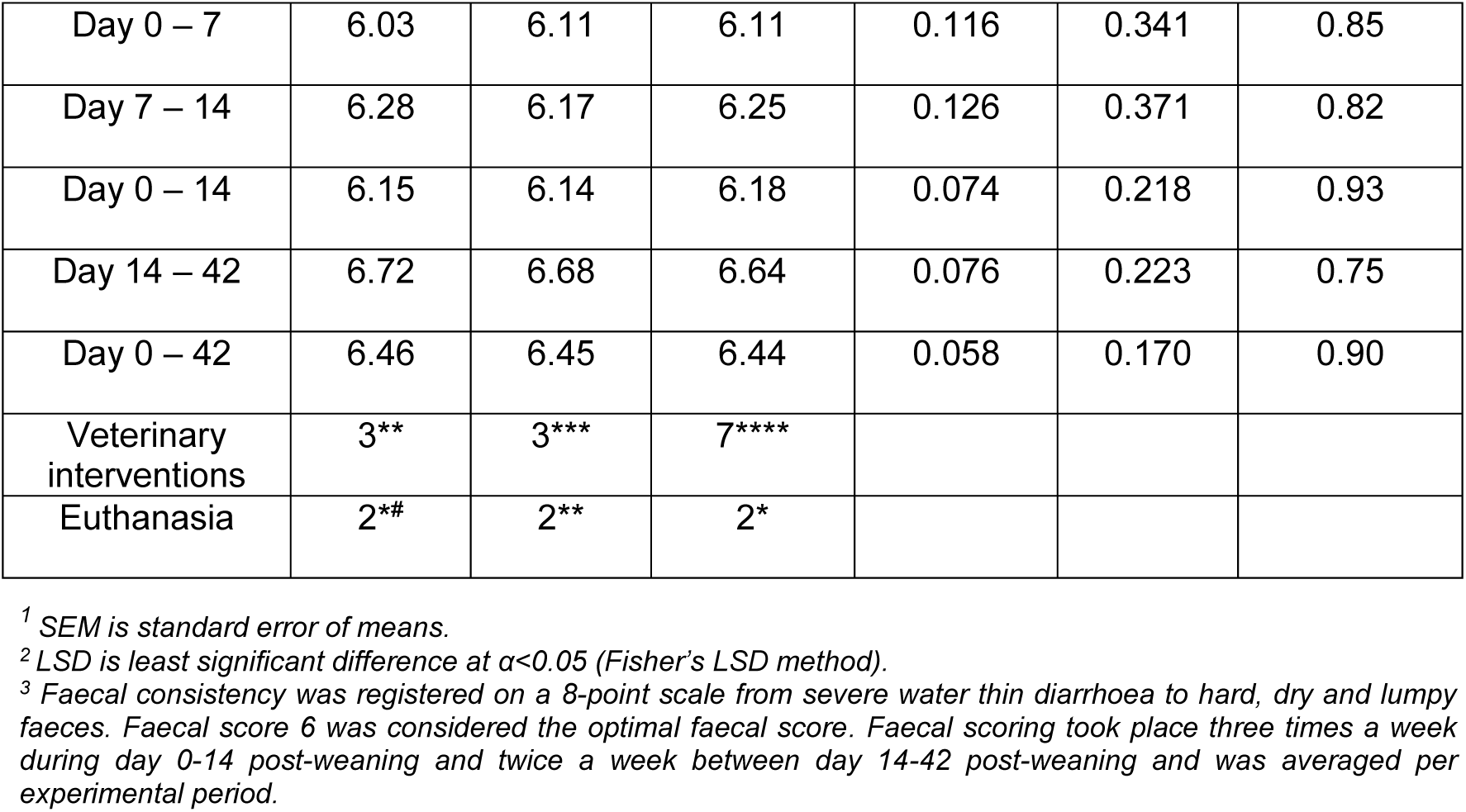
Effect of experimental diet on piglet performance between day 0-42 post-weaning (including outliers), veterinary interventions (**meningitis (2) & pneumonia(1),***arthritis(2) & general weakness (1), ****arthritis (4), meningitis (2) & pneumonia(1)) and mortality (* meningitis (3), # unknown (1),** meningitis (1) and pneumonia (1))

**Table 8.**
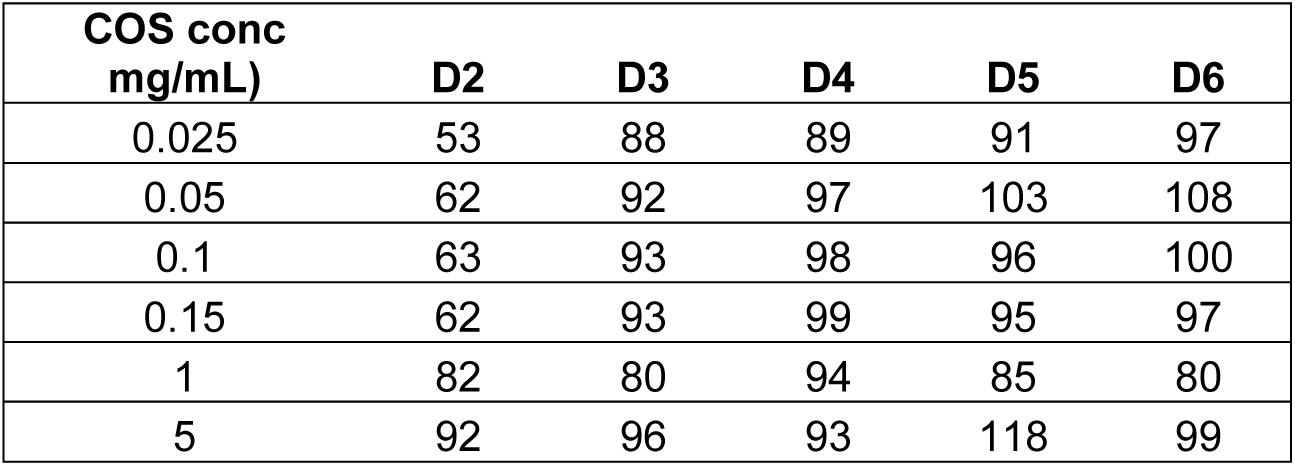
Recovery (%) of COS at different concentrations in the presence of maize (4 mg/ml). D: Glucosamine.

| **COS conc**<br>**mg/mL)** | **D2** | **D3** | **D4** | **D5** | **D6** |
| --- | --- | --- | --- | --- | --- |
| 0.025 | 53 | 88 | 89 | 91 | 97 |
| 0.05 | 62 | 92 | 97 | 103 | 108 |
| 0.1 | 63 | 93 | 98 | 96 | 100 |
| 0.15 | 62 | 93 | 99 | 95 | 97 |
| 1 | 82 | 80 | 94 | 85 | 80 |
| 5 | 92 | 96 | 93 | 118 | 99 |

The faecal consistency scores (FS) (Figure S3) throughout the post-weaning period ranged from 5.7 to 7.2. A one-point drop in the faecal score was observed during the first week post-weaning, which quickly recovered to the optimal score of 6 in the second week. No statistically significant differences in FS were observed between the treatments and negative control groups.

F4-ETEC faecal shedding at day 7 and 14 post-weaning did not significantly differed between groups (Figure 7B and 7C).

Faecal microbiome analysis of samples collected at day 7 and 14 post-weaning showed a similar bacterial composition at family and genus levels (Figure 8A and 8B).

Similarly, no differences in alpha- and beta-diversity were observed between treatments group and control between both at day 7 and 14 post-weaning (Figure 8C-E).

##### Detection and Quantification of chito-oligosaccharide HCl or LMW and MMW chitosan in complex matrices

#### 2.1.5. Chito-oligosaccharide HCl

The matrix effect of maize and feed on the detection and quantification of COS HCl was analysed by SEC-RI-MS. Pre-tests were run using a higher concentration of COS HCl than the actual premix and feed concentration.

The addition of maize (4 mg/mL) to pure COS HCl (1 mg/mL) led to a slight reduction of detectable COS. The further addition of feed (10 mg/mL) to COS HCl (1 mg/mL), which was already mixed with maize (4 mg/ml), has almost no influence.

This is visible by overlying the RI signal of COS HCl (1 mg/mL) alone, with maize and with maize and feed, as shown in Figure 6-a. Maize with a concentration of 4 mg/mL and feed with a concentration of 10 mg/ml have almost no effect in the signal intensity of the COS independent of their one concentration. To further investigate maize effects on COS HCl detection and quantification, different concentration of COS HCl (0.025 mg/mL – 5 mg/mL) were measured alone or in combination with maize (4mg/ml) or maize (4 mg/ml) and feed (10 mg/ml) (Supplementary Figure S4). The recovery rate of these compounds is shown in Table 8. It was evident that to a lower concentration of COS HCl in the sample corresponds a lower recovery rate of the oligomers, especially for the smallest oligomer D2, whose recovery rate dropped from 92% (concentration 5 mg/mL) to 53% (concentration 0.025 mg/mL). When the actual premix and feed samples were analysed (Figure 6-b), the reduction in COS HCl signal was much stronger compared to the pre-tests.

**Figure 6.**
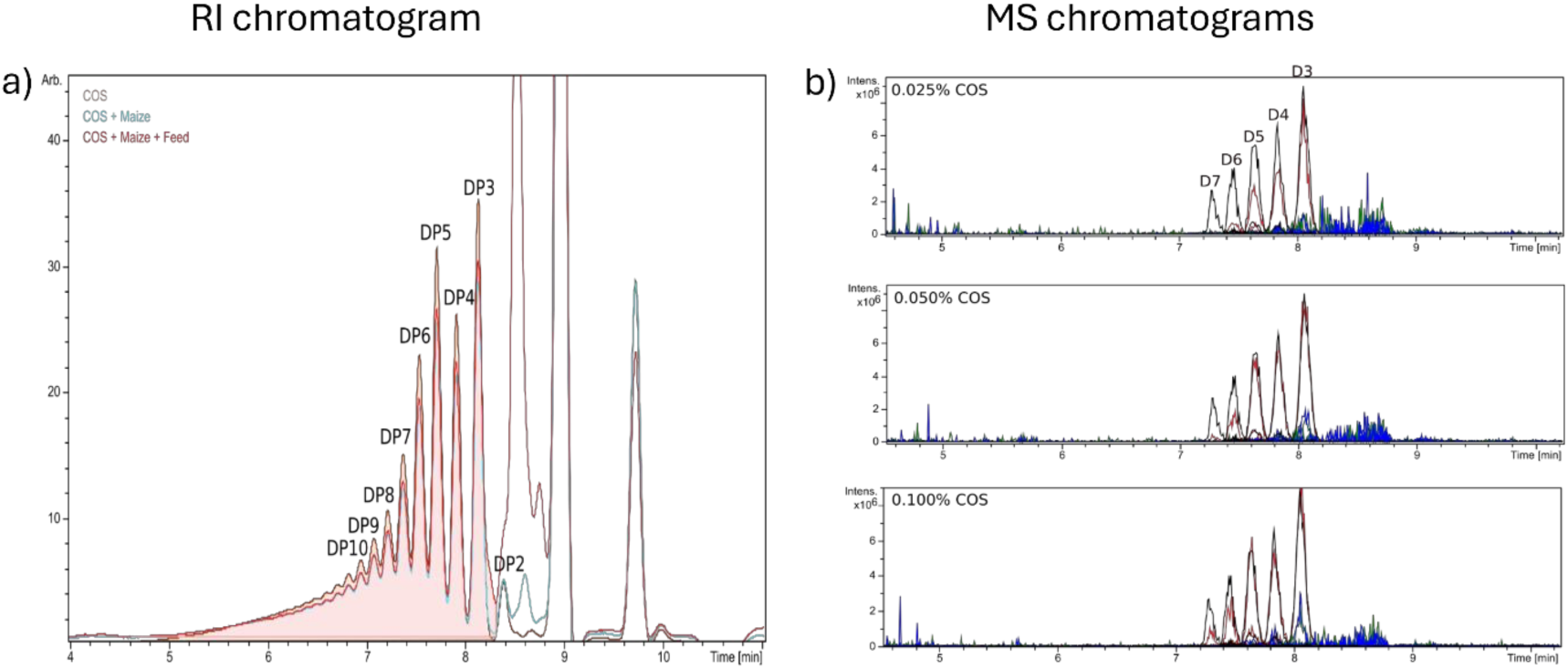
a) RI-Chromatograms showing the matrix effect on COS signals. Three different simulated samples were analysed: COS at 1 mg/ml, COS at 1 mg/ml mixed with 4 mg/ml of maize, and COS mixed with 4 g/ml of maize and 10 mg/ml of feed. Peaks are labelled with the corresponding DP of the COS eluting at each time point. b) Extracted ion chromatograms of COS oligomers (D3 - D7) which were detected in COS alone. (black), Premix (red), Mash (green) or Pellets (blue) samples contain different concentration of COS (0.025% (series B), 0.05% (series C) or 0.1% (series D)). All samples were dissolved in water, filtrated using the 0.2 µm filter, freeze dried and afterwards dissolved in the SEC buffer (150 mM NH4Ac pH 4.2) to have a concentration of 0.1 mg/ml COS in all samples. Then 5 µl of each sample was injected into the SEC-MS-RI system

**Figure 7.**
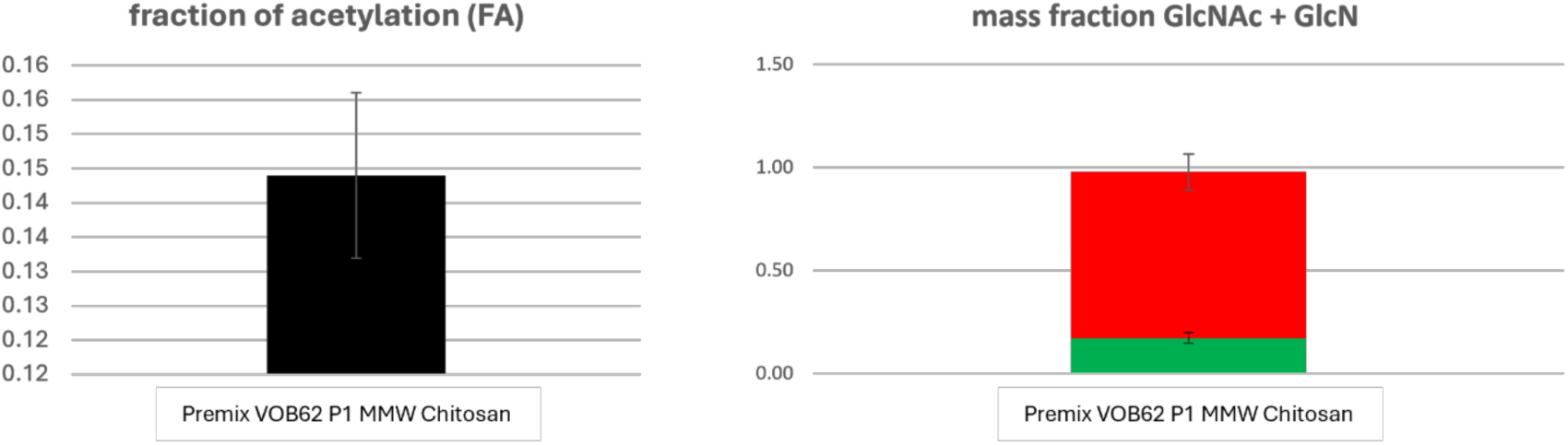
Fraction of acetylation (FA) and mass fraction for MMW chitosan in the premix obtained by enzymatic method/ UPLC-MS. The green bar correpsonds to N-acetyl glucosamine (GlcNAc) content whereas the red bar corresponds to glucosamine (GlcN) content

For the premix sample (red) the loss was not very strong, especially for the very small oligomers like D3, D4 or D5. However, the larger oligomers showed a much higher loss. In the sample with COS, maize and feed, almost no COS was detectable. Only a very small peak for D3 and D4 are visible.

Preliminary analyses (Figure 6-b) showed that, although there is a signal suppression, detection in premix is possible, but not in feed as COS HCl signal almost disappeared completely when mixed in feed.

#### 2.1.6. LMW and MMW chitosan

The SEC-RI-MS method, used for COS HCl, failed to provide satisfactory detection (see Supplementary Figure S9). As an alternative, an enzymatic method coupled to UHPLC-MS (see Materials and methods “Enzymatic method 1” - Urs et al. 2023) was used to try to detect and quantify COS HCl, LMW and MMW chitosan in the premix.

This method worked only for MMW chitosan samples. The results for MMW chitosan show a degree of acetylation (DA) close to 15%, which is consistent with the value found by NMR (15%), and a mass fraction close to 1, consistent with the percentage of inclusion (1%) of MMW chitosan in the premix (Figure 7).

The Urs et al. method was modified to improve low molecular weight chitosan and COS quantification. In particular, the washing steps were skipped, but there was no improvement in the results as the salt concentration was so high that the samples could not be concentrated in the last steps and therefore could not be analysed by HILIC-MS without causing salt precipitation. These salts also interfered with the MS analysis itself. A second method (see Materials and Methods “Enzymatic method 2”) was developed but the recovery rate of the chitosan was between 12 – 160%, making the method unsuitable for this matrix.

## Discussion

The results provide a robust evaluation of the potential for chitosan to improve piglet zootechnical performance, consisting of a scoping review followed by chitosan selection, performance comparisons and chemical analysis. The scoping review provides the first semi-systematic effect size comparison across in vivo studies and a variety of GMP+ grade chitosans, revealing several gaps and opportunities. A prominent gap in analysis of fundamental chemical and physical characteristics was highlighted and led to us performing throughout analyses of a spectrum of chitosans selected for *in vivo* studies. Two large scale zootechnical performance trials indicated that chitosan inclusion in feed did not affect zootechnical parameters. Progress in chemical analysis demonstrated feasibility in quantification of MMW chitosan in feed pre-mix, as required for regulatory compliance.

### Challenges in lead selection and chitosan characterisation

Unlike small molecules with well-defined structures, the absence of a standardised composition in chitosan, with each polymer representing a unique mixture, significantly complicates the selection of the product to test. This heterogeneity is not well recognised and the use of a single CAS number for all adds to confusion. The structure of COSs and chitosans is reflected in the results of our scoping review, which showed a high variability in zootechnical effects, along with uncertainty about the chemical characteristics of the chitosan products used. Dietary inclusion levels ranged between 0.003 and 0.5%, while MW ranged between 1 and 232 kDa. Very negative effect on ADG were only associated with higher MMW chitosans, possibly due to reduction in lipids absorption (Liu et al., 2008). The highest ADG was found when using a LMW chitosan (62 g/d), followed by a MMW chitosan (56 g/d) and a COS (52 g/d), possibly suggesting that all chitosans have potential as feed additives.

For this reason, a criterion to select test products was established, opting for a GMP+ food-grade water-soluble COS (COS HCl), a commercialized LMW chitosan, and a GMP+ food-grade MMW chitosan.

We chose to explore polymer characterization to address the existing gap in data regarding the composition of these polymers. This effort aims to advance our understanding of the patterns that may make chitosan an effective feed additive. Comprehensive analysis by Münster University revealed detailed molecular compositions, including oligomeric and polymeric distributions, far exceeding the limited information found in certificates of analysis (CoAs). As expected, COS HCl was the best characterised having a less diverse composition. While the CoA for COS HCl only reported information about the molecular weight range (<3000) and the degree of deacetylation, our analysis produced detailed information on the oligomeric composition. This included the percentages of oligomers present, which were mainly short chain oligomers.

For LMW and MMW chitosan, the analysis provided details on average molecular weight, polymer composition, molar distribution, and dispersity, well beyond the limited information on deacetylation degree and viscosity in CoAs provided by the manufacturers. These findings indicate that the LMW chitosan used was a mixture of both polymers and oligomers, whereas the MMW chitosan consisted exclusively of polymers.

### Zootechnical study with COS HCl

The relatively low doses of COS HCl used in this study (0.025%, 0.05% and 0.10%) may have been insufficient to elicit the growth-promoting effects observed at higher concentrations. A study by Zhou et al. (date) found that a low dose of COS did not significantly improve growth performance in weanling pigs, whereas a higher dose enhanced nutrient digestibility and reduced diarrhoea incidence. This suggests that a threshold dose could be necessary to achieve beneficial effects in growth and feed efficiency. Also, a low, “sub-therapeutic” dose might disrupt existing microbial equilibrium without providing enough of a positive influence on growth performance (Xu et al., 2018).

### Zootechnical study with Low and Medium molecular weight Chitosans

The dose comparison used in the second in vivo study was influenced by the promising results obtained by Zhang et al. (2020), where 0.01% of LMW chitosan resulted in increased ADG and reduced *E. coli* in the ileal, caecal and colonic digesta of weaned pigs challenged with ETEC. Unfortunately, no significant difference was observed between the negative control and chitosan treated groups for ADFI, BW, CV in BW, ADG, FCR, faecal consistency, and clinical health. Despite extensive reports of the antimicrobial activities of chitosans, faecal microbiome analysis revealed no notable changes in microbial composition. The generally good health status of piglets in the trial may have minimised the potential benefits of chitosan supplementation. An effect may have been masked in this trial which showed an unusually low drop in faecal scores within the facility.

### Challenges in chitosan analytics

The results provide additional information about the structure of the chitosans used in this study and in a wider context demonstrate methodological improvements to the characterisation of chitosans. Further work is needed to achieve robust detection and quantification in challenging premix and feed matrices.

SEC-RI-MS analyses indicate that despite signal suppression, COS HCl can still be detected in premix samples. However, in feed samples, the COS HCl signal nearly disappeared completely. The suppression effect is more pronounced for larger oligomers, and when the method was used to analysed LMW and MMW chitosan did not work at all, supporting the theory that polymers with higher MW bind to components in the maize. To overcome this problem an enzymatic method was employed to quantify the chitosans in the premix.

The first method involved acetylation followed by enzymatic degradation successfully quantified MMW chitosan but struggled with smaller oligomers. Most likely, the small oligomers present in COS HCl and LMW, despite being acetylated, remained water soluble and were removed during the washing steps. In contrast, the larger acetylated chitosans in MMW were not soluble under these experimental conditions. It is important to note that small, fully acetylated chitin oligomers up to DP6 are water-soluble regardless of pH (Ellinor B. Heggset, 2009).

To further improve analysis, a second enzymatic method was developed, involving an initial enzymatic degradation, followed by acetylation and a final enzymatic degradation. This method exploits the high efficiency of the enzymes, and the fact that the resulting compounds from the first enzymatic degradation should be too small to strongly interact with the insoluble particles of the maize. The assay was repeated several times but unfortunately showed inconsistent recovery rates (12 – 160%). There are at least three potential factors that could affect the assay. First, the method involves many steps, leading to potential product loss at each stage. Second, galactose, a small sugar present in maize, can interfere with the enzymatic degradation by reacting with the amino groups of the chitosan leading to inactivity of the enzymes(Hellmann et al., 2025). These side reactions would not occur when acetylation is performed first (see method 1), as the reaction between the amino groups on the chitosan and the acetic anhydride is faster than the reaction between the amino groups of the chitosan and galactose. Third, small components in maize could affect detection by overlapping with the signals of GlcNAc or GlcNAc-d3. The possibility that all these factors contribute simultaneously cannot be excluded.

### Composite opinion on the effects of dietary chitosan on piglet performance?

Variability in chitosan efficacy could be affected by dietary composition and genetics. For instance, maize-based diets rich in negatively charged compounds may reduce chitosan bioavailability by binding to it, as observed during attempts to analyse chitosan in feed. Additionally, genetic differences among pig breeds may influence responses to chitosan (Liufu et al., 2024).

Chemical stability is a further factor, little considered, that could lead to degradation before or during feeding experiments, affecting outcomes. It is difficult to monitor chitosan structure in feed and the gut, bringing uncertainty to the question of stability in practice.

### Future directions and conclusions

While the present study results do not show improvement of zootechnical performances, chitosan remains a promising candidate as a feed additive. Optimisation of dosage, charge and molecular weight properties could be further investigated along with method development needed to both better characterise the chitosans employed and establish reliable detection methods in complex feed matrices, which is needed for regulatory compliance and better understanding of effects and mechanisms. Finally, the results provide encouraging data to support a PWD challenge study, which will be reported separately.

## CRediT authorship contribution statement

**Simona Di Blasio:** Conceptualization, Writing- Original draft preparation, Visualization, Data curation, Writing-Reviewing and Editing. **Anouschka Middelkoop:** Methodology, Investigation, Data curation, Writing-Reviewing and Editing. **Francesc Molist:** Methodology, Writing-Reviewing and Editing, Funding acquisition. **Stefan Cord-Landwehr:** Methodology, Formal analysis, Data curation, Writing-Reviewing and Editing. **Mattia Pirolo:** Formal analysis, Data curation, Writing-Reviewing and Editing. **Alhussein Abdelrahman Elrayah:** Formal analysis (Scoping review), Visualisation (Scoping review), Writing-Reviewing and Editing. **Luca Guardabassi:** Writing-Reviewing and Editing, Funding acquisition, Project administration. **Liam Good:** Conceptualization, Supervision, Writing-Reviewing and Editing, Project administration, Funding acquisition. **Ludovic Pelligand:** Conceptualization, Supervision, Writing-Reviewing and Editing, Visualization (Roadmap), Formal analysis (Scoping review), Funding acquisition.

## Data availability

Data will be made available on request.

