## Supplementary material for "Comparative chemical characterisation of chitosans and their impact on growth, faecal consistency and microbiota composition in weaned piglets": Supplementary figures_Submission.pdf

***Supplementary Figure S1***. *Forest plot*

| Author & Date | Inclusion (%) | DA (%)* | MW (kDa) * |
|---|---|---|---|
| SFR 1 | 0.025 | 9** | 1.1 |
| SFR 1 | 0.05 | 9** | 1.1 |
| SFR 1 | 0.1 | 9** | 1.1 |
| SFR 2 | 0.01 | 8** | 20 to 30 |
| SFR 2 | 0.01 | 10** | 150 to 500 |
| Walsh et al. (2013b)^ | 0.025 | <30** | <1 |
| Wan et al. (2017)^ | 0.0001 | <5** | ≤1 |
| Zhao et al. (2017)^ | 0.016 | ? | 1 |
| Liu et al. (2008)^ | 0.01 | ? | 1.5 |
| Liu et al. (2008)^ | 0.02 | ? | 1.5 |
| Liu et al. (2008)^ | 0.04 | ? | 1.5 |
| Liu et al. (2010)****^ | 0.016 | ? | 1.5 |
| Xiong et al. (2015)^ | 0.003 | ? | 1.5 |
| Yang et al. (2012)^ | 0.02 | ? | 1.5 |
| Yang et al. (2012)^ | 0.04 | ? | 1.5 |
| Yang et al. (2012)^ | 0.06 | ? | 1.5 |
| Yin et al. (2008)^ | 0.025 | ? | 1.5 |
| Zhou et al. (2012) | 0.1 | ? | 1.5 |
| Zhou et al. (2012) | 0.2 | ? | 1.5 |
| Chen et al. (2009)^ | 0.25 | ? | 2 |
| Chen et al. (2009)^ | 0.5 | ? | 2 |
| Yan and Kim (2011)^ | 0.0003 | 75 | 2 |
| Walsh et al. (2013b)^ | 0.025 | <30** | 3 to 5 |
| Walsh et al. (2013b)^ | 0.025 | <30** | 5 to 10 |
| Walsh et al. (2013b)^ | 0.025 | <30** | 10 to 50 |
| Walsh et al. (2013b)^ | 0.025 | <30** | 50 to 100 |
| Egan et al. (2015) | 0.1 | 15** | 114 to 134 |
| Xu et al. (2018)^ | 0.05 | 15 | 232 |
| Han et al. (2007a) | 0.04→0.02→0.02" | ? | low |
| Han et al. (2007a) | 0.1→0.05→0.025" | ? | low |
| Han et al. (2007b) | 0.2→0.1" | ? | low |
| Han et al. (2007b) | 0.4→0.3" | ? | low |
| Duan et al. (2020)^ | 0.003 | ? | low |
| Hu et al. (2018)^ | 0.005 | ? | low |
| Oliveira et al. (2017) | 0.005 | ? | low |
| Oliveira et al. (2017) | 0.01 | ? | low |
| Oliveira et al. (2017) | 0.015 | ? | low |
| Wang et al. (2009) | 0.5 | ? | low |
| Walsh et al. (2013a) | 0.03 | 10** | ? |
| Walsh et al. (2013a) | 0.06 | 10** | ? |
| Walsh et al. (2013a) | 0.12 | 10** | ? |
| Xu et al. (2013)^ | 0.01 | 15** | ? |
| Xu et al. (2013)^ | 0.05 | 15** | ? |
| Xu et al. (2013)^ | 0.1 | 15** | ? |
| Xu et al. (2013)^ | 0.2 | 15** | ? |

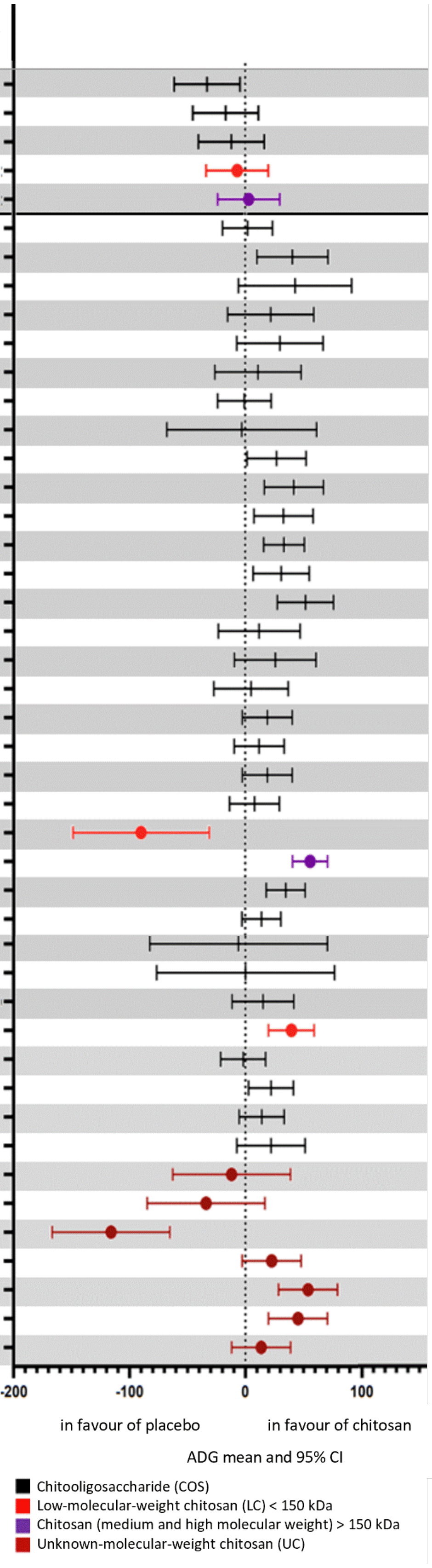

*Table and forest plot summarising the results of Average Daily Growth (ADG in grams per day) of the placebo-controlled studies selected. The point and whisker for each line represent the point estimate of the difference in ADG between treatment and placebo and the bounds of the 95% confidence interval of the difference, respectively.*

***Legend***

** Abbreviations: DA = degree of acetylation; MW = molecular weight- expressed as an average, range or low/high; kDa = kilodaltons. SFR 1 (COS) and SFR 2 (MMW and LWM) are the studies reported in this manuscript. For sources other than 'SFR1 and SFR2', the molecular weight was sorted from lowest to highest to help identify a pattern visually. For MW ranges, sorting referred to the highest number in the range to sort the table. MW marked as 'low' were put at the end to more easily identify any patterns between MW values referring to a specific number.*
*** All these values for degree of acetylation have been converted from degree of deacetylation by subtracting them from 100%.*
**** Figures for this source were only for pigs in the pre-challenge phase (i.e., before being challenged with E. coli) and excluded the post-challenge phase of the study. This allowed better comparison with other studies that did not involve challenging pigs with E. coli.*
*"Arrows refer to studies where different inclusions were given to pigs at different stages of their life cycle. The following are the sources and the respective life cycles they focused on: Han et al. 2007a (grower → early finisher → late finisher), Han et al. 2007b (starter → grower).*
*^ These studies had durations that were lower than 42 days. Studies lower than 42 days in piglet zootechnical studies are typically associated with performance results that are of lower confidence than studies equal to or above this threshold (EFSA FEEDAP Panel et al., 2024).*

***Supplementary Figure S2.*** *COS HCl HNMR – The signal labelled HD corresponds to the solvent residual peak. The H3-H6 region represents the protons attached to the carbon atoms of the sugar ring. The H2 signal corresponds to the proton at the C2 position, which is attached to a nitrogen atom. Finally, CH3 represents the methyl group of the N-acetyl moiety.*

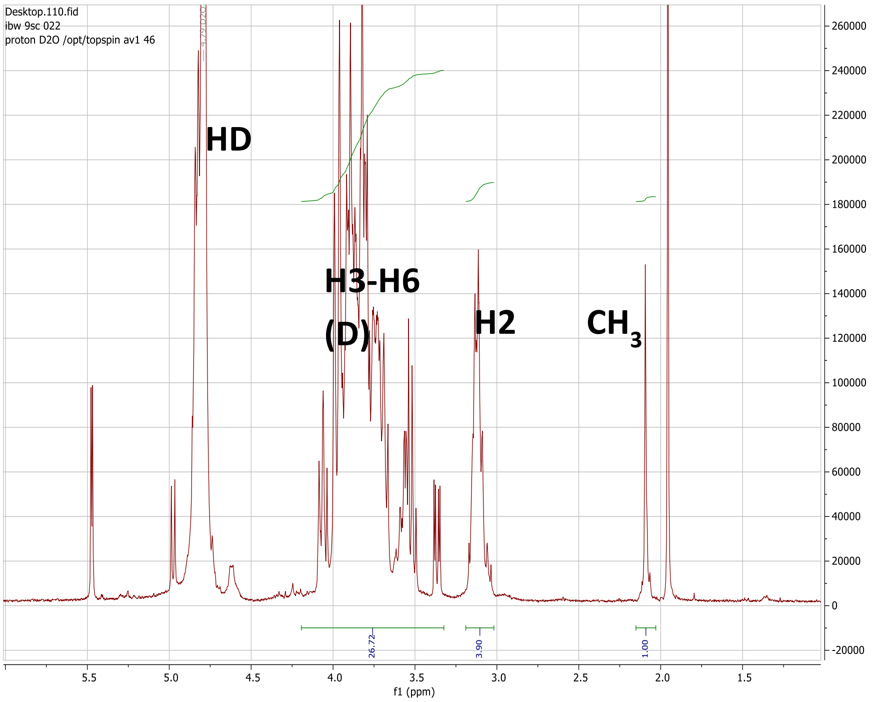


| COS HCl | |
|---|---|
| *Mn [g/mol] | 678 |
| Mw [g/mol] | 824 |
| DP | 4.99 |
| Dispersity | 1.22 |
| DA | 9.5 |

**Abbreviations: Mn is Number Average Molecular Weight, Mw is Weight Average Molecular Weight, DP is degree of polymerisation and DA is the degree acetylation, derived from HNMR.*

***Supplementary Figure S3.*** *Semi-quantitative composition of the COS HCl sample. The percentages were calculated based on the intensities of the different COS in the MS and appear clockwise.*

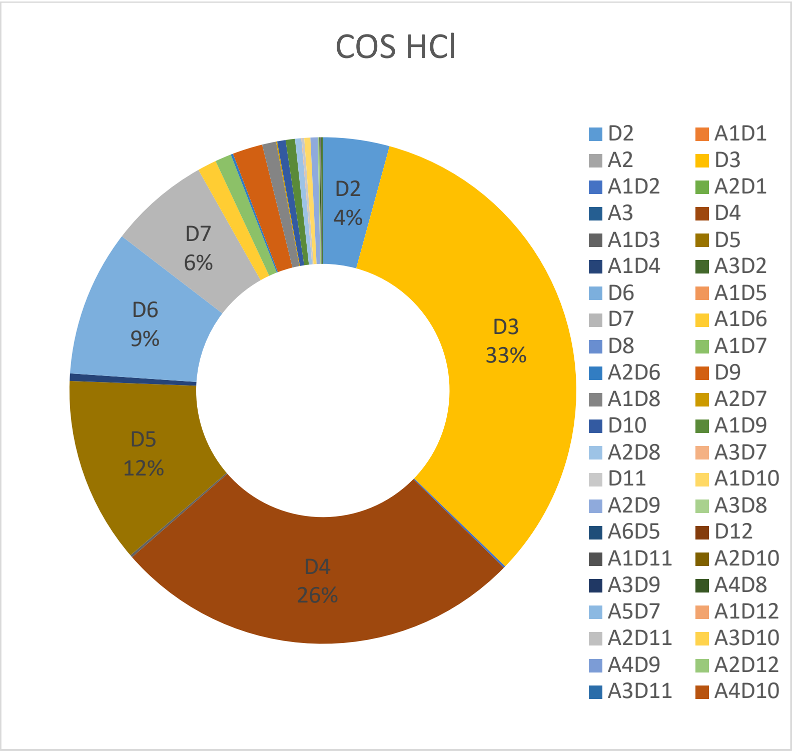

***Supplementary Figure S4** – Faecal consistency score of pigs after weaning in the first in vivo study (chitooligosaccharide).*

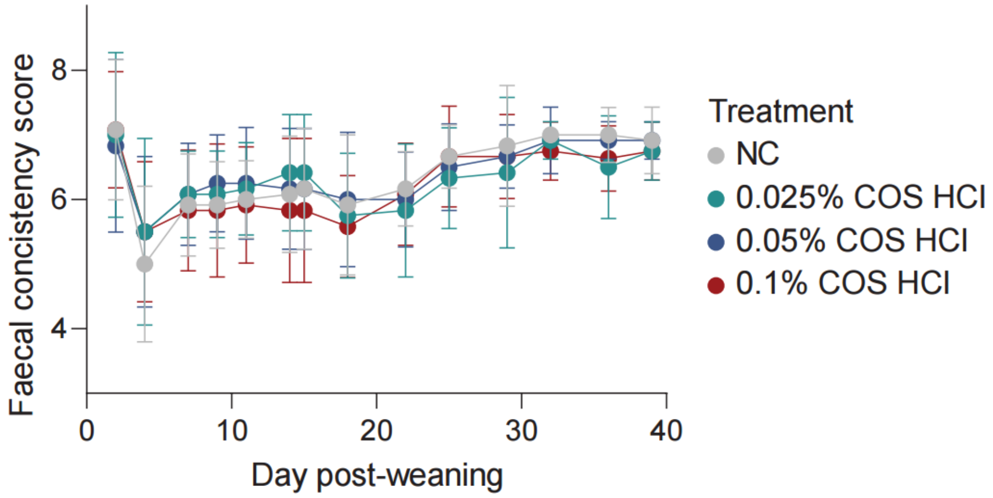


***Supplementary Figure S5** – Faecal consistency score of pigs after weaning in the second in vivo study (Low and medium molecular weight chitosans).*

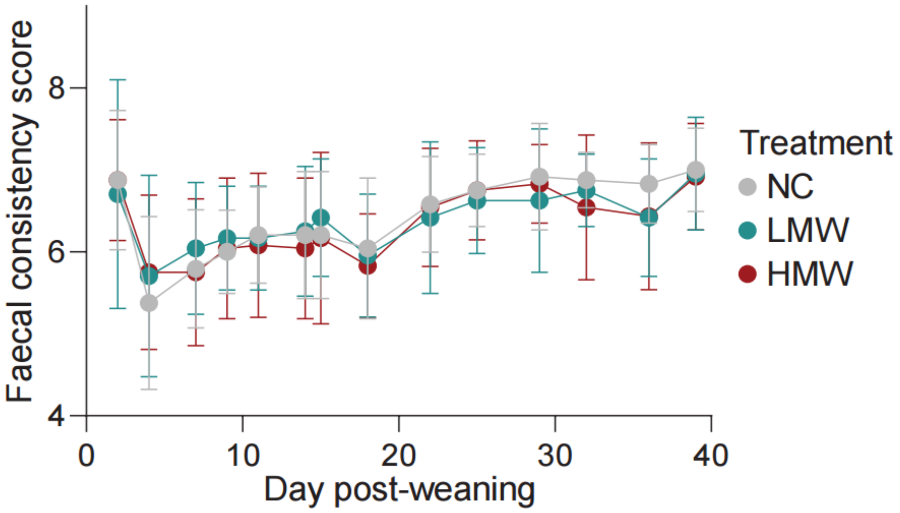

***Supplementary Figure S4*** *– Matrix effect: Intensity of D2 -D4 in COS HCl (0.025 mg/ml – 5 mg/ml) alone or in combination with maize (4mg/ml) or maize (4 mg/ml) and feed (10 mg/ml) in the MS using the SEC-RI-MS approach. The x-axis is the concentration of COS HCl in the sample.*
*The presence of maize and feed leads to slight signal suppression for dimers (D2), signal enhancement for trimers (D3), and negligible interference for tetramers (D4).*

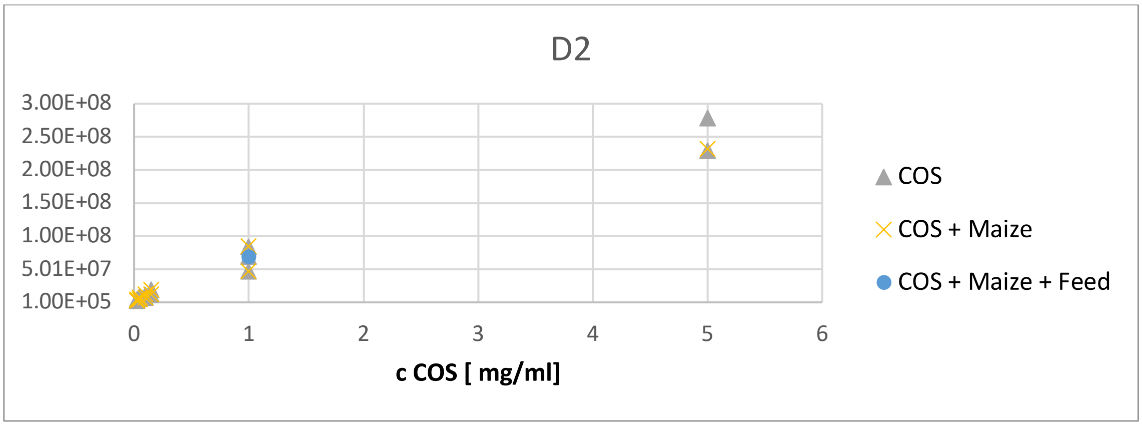


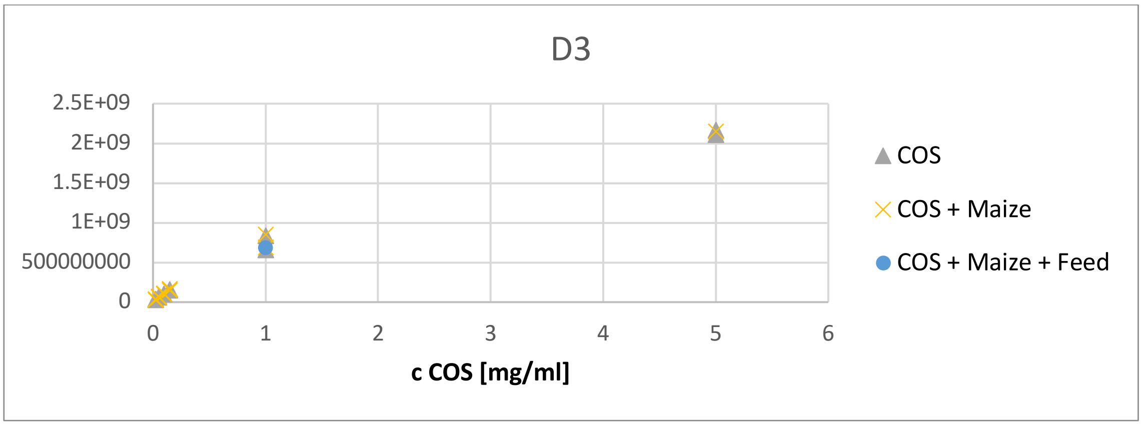


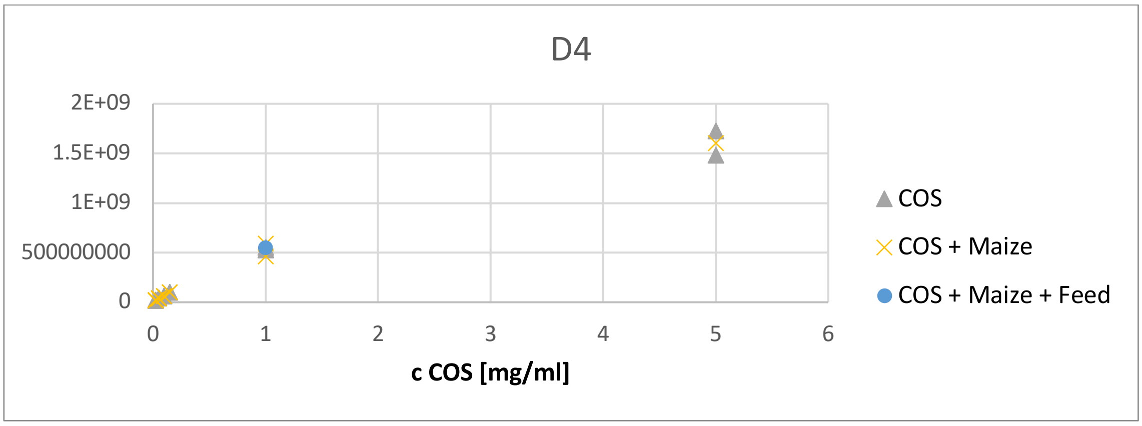

***Supplementary Figure S5*** *– Matrix effect: Intensity of D5 -D6 in COS HCl alone or in combination with maize (4mg/ml) or maize (4 mg/ml) and feed (8 mg/ml) in the MS using the SEC-RI-MS approach. The x-axis is the concentration of COS HCl in the sample.*
*For larger oligomers (D5–D6), the signal intensity of the maize and feed mixture is lower than the maize-only sample, most notably for the pentamer (D5), demonstrating significant signal suppression caused by the feed matrix.*

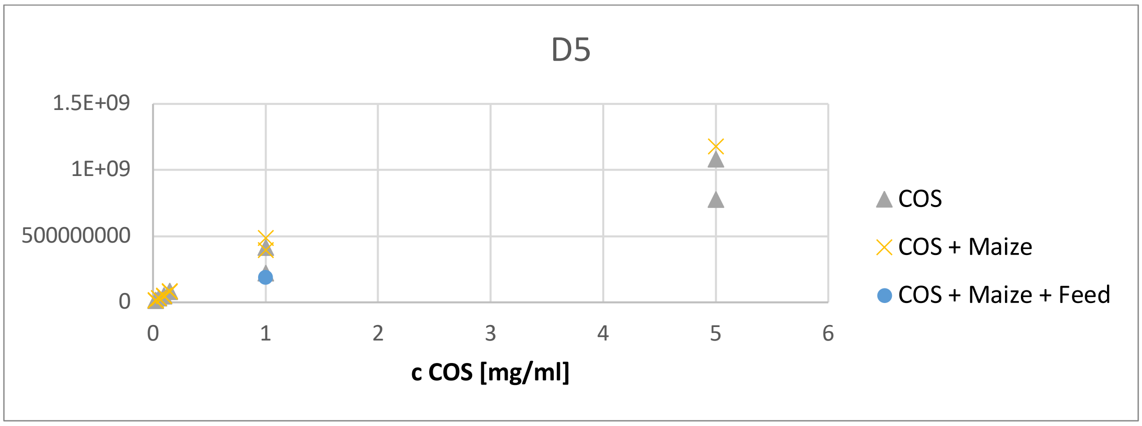


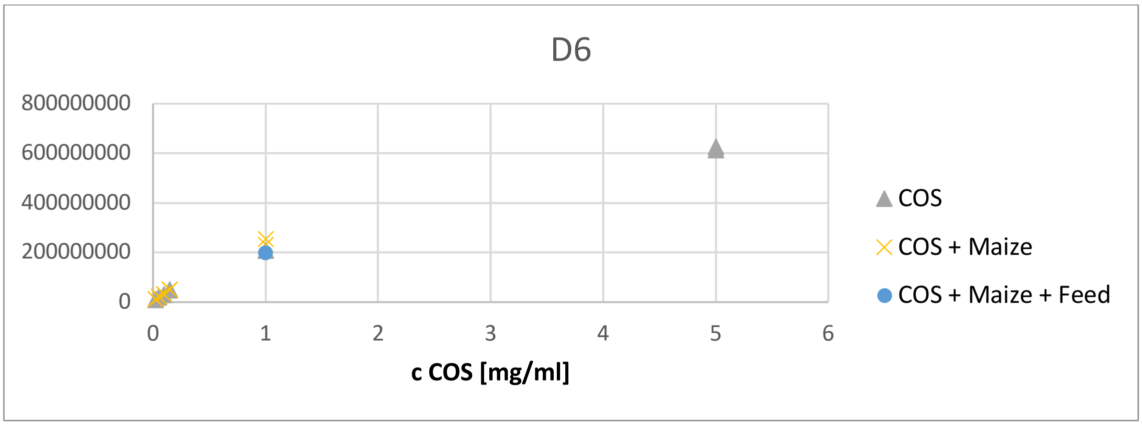

***Supplementary Figure S6*** *– All samples (actual Premix, Mesh, and Pellets) were dissolved in water and stirred. They were then centrifuged, and the supernatant was filtered using a 0.22 µm filter to remove particles. The filtrate was freeze-dried and subsequently dissolved in water to achieve a COS concentration of 0.1 mg/ml in all samples. These filtered and concentrated samples were then used for the SEC-MS-RI measurement. COS was spiked in blank matrix at 0.1 mg/mL for comparison.*
*This process was designed to ensure that COS peaks remained consistent in all measurements, except for the samples without COS (5001 and 6001). However, as observed in the overlay figure, COS was still visible in the Premix sample (A), though the peak intensities decreased as the COS to sample ratio was reduced. In contrast, COS was almost undetectable in Mesh (B) and Pellets (C).*

A. Premix

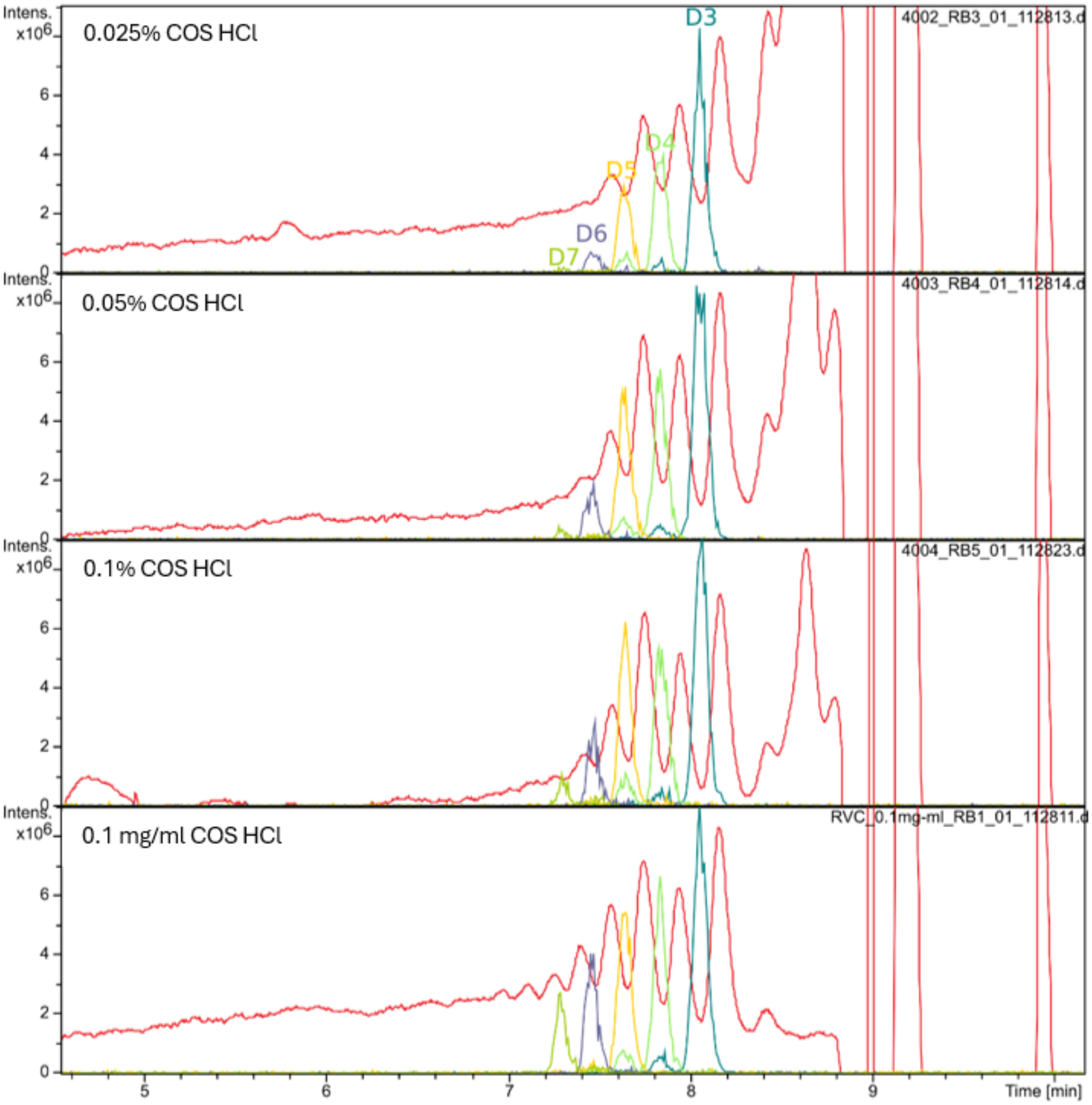


red: RI Chromatogram

B. Mash

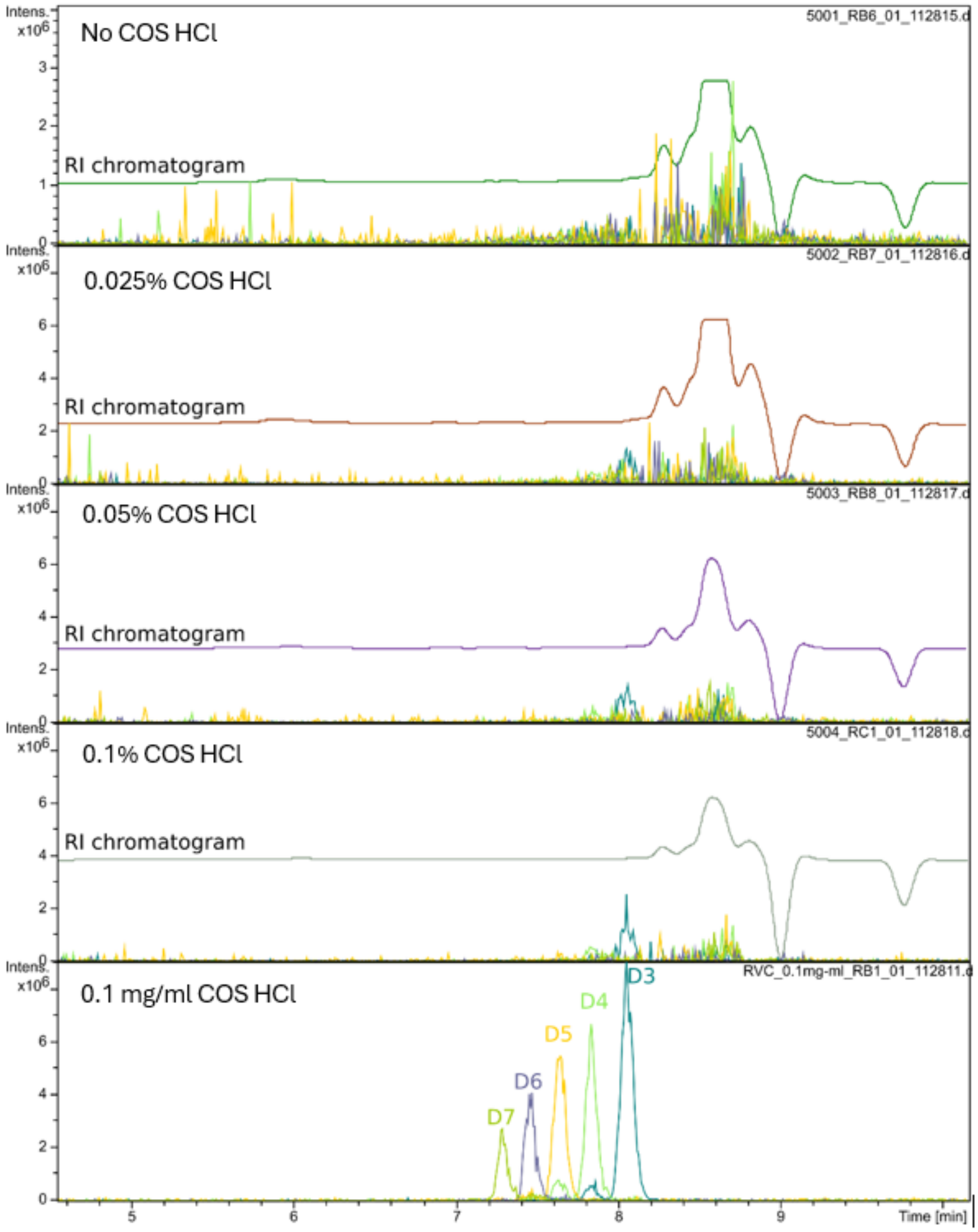

No COS HCl
RI chromatogram
5001_RB6_01_112815.d
0.025% COS HCl
RI chromatogram
5002_RB7_01_112816.d
0.05% COS HCl
RI chromatogram
5003_RB8_01_112817.d
0.1% COS HCl
RI chromatogram
5004_RC1_01_112818.d
0.1 mg/ml COS HCl
RVC_0.1mg-ml_RB1_01_112811.d
D3
D4
D5
D6
D7
Intens. x10^6
Time [min]

C. Pellets

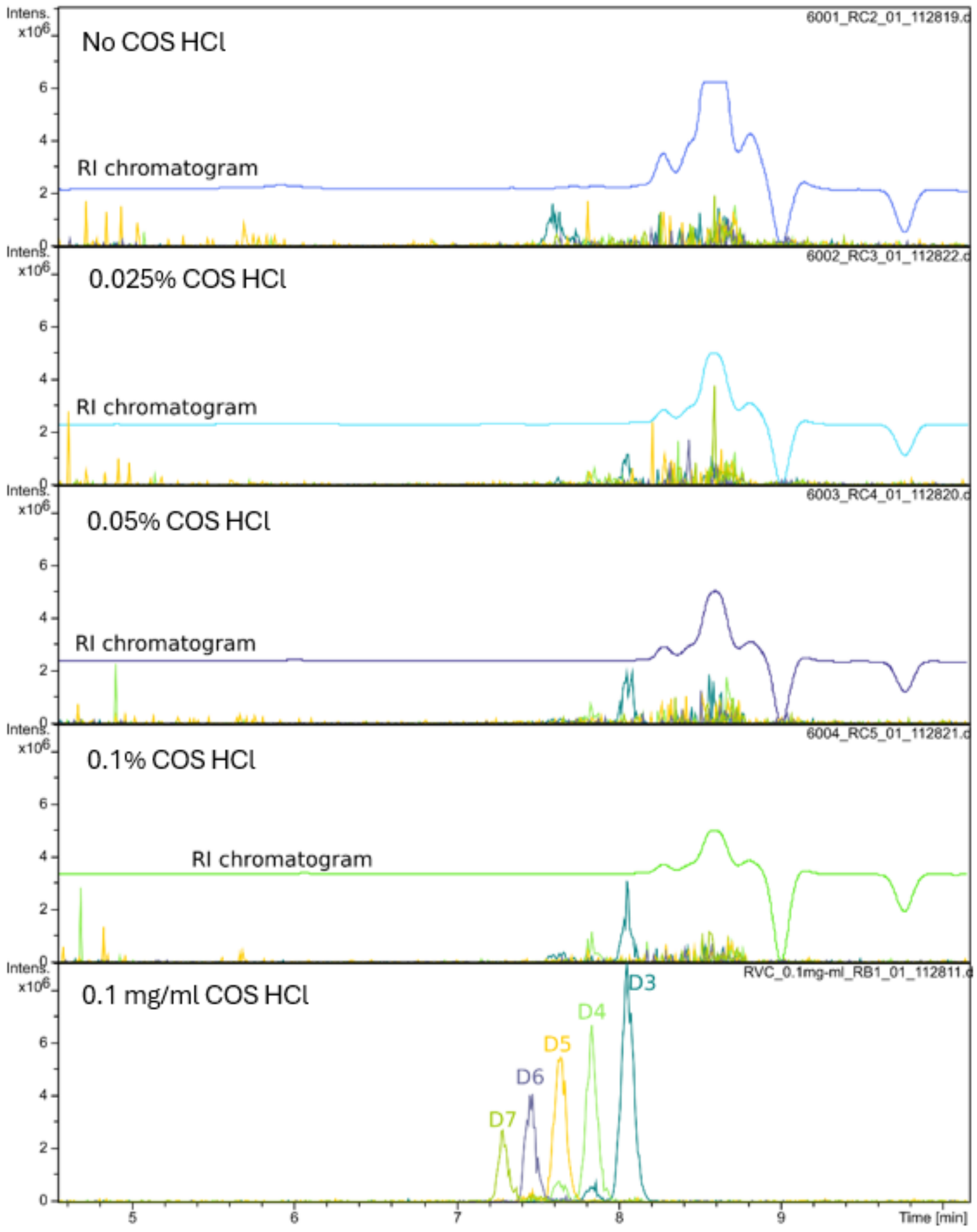

Intens. x10^6
6001_RC2_01_112819.d
No COS HCl
RI chromatogram
6002_RC3_01_112822.d
0.025% COS HCl
RI chromatogram
6003_RC4_01_112820.d
0.05% COS HCl
RI chromatogram
6004_RC5_01_112821.d
0.1% COS HCl
RI chromatogram
RVC_0.1mg-ml_RB1_01_112811.d
0.1 mg/ml COS HCl
D3
D4
D5
D6
D7
0
2
4
6
5
6
7
8
9
Time [min]

*Supplementary Figure S7. LMW HNMR. The signal labelled HD corresponds to the solvent residual peak. The H3-H6 region represents the protons attached to the carbon atoms of the sugar ring. The H2 signal corresponds to the proton at the C2 position, which is attached to a nitrogen atom. Finally, CH3 represents the methyl group of the N-acetyl moiety.*

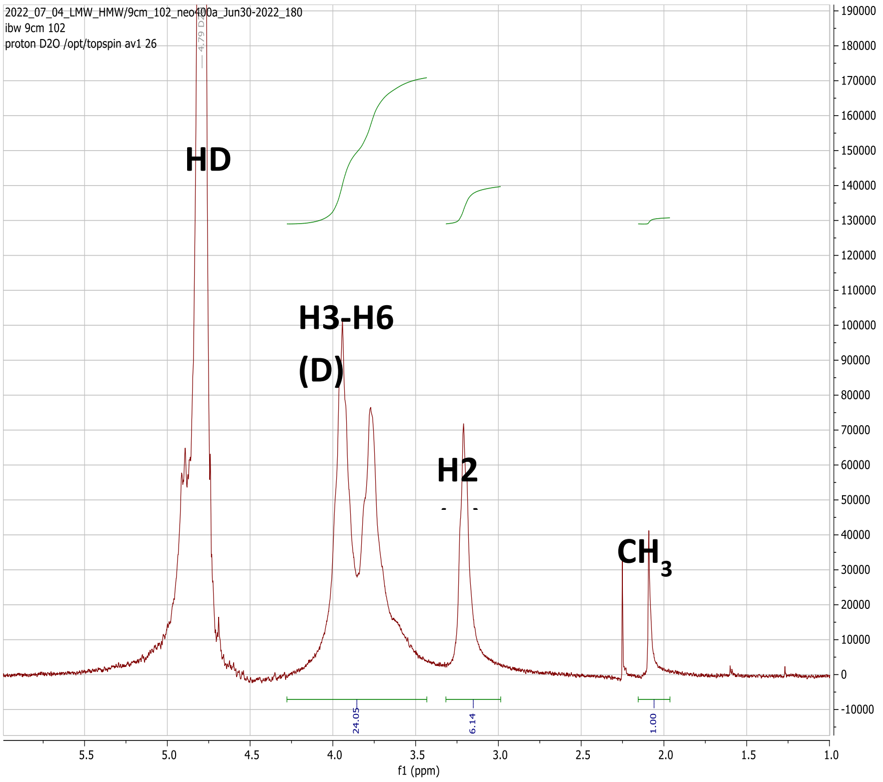


***Supplementary Figure S8*** *- MMW HNMR*

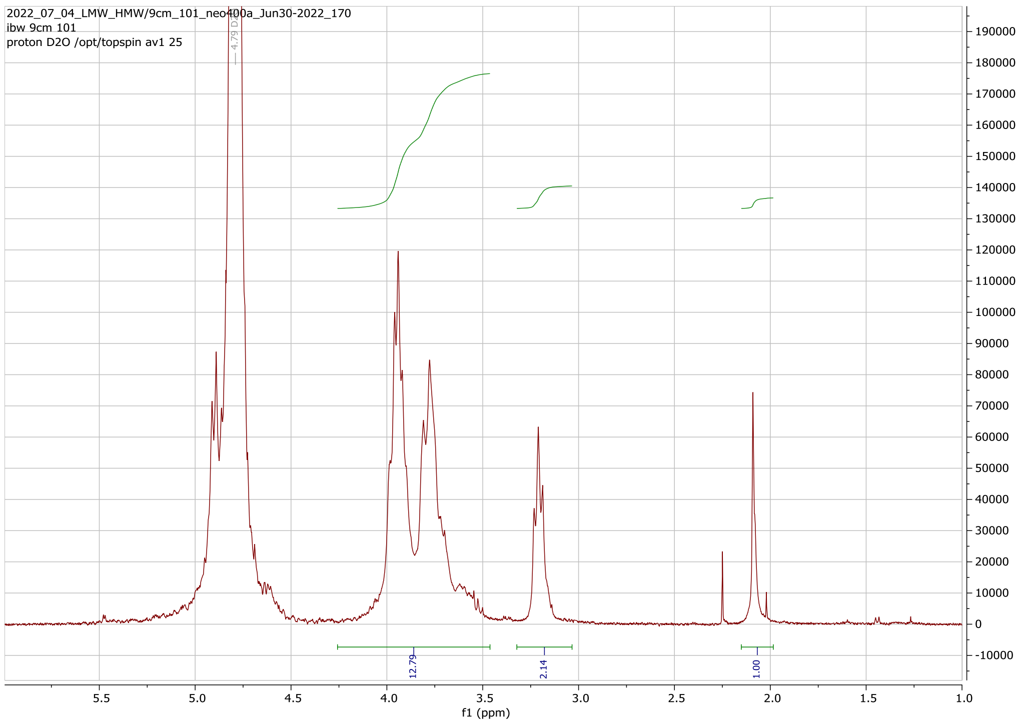

***Supplementary Figure S9** – Mw (Weight Average Molecular Weight), Mn (Number Average Molecular Weight), and dispersity values for Medium and Low molecular weight (LMW and MMW) chitosan, calculated from data obtained using the HPSEC-RI-MALLS system*

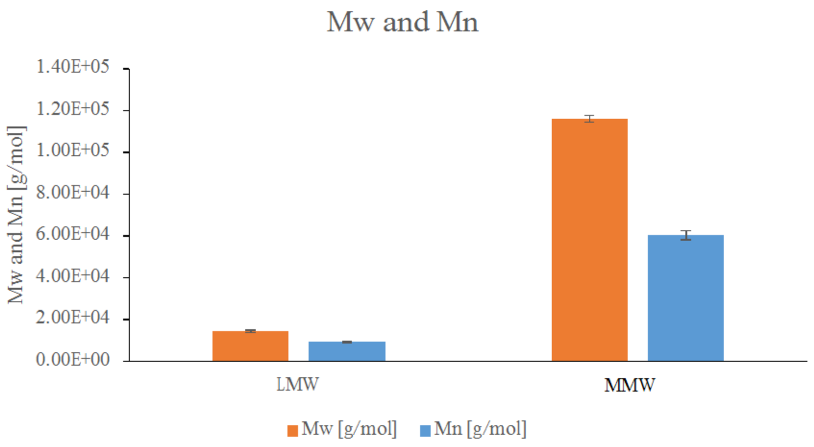


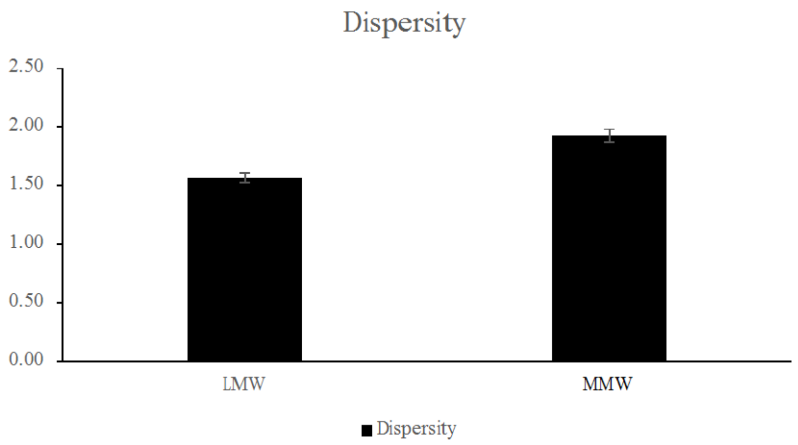

***Supplementary Figure S10*** *- The sample preparation resulted in a theoretical chitosan concentration of 10 mg/ml for the 50x and 40 mg/ml for the 200x. HPSEC-RI-MALLS system was used for the analyses. Usually, the chitosan polymer should be visible at concentration of 10 mg/ml or 40 mg/ml, but here are no peaks which belong to the chitosan polymers visible in the RI-signal. The 3 matrices are Premix (A) , mash (B) and pellet (C)?*

A. Premix

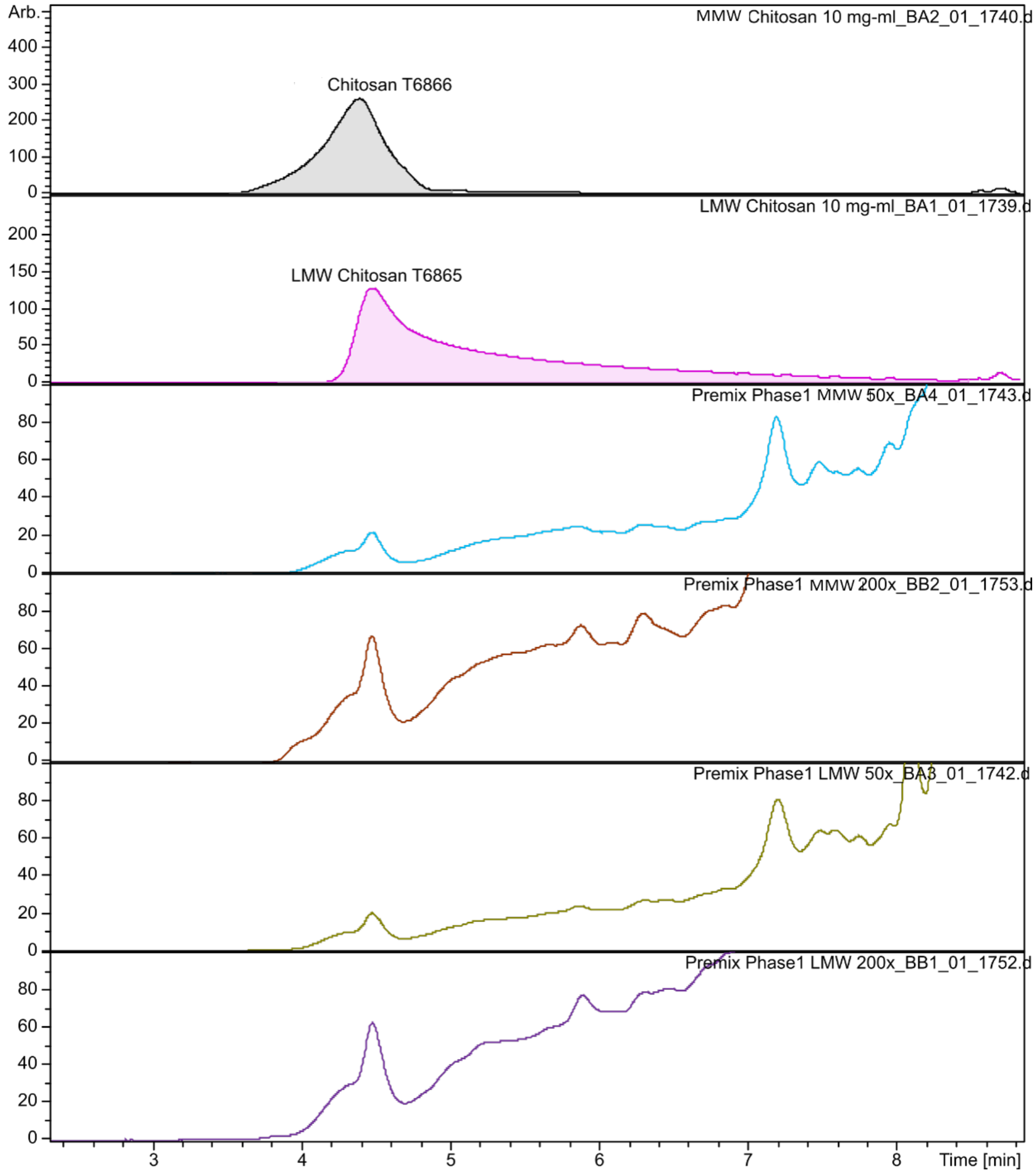

B. Mash

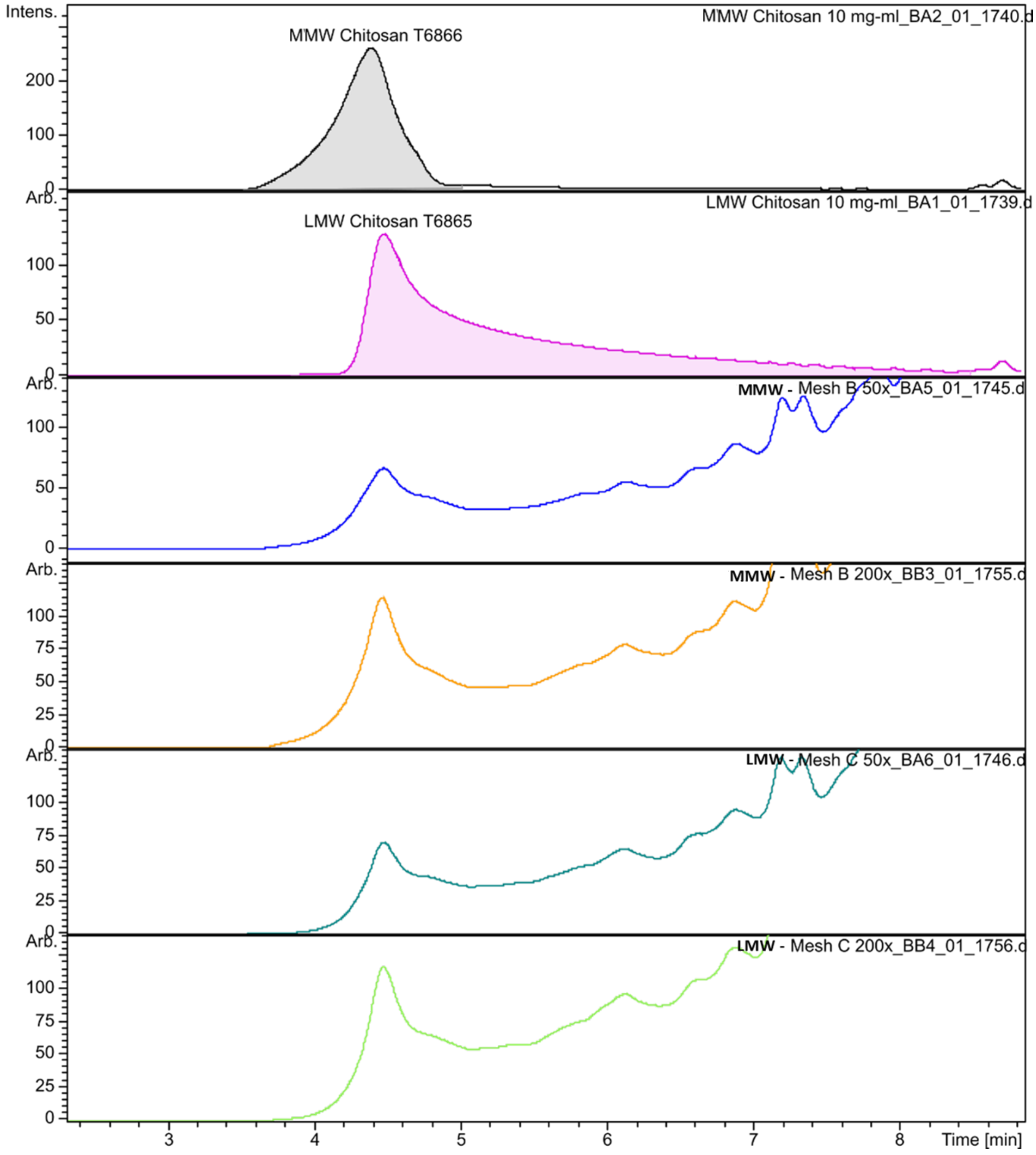

Intens.
MMW Chitosan 10 mg-ml_BA2_01_1740.d
MMW Chitosan T6866
200
100
0
Arb.
LMW Chitosan 10 mg-ml_BA1_01_1739.d
LMW Chitosan T6865
100
50
0
Arb.
MMW - Mesh B 50x_BA5_01_1745.d
100
50
0
Arb.
MMW - Mesh B 200x_BB3_01_1755.d
100
75
50
25
0
Arb.
LMW - Mesh C 50x_BA6_01_1746.d
100
75
50
25
0
Arb.
LMW - Mesh C 200x_BB4_01_1756.d
100
75
50
25
0
3
4
5
6
7
8
Time [min]

C. Pellet

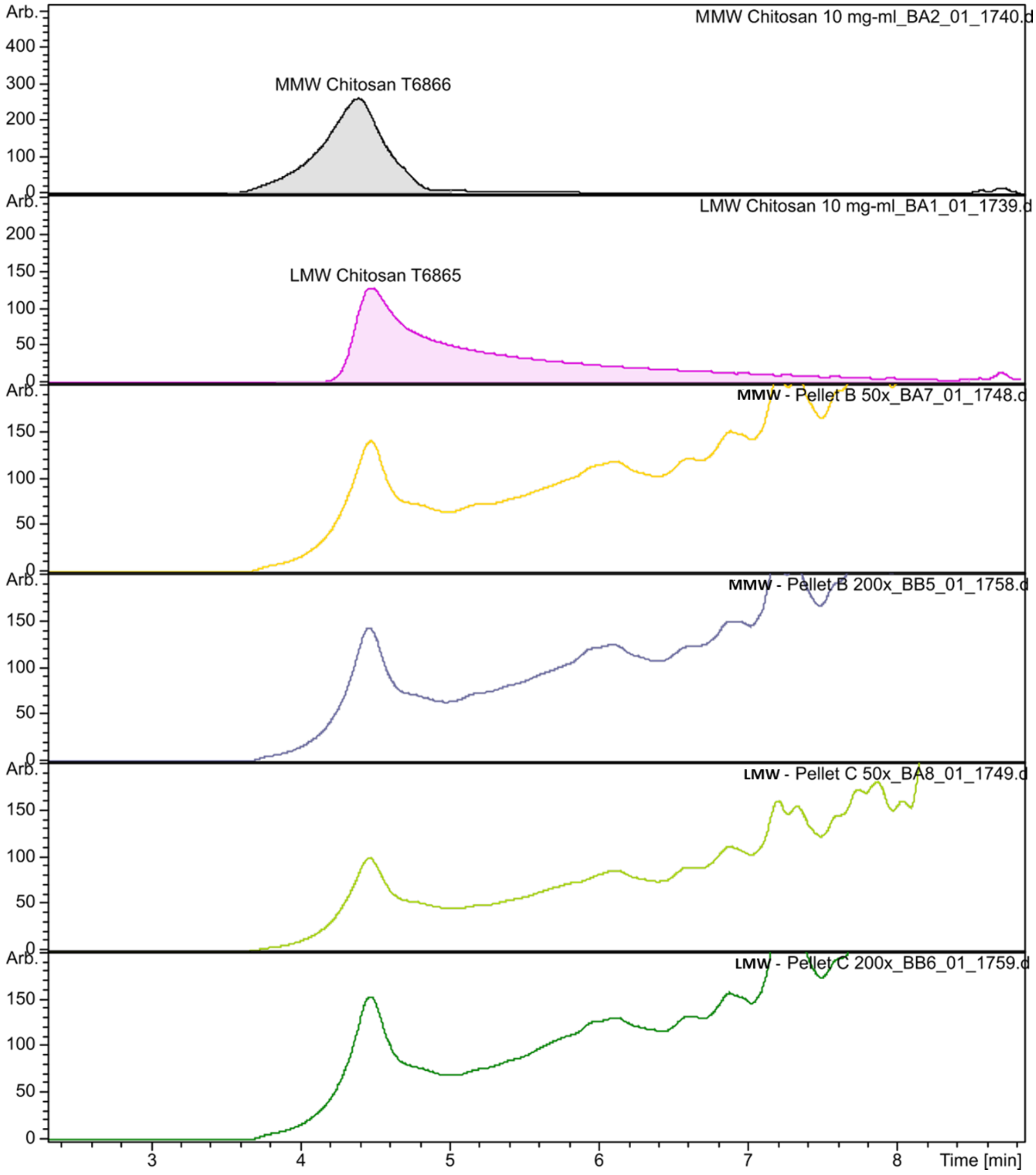

MMW Chitosan 10 mg-ml_BA2_01_1740.d
MMW Chitosan T6866
LMW Chitosan 10 mg-ml_BA1_01_1739.d
LMW Chitosan T6865
MMW - Pellet B 50x_BA7_01_1748.d
MMW - Pellet B 200x_BB5_01_1758.d
LMW - Pellet C 50x_BA8_01_1749.d
LMW - Pellet C 200x_BB6_01_1759.d
Arb.
Time [min]
