## Supplementary material for "Comparative chemical characterisation of chitosans and their impact on growth, faecal consistency and microbiota composition in weaned piglets": Supplementary methods_Submission.pdf

**Method Supplements**

M1: *Proton Nuclear Magnetic Resonance ($^1$H-NMR): all chitosans*

The degree of acetylation (DA) was determined using proton NMR ($^1$H-NMR). Spectra were recorded using a 400 MHz spectrometer (Bruker NEO 400, Germany) after sample preparation according to Kurmiska et al. (2009). Approximately 10 mg of the chitosan solutions were dissolved in $D_2O$ (1 ml 99.9% $D_2O$ and 2 µl DCl), and subsequently lyophilized. The procedure was repeated twice to replace as much hydrogen with deuterium as possible to minimize the $H_2O$ signal in the recorded spectrum. The determination of the DA by $^1$H-NMR measurements was evaluated according to Hirai et al. (1991) and Lavertu et al. (2003). To adjust the chemical shifts, we used the deuterium oxide signal at 4.79 ppm.

M2: *Size exclusion chromatography, refractive index detection, mass spectroscopy (SEC-RI-MS) – all chitosans*

The setup enables quantitative detection of large oligomers and polymers - undetectable by MS - using the RI signal. It is the preferred approach for characterizing the COS spectrum and was used to determine the relative abundance of each oligomer, primarily de-acetylated (degree of deacetylation > 90%), based on the RI detector signal.
Sample analysis was performed according to Hellmann et al. 2024. The COS and chitosan were dissolved in 150 mM ammonium acetate buffer pH 4.2 to a final concentration of 1 mg/ml and 5 mg/ml respectively. The samples were afterwards filtrated using a 0.2 µm modified nylon filter (VWR, Randon, DE, USA). 5 µl of the sample were injected into the SEC-RI-MS system using an Acquity UPLC Protein BEH SEC column (125Å; 1.7µm; 4.6 mm x 300 mM; Waters, Milford, MA, USA), a flow rate of 0.4 ml/min, a column oven temperature of 40°C and a total run time of 14 minutes. The RI chromatogram was recorded with an ERC RefractoMax 524 (Thermo Fisher Scientific) at 10 Hz and 40 °C with recorder and integrator ranges of 512 µRIU and 125 µRIU/V, respectively.

M3: *Hydrophilic interaction chromatography, mass spectroscopy (HILIC-MS), COS HCl only*

The analysis of the chitosan samples by HILIC-MS was performed according to Hamer et al. 2015. The sample was dissolved in water to a final concentration of 1 mg/ml. Afterwards a 3 kDa PES Filter (VWR, Radnor, DE, USA) was used to remove all larger particles and 1-2 µl of the solution were injected into the system. The separation of the different oligos was achieved by using an Acquity UPLC BEH Amide column (2.1 mm x 150 mm; 1.7 µm particle size; Waters, Milford, MA, USA). For the elution, a gradient system consisting of two solvents was used: solvent A (80:20 ACN:H2O) and solvent B (20:80 ACN:H2O), both containing 10 mM ammonium formate and 0.1% (v/v) formic acid was used. From minute 0 to minute 3.0 isocratic 100% (v/v) A, linear from 0% to 85% (v/v) B from minute 3 to minute 23; isocratic 85% B (v/v) from minute 23.0 to minute 28.0. The column equilibration was done as followed: minute 28.0 to minute 29.0 linear from 85% (v/v) B to 100% (v/v) A, followed by isocratic 100% (v/v) A from minute 29 to minute 32. The column oven temperature was set to 35°C, the flow rate to 0.4 ml/min and the MS (amaZon Speed; Bruker, Germany) was operating in enhanced resolution and positive mode over a scan range from m/z 50 – 2000.

M4: *High-performance size-exclusion chromatography (HP-SEC) coupled to multiangle laser-light scattering (MALLS) and refractive index (RI) detector) – LMW and MMW chitosan*

The weight-average molecular weight (Mw), the number-average molecular weight (Mn) and the dispersity index (Đ) of the chitosan polymers were measured by gel permeation chromatography (TSKgel® columns from Tosoh: PWXL-CP-guard column + G6000 PWXL-CP +G5000 PWXL-CP) coupled with a multi-angle-laser-light-scattering (PSS SLD 7000 MALLS®) equipped with a 5 mW He/Ne laser operating at Abs = 632.8 nm and a refractive index detector (Agilent Serie 1200 RID®). Light intensity measurements were derived using the classical Rayleigh-Debye equation, enabling the deduction of Mw and Mn. The refractive index increment (dn/dc) was computed from a polynomial based on previous studies that correlates the dn/dc with the degree of acetylation (Schatz et al ). For these chitosans with DAs (%) between 5 and 15 the value of dn/dc employed in the analysis was 0.185. A 150 mM ammonium acetate buffer pH 4.5 adjusted with 200 mM acetic acid () was used as eluent. The flow rate was set to 0.6 mL/min. The chitosan samples were dissolved in the aforementioned ammonium acetate buffer at a final concentration of 1 mg/ml and stirred overnight at room temperature to ensure complete dissolution. The samples were afterwards filtrated using a 0.2 µm modified nylon filter (VWR, Randon, DE, USA) and 100 µl of each were injected into the HPSEC system.

M5: *Enzymatic method 1*

The chitosan content and its degree of acetylation in the feed samples was quantified using the method by Urs et al. [Ref] with the following modifications. The material containing chitosan was beat twice using 2-3 steel balls at full power (30 Hz) for 2.5 min (2x) with 2.5 min break using a Mixer Mill MM 400 (Retsch, Haan, Germany) to homogenise the sample. Then around 1 mg of the homogenized sample was used in the chemically *N*-acetylation step. For this step 0.5 ml sodium bicarbonate buffer (1 M) and 50 µL of $d_6$-acetic anhydride (Merck, Darmstadt, Germany) were added to the samples, immediately mixed and incubated for 20 min at room temperature under periodic gentle mixing. To achieve a full acetylation of the glucosamine units, the temperature was increased to 100°C for 5 min in the last step of this reaction. Afterwards, the samples were centrifuged at 13 000 g at 4°C for 10 minutes. The pellet was washed twice with water and in the last step resuspended in 200 µL ammonium acetate buffer (100 mM; pH 5.5). To this 5 µL of *Trichoderma viride* chitinase (Sigma-Aldrich, St-Louis, MO, USA), 1 µl of ChiB - chitinase ChiB from *Serratia marcescens* - (1 mg/mL) and 1 µL of CSN174 - recombinant chitosanase CSN-174 from

*Streptomyces* sp. N174 - (1 mg/mL) were added and incubated for 3 days at 37°C. The samples were freeze dried and, depending on the chitin and chitosan concentration in the starting material, dissolved in 100 – 500 µL $H_20$, filtered with a 3 kDa PES Eppi filter (VWR, Radnor, DE, USA) and finally mixed 1:1 with the internal standard R* _- double isotopically labelled [$^{13}C_2$,$^{2}H_3$] GlcNAc -(0.1 mg/ml). These samples were used for the UHPLC-MS analysis.

M6: *Enzymatic method 2*

The samples (100 µL with a final concentration of 5 mg/mL dissolved in 150 mM ammonium acetate buffer pH 4.5) were incubated at 37 °C with 5 µg ChiB and 5 µg CSN174 for 16 hours. Afterwards 400 µL of 100 mM $NaHCO_3$ buffer were added and the samples were centrifuged for 30 s at 13000 *g* at 20°C to precipitate all insoluble material, which were part of the feed sample. The supernatant containing the water-soluble chitin and chitosan oligomers was filtrated using a 0.2 µm nylon filter (VWR, Radnor, DE, USA) for 60 min at 13000 g and 20 °C and the filtrate was stored at -20 °C. The volume of each sample corresponding to 25 µg of chitosan in the starting material, was *N*-acetylated using $d_6$-acetic anhydride according to Cord-Landwehr et al. 2017. Afterwards the samples were freeze dried, dissolved in 100 µl ammonium acetate buffer (100 mM; pH 5.0) containing 0.1 units of *Trichoderma viride* chitinase (Sigma-Aldrich, St-Louis, MO, USA) and incubated for three days at 37°C. Each 2 µL of the sample, which was first filtered using the 3 kDa PES filter (VWR, Radnor, DE, USA), was mixed with 2 µl of R* (0.025 mg/mL) serving as internal standard and analysed using the UHPLC-MS method described above.
