## Supplementary material for "Comparative chemical characterisation of chitosans and their impact on growth, faecal consistency and microbiota composition in weaned piglets": Supplementary Tables_Submission.pdf

***Supplementary Table S1.*** *Diet composition and calculated nutrient values of the experimental diets for the first in vivo trial at SFR (VOF-61). Weaner I was fed until day x of age and Weaner II thereafter*

| Ingredients (%) | Weaner I | Weaner II |
|---|---|---|
| Wheat | 28.516 | 25.012 |
| Maize | 20.000 | 25.000 |
| Soybean meal (>48% crude protein) | 16.000 | 15.453 |
| Barley | 15.000 | 20.000 |
| Milkpowder skimmed | 5.241 | 0.000 |
| Wheat middlings | 3.295 | 2.500 |
| Molasses beet | 3.000 | 3.000 |
| Potato protein - Protastar | 2.531 | 1.000 |
| Animal fat - Pork | 1.639 | 2.049 |
| Water | 1.000 | 1.000 |
| Limestone | 0.696 | 1.299 |
| Monocalcium phosphate | 0.532 | 0.751 |
| Salt | 0.508 | 0.569 |
| Soybean oil | 0.500 | 0.000 |
| Calcium formate | 0.500 | 0.000 |
| Vitamin and mineral premix[1] | 0.500 | 0.500 |
| Soy protein concentrate - Soycomil | 0.000 | 0.821 |
| Lysine-HCL (79%) | 0.271 | 0.521 |
| Methionine (DL 99%) | 0.114 | 0.193 |
| Threonine (L 98%) | 0.087 | 0.205 |
| Tryptophan (L 98%) | 0.019 | 0.051 |
| Valine (L 99%) | 0.000 | 0.045 |
| Phytase (0-500) | 0.002 | 0.002 |
| Phytase (500-750) | 0.000 | 0.001 |
| Copper sulphate (99%) | 0.049 | 0.028 |
| | **100.00** | **100.00** |

| Nutrients | Unit | | |
|---|---|---|---|
| Moisture | g/kg | 130.46 | 133.30 |
| Ash | g/kg | 48.78 | 53.03 |
| Crude protein | g/kg | 190.00 | 170.22 |
| Crude fat (AH) | g/kg | 45.45 | 45.27 |
| Crude fibre | g/kg | 24.71 | 26.39 |
| Starch (AM) | g/kg | 379.15 | 414.27 |
| Sugar | g/kg | 67.50 | 41.28 |
| Ca | g/kg | 6.41 | 6.87 |
| P | g/kg | 4.83 | 4.92 |
| Ca/P | ratio | 1.33 | 1.40 |
| avCa | g/kg | 7.00 | 7.50 |
| ATTD P-s intense | g/kg | 3.35 | 3.37 |
| STTD P-s intense | g/kg | 3.51 | 3.52 |
| Mg | g/kg | 1.55 | 1.38 |
| K | g/kg | 8.41 | 7.79 |
| Na | g/kg | 2.50 | 2.50 |
| Cl | g/kg | 4.90 | 5.23 |
| S total | g/kg | 2.26 | 2.04 |
| dEB | meq/kg | 186.34 | 161.01 |
| ABC4 | meq/kg | 322.19 | 384.07 |
| FTU | kFTU | 0.42 | 0.64 |
| Zn | mg/kg | 31.99 | 30.60 |
| Cu | mg/kg | 150.00 | 100.00 |
| E-Piglet | MJ/kg | 9.59 | 9.59 |
| SID LYS-s | g/kg | 11.00 | 11.00 |
| SID Lys/ E-piglet | ratio | 1.15 | 1.15 |
| SID MET/SID LYS-s | ratio | 0.37 | 0.38 |
| SID M+C/SID LYS-s | ratio | 0.60 | 0.60 |
| SID THR/SID LYS-s | ratio | 0.65 | 0.65 |
| SID TRP/SID LYS-s | ratio | 0.20 | 0.20 |
| SID VAL/SID LYS-s | ratio | 0.75 | 0.66 |
| SID ILE/SID LYS-s | ratio | 0.65 | 0.54 |
| SID LEU/SID LYS-s | ratio | 1.25 | 1.04 |
| SID ARG/SID LYS-s | ratio | 0.91 | 0.82 |
| SID HIS/SID LYS-s | ratio | 0.39 | 0.34 |
| SID PHE/SID LYS-s | ratio | 0.77 | 0.66 |
| NSP-s total | g/kg | 135.76 | 141,48 |
| AID NSP-s | g/kg | 88.40 | 91.32 |
| fCHO-s | g/kg | 97.46 | 100.13 |
| iCHO-s | g/kg | 47.33 | 50.13 |

[1] A vitamin and mineral premix was added at 0.50% to provide the following nutrients per kg of diet: vitamin A: 10000 IU, vitamin D3: 2000 IU, vitamin E: 40 mg, vitamin K: 1.5 mg, vitamin B1: 1 mg, vitamin B2: 4 mg, vitamin B6: 1.5 mg, vitamin B12: 0.02 mg, niacin: 30 mg, D-pantothenic acid: 15 mg, choline chloride: 150 mg, folate: 0.4 mg, biotin: 0.05 mg, iron: 100 mg, copper: 20 mg, manganese: 30 mg, zinc: 70 mg, iodine: 0.7 mg, selenium: 0.25 mg, anti-oxidant: 125 mg.

***Supplementary Table S2.*** *Diet composition and calculated nutrient values of the experimental diets for the first in vivo trial at SFR (VOB-62). Weaner I was fed until day x of age and Weaner II thereafter*

| **Ingredients (%)** | **Weaner I** | **Weaner II** |
|---|---|---|
| Wheat | 35.000 | 35.000 |
| Barley | 16.020 | 25.823 |
| Maize | 15.000 | 10.000 |
| Soybean meal (>48% crude protein) | 12.500 | 25.000 |
| Wheat middlings | 4.241 | 5.000 |
| Milkpowder skimmed | 4.193 | 0.000 |
| Potato protein - Protastar | 3.811 | 2.644 |
| Oat hulls | 2.000 | 0.000 |
| Molasses beet | 1.872 | 2.000 |
| Animal fat - Pork | 1.116 | 0.527 |
| Salt | 0.663 | 0.578 |
| Monocalcium phosphate | 0.639 | 0.606 |
| Limestone | 0.516 | 1.340 |
| Soybean oil | 0.500 | 0.000 |
| Calcium formate | 0.500 | 0.000 |
| Vitamin and mineral premix[1] | 0.500 | 0.500 |
| Lysine-HCL (79%) | 0.465 | 0.514 |
| Methionine (L/DL 99%) | 0.193 | 0.190 |
| Threonine (L 98%) | 0.171 | 0.189 |
| Tryptophan (L 98%) | 0.046 | 0.041 |
| Valine (L 99%) | 0.000 | 0.013 |
| Phytase (0-500) | 0.0025 | 0.0025 |
| Copper sulphate (99%) | 0.051 | 0.031 |
| | **100.00** | **100.00** |

| **Nutrients** | **Unit** | | |
|---|---|---|---|
| Moisture | g/kg | 118.88 | 122.98 |
| Ash | g/kg | 47.11 | 51.45 |
| Crude protein | g/kg | 189.79 | 185.77 |
| Crude fat (AH) | g/kg | 40.69 | 30.31 |
| Crude fibre | g/kg | 31.44 | 30.64 |
| Starch (AM) | g/kg | 394.04 | 413.93 |
| Sugar | g/kg | 56.48 | 39.78 |
| Ca | g/kg | 5.80 | 6.80 |
| P | g/kg | 4.98 | 4.86 |
| Ca/P | ratio | 1.16 | 1.40 |
| avCa | g/kg | 6.50 | 7.50 |
| ATTD P-s intense | g/kg | 3.60 | 3.27 |
| STTD P-s intense | g/kg | 3.77 | 3.45 |
| Mg | g/kg | 1.52 | 1.52 |
| K | g/kg | 7.42 | 7.64 |
| Na | g/kg | 3.00 | 2.50 |
| Cl | g/kg | 6.08 | 5.32 |
| S total | g/kg | 2.58 | 2.39 |
| dEB | meq/kg | 149.24 | 154.59 |
| ABC4 | meq/kg | 274.80 | 399.63 |
| FTU | kFTU | 0.50 | 0.50 |
| Zn | mg/kg | 102.27 | 104.29 |
| Cu | mg/kg | 150.00 | 100.00 |
| E-Piglet | MJ/kg | 9.50 | 9.25 |
| SID LYS-s | g/kg | 12.35 | 11.84 |
| SID Lys/ E-piglet | ratio | 1.30 | 1.28 |
| SID MET/SID LYS-s | ratio | 0.39 | 0.38 |
| SID M+C/SID LYS-s | ratio | 0.60 | 0.60 |
| SID THR/SID LYS-s | ratio | 0.65 | 0.65 |
| SID TRP/SID LYS-s | ratio | 0.20 | 0.20 |
| SID VAL/SID LYS-s | ratio | 0.67 | 0.66 |
| SID ILE/SID LYS-s | ratio | 0.57 | 0.56 |
| SID LEU/SID LYS-s | ratio | 1.10 | 1.04 |
| SID ARG/SID LYS-s | ratio | 0.78 | 0.83 |
| SID HIS/SID LYS-s | ratio | 0.34 | 0.34 |
| SID PHE/SID LYS-s | ratio | 0.69 | 0.68 |

[1] A vitamin and mineral premix was added at 0.50% to provide the following nutrients per kg of diet: vitamin A: 10000 IU, vitamin D3: 2000 IU, vitamin E: 40 mg, vitamin K: 1.5 mg, vitamin B1: 1 mg, vitamin B2: 4 mg, vitamin B6: 1.5 mg, vitamin B12: 0.02 mg, niacin: 30 mg, D-pantothenic acid: 15 mg, choline chloride: 150 mg, folate: 0.4 mg, biotin: 0.05 mg, iron: 100 mg, copper: 20 mg, manganese: 30 mg, zinc: 70 mg, iodine: 0.7 mg, selenium: 0.25 mg, anti-oxidant: 125 mg.

***Supplementary Table S3. Power calculations***

Power calculations VOF-61 – GenStat® output

- Key endpoint: Average daily gain between day 0-42 post-weaning, in grams

```
SCALAR diff; VALUE = 41
 SCALAR se; VALUE = 12.7
 CALC sd = se * SQRT ( 6 )
 CALC component = sd ** 2
 AGHIERARCHICAL [ PRINT=*; ANALYSE=no; SEED=-1] Block,Plot;\
              TREATMENTFACTORS=*,Treat; LEVELS=1,2   " 2 = NUMBER OF TREATMENTS / BLOCK "

  ASAMPLESIZE    [PRINT=power,rep; TERM=Treat; REPLICATES=Block; TMETHOD = twosided; \
               TREATMENTSTRUCTURE=Treat; BLOCKSTRUCTURE=Block/Plot;\
              COMPONENT=component; POWER = 0.8 ] diff; NREPLICATES=Nrep
```

### Sample size for analysis of variance

#### Power

| Number of replicates | Residual d.f. | Residual m.s. | s.e.d. | RESPONSE / s.e.d. | t-value | Power |
|---|---|---|---|---|---|---|
| 7 | 6 | 967.7 | 16.63 | 2.466 | 2.447 | 0.543 |
| 8 | 7 | 967.7 | 15.55 | 2.636 | 2.365 | 0.621 |
| 9 | 8 | 967.7 | 14.66 | 2.796 | 2.306 | 0.689 |
| 10 | 9 | 967.7 | 13.91 | 2.947 | 2.262 | 0.746 |
| 11 | 10 | 967.7 | 13.26 | 3.091 | 2.228 | 0.795 |
| 12 | 11 | 967.7 | 12.70 | 3.228 | 2.201 | 0.836 |
| 13 | 12 | 967.7 | 12.20 | 3.360 | 2.179 | 0.869 |
| 14 | 13 | 967.7 | 11.76 | 3.487 | 2.160 | 0.896 |
| 15 | 14 | 967.7 | 11.36 | 3.609 | 2.145 | 0.918 |
| 16 | 15 | 967.7 | 11.00 | 3.728 | 2.131 | 0.936 |
| 17 | 16 | 967.7 | 10.67 | 3.843 | 2.120 | 0.950 |

#### Replication

To detect a treatment difference of 41.00, at a significance level of 0.050, with a power of 0.800, using a two-sided test, requires a replication of 12.

```
SCALAR diff; VALUE = 0.037
 SCALAR se; VALUE = 0.01
 CALC sd = se * SQRT ( 6 )
 CALC component = sd ** 2
 AGHIERARCHICAL [ PRINT=*; ANALYSE=no; SEED=-1] Block,Plot;\
              TREATMENTFACTORS=*,Treat; LEVELS=1,2   " 2 = NUMBER OF TREATMENTS / BLOCK "

  ASAMPLESIZE    [PRINT=power,rep; TERM=Treat; REPLICATES=Block; TMETHOD = twosided; \
               TREATMENTSTRUCTURE=Treat; BLOCKSTRUCTURE=Block/Plot;\
              COMPONENT=component; POWER = 0.8 ] diff; NREPLICATES=Nrep
```

#### Sample size for analysis of variance

##### Power

| Number of replicates | Residual d.f. | Residual m.s. | s.e.d. | RESPONSE / s.e.d. | t-value | Power |
|---|---|---|---|---|---|---|
| 4 | 3 | 0.0006000 | 0.01732 | 2.136 | 3.182 | 0.320 |
| 5 | 4 | 0.0006000 | 0.01549 | 2.388 | 2.776 | 0.445 |
| 6 | 5 | 0.0006000 | 0.01414 | 2.616 | 2.571 | 0.559 |
| 7 | 6 | 0.0006000 | 0.01309 | 2.826 | 2.447 | 0.656 |
| 8 | 7 | 0.0006000 | 0.01225 | 3.021 | 2.365 | 0.736 |
| 9 | 8 | 0.0006000 | 0.01155 | 3.204 | 2.306 | 0.801 |
| 10 | 9 | 0.0006000 | 0.01095 | 3.378 | 2.262 | 0.851 |
| 11 | 10 | 0.0006000 | 0.01044 | 3.542 | 2.228 | 0.890 |
| 12 | 11 | 0.0006000 | 0.01000 | 3.700 | 2.201 | 0.920 |
| 13 | 12 | 0.0006000 | 0.00961 | 3.851 | 2.179 | 0.942 |
| 14 | 13 | 0.0006000 | 0.00926 | 3.996 | 2.160 | 0.958 |

##### Replication

To detect a treatment difference of 0.03700, at a significance level of 0.050, with a power of 0.800, using a two-sided test, requires a replication of 9.

*Power calculations VOB-62 – GenStat® output*

Data from Xu et al. (2018) were used, in which the effect of supplementing the diet with chitosan at the inclusion of 500 mg/kg was tested on weaner pig performance, measured between day 0 and 14 post-weaning, with 12 piglets per treatment group (individually housed).

- Key endpoint: Average daily gain between day 0-14 post-weaning, in grams

```
SCALAR diff; VALUE = 56
SCALAR se; VALUE = 8
CALC sd = se * SQRT ( 12 )
CALC component = sd ** 2
AGHIERARCHICAL [ PRINT=*; ANALYSE=no; SEED=-1] Block,Plot;\
               TREATMENTFACTORS=*,Treat; LEVELS=1,2  " 2 = NUMBER OF TREATMENTS / BLOCK "

ASAMPLESIZE    [PRINT=power,rep; TERM=Treat; REPLICATES=Block; TMETHOD = twosided; \
               TREATMENTSTRUCTURE=Treat; BLOCKSTRUCTURE=Block/Plot;\
               COMPONENT=component; POWER = 0.8 ] diff; NREPLICATES=Nrep
```

### Sample size for analysis of variance

#### Power

| Number of replicates | Residual d.f. | Residual m.s. | s.e.d. | RESPONSE / s.e.d. | t-value | Power |
|---|---|---|---|---|---|---|
| 2 | 1 | 768.0 | 27.71 | 2.021 | 12.706 | 0.126 |
| 3 | 2 | 768.0 | 22.63 | 2.475 | 4.303 | 0.295 |
| 4 | 3 | 768.0 | 19.60 | 2.858 | 3.182 | 0.497 |
| 5 | 4 | 768.0 | 17.53 | 3.195 | 2.776 | 0.671 |
| 6 | 5 | 768.0 | 16.00 | 3.500 | 2.571 | 0.797 |
| 7 | 6 | 768.0 | 14.81 | 3.780 | 2.447 | 0.880 |
| 8 | 7 | 768.0 | 13.86 | 4.041 | 2.365 | 0.931 |
| 9 | 8 | 768.0 | 13.06 | 4.287 | 2.306 | 0.962 |
| 10 | 9 | 768.0 | 12.39 | 4.518 | 2.262 | 0.979 |
| 11 | 10 | 768.0 | 11.82 | 4.739 | 2.228 | 0.989 |
| 12 | 11 | 768.0 | 11.31 | 4.950 | 2.201 | 0.994 |

#### Replication

To detect a treatment difference of 56.00, at a significance level of 0.050, with a power of 0.800, using a two-sided test, requires a replication of 7.

```
PRIN Nrep
```

Nrep
7.000

```
stop
```

End of job.

- Key endpoint: Feed conversion ratio (daily feed intake/daily weight gain) between day 0-14 post-weaning, in g/g

```
SCALAR diff; VALUE = 0.32
SCALAR se; VALUE = 0.07
CALC sd = se * SQRT ( 12 )
CALC component = sd ** 2
AGHIERARCHICAL [ PRINT=*; ANALYSE=no; SEED=-1] Block,Plot;\
                     TREATMENTFACTORS=*,Treat; LEVELS=1,2  " 2 = NUMBER OF TREATMENTS / BLOCK "

ASAMPLESIZE    [PRINT=power,rep; TERM=Treat; REPLICATES=Block; TMETHOD = twosided; \
                     TREATMENTSTRUCTURE=Treat; BLOCKSTRUCTURE=Block/Plot;\
                     COMPONENT=component; POWER = 0.8 ] diff; NREPLICATES=Nrep
```

### Sample size for analysis of variance

#### Power

| Number of replicates | Residual d.f. | Residual m.s. | s.e.d. | RESPONSE / s.e.d. | t-value | Power |
|---|---|---|---|---|---|---|
| 7 | 6 | 0.05880 | 0.1296 | 2.469 | 2.447 | 0.544 |
| 8 | 7 | 0.05880 | 0.1212 | 2.639 | 2.365 | 0.622 |
| 9 | 8 | 0.05880 | 0.1143 | 2.799 | 2.306 | 0.690 |
| 10 | 9 | 0.05880 | 0.1084 | 2.951 | 2.262 | 0.748 |
| 11 | 10 | 0.05880 | 0.1034 | 3.095 | 2.228 | 0.796 |
| 12 | 11 | 0.05880 | 0.0990 | 3.232 | 2.201 | 0.837 |
| 13 | 12 | 0.05880 | 0.0951 | 3.364 | 2.179 | 0.870 |
| 14 | 13 | 0.05880 | 0.0917 | 3.491 | 2.160 | 0.897 |
| 15 | 14 | 0.05880 | 0.0885 | 3.614 | 2.145 | 0.919 |
| 16 | 15 | 0.05880 | 0.0857 | 3.733 | 2.131 | 0.936 |
| 17 | 16 | 0.05880 | 0.0832 | 3.847 | 2.120 | 0.950 |

#### Replication

To detect a treatment difference of 0.3200, at a significance level of 0.050,
with a power of 0.800, using a two-sided test, requires a replication of 12.

```
11  PRIN Nrep
```

| Nrep |
|---|
| 12.00 |

```
12
13
14  stop
```

End of job.

***Supplementary Table S4.*** *Faecal consistency 8-point scale*

SCHOTHORST FEED RESEARCH

### Faecal consistency weaner piglets (8-point scale)

| Picture | Score | Description | Explanation |
|---|---|---|---|
| 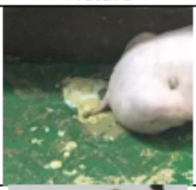 | 2 | Severe water-thin diarrhoea | Thin liquid, like water. Flows through slatted floor. |
| 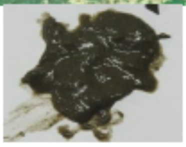 | 3 | Water-thin faeces | Flows through slatted floor |
| 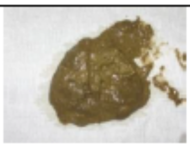 | 4 | Like custard | |
| 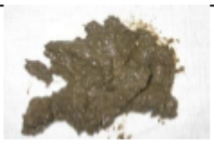 | 5 | Smooth | Shapeless pile |
| 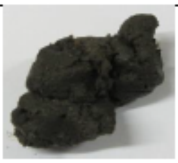 | 6 | Solid dropping without structure (mushy) | Like peanut butter. Sticks to glove when picking up. |
| 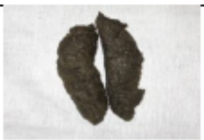 | 7 | Firm and shaped | |
| 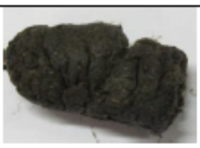 | 8 | Firm and shaped with structure | Easy to pick it up as a whole. Cracks on the surface. |
| 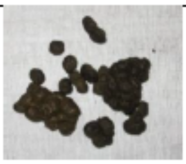 | 9 | Lumpy | Falls apart after shaking. If you pick it up your glove does hardly or not become dirty. |
| | 10 | No faeces present | |

***Supplementary Table S5.*** *Overview of mortality between day 0 – 42 post-weaning (PW) for the first SFR study (VOF-61). Trt = dietary treatment.*

| Piglet | Sex | Room | Pen | Replicate | Trt | Batch | Day PW | Date | BW, kg | Reason |
|---|---|---|---|---|---|---|---|---|---|---|
| 2830 | Female | 10 | 9 | 2 | 2 | 1 | 16 | 03-11-21 | 15.0 | Meningitis |
| 3186 | Male | 18 | 2 | 11 | 2 | 2 | 18 | 26-11-21 | 8.7 | Meningitis |
| 3264 | Male | 16 | 2 | 6 | 3 | 2 | 32 | 10-12-21 | 19.2 | Meningitis |
| 2536 | Male | 15 | 4 | 3 | 4 | 1 | 39 | 26-11-21 | 20.1 | Meningitis |

***Supplementary Table S6****. Overview of antibiotic treatments between day 0 – 42 post-weaning (PW). Trt = dietary treatment. Repl = replicate.*

| Piglet | Room | Pen | Repl | Trt | Batch | Day PW | Symptom | Product | Start | End | Duration |
|---|---|---|---|---|---|---|---|---|---|---|---|
| 3108[1] | 7 | 10 | 8 | 1 | 2 | 17 | Arthritis | Cyclosol + Ketoprosol | 25-11 | 07-12 | 12 |
| 3108[1] | 7 | 10 | 8 | 1 | 2 | 35 | Pneumonia | Cyclosol | 13-12 | 19-12 | 6 |
| 2830[2] | 10 | 9 | 2 | 2 | 1 | 14 | Meningitis | Cyclosol | 01-11 | 03-11 | 3 |
| 3186[3] | 18 | 2 | 11 | 2 | 2 | 15 | Meningitis | Cyclosol + Ketoprosol | 23-11 | 29-11 | 6 |
| 2287 | 15 | 4 | 3 | 4 | 1 | 10 | Arthritis | Cyclosol + Ketoprosol | 28-10 | 31-10 | 3 |
| 2536[4] | 15 | 4 | 3 | 4 | 1 | 13 | Arthritis | Cyclosol + Ketoprosol | 31-10 | 03-11 | 3 |
| 2536[4] | 15 | 4 | 3 | 4 | 1 | 26 | Meningitis | Cyclosol | 13-11 | 16-11 | 3 |
| 3228 | 17 | 4 | 9 | 4 | 2 | 41 | Eye infection | Depocillin | 19-12 | 20-12 | 1 |

[1] The piglet was treated for arthritis and pneumonia.

[2] The piglet died on 03-11-21.

[3] The piglet died on 26-11-21.

[4] The piglet was treated for arthritis and meningitis and died on 26-11-21.

***Supplementary Table S7.*** *Overview of the medical treatments per dietary treatment between day 0 – 42 post-weaning.*

| Treatment | 1 | 2 | 3 | 4 | |
|---|---|---|---|---|---|
| | Negative control (NC) | NC + low dose COS | NC + medium dose COS | NC +high dose COS | Total |
| Arthritis | 1 | 0 | 0 | 2 | 3 |
| Meningitis | 0 | 2 | 0 | 1 | 3 |
| Other | 1 | 0 | 0 | 1 | 2 |
| Total | 2 | 2 | 0 | 4 | 8 |

***Supplementary Table S8.*** *Descriptive statistics (mean ± SEM) of the effect of a low, medium and high dosage of chito-oligosaccharide (COS) in phase 1 and 2 weaner diets compared to control diets (CON) on piglet performance in batch 1 and 2 of the weaner phase. Day represents day post-weaning.*

| | Batch 1 (n=20 pens) | | | | Batch 2 (n=28 pens) | | | |
|---|---|---|---|---|---|---|---|---|
| | CON (n=5) | COS-low (n=5) | COS-medium (n=5) | COS-high (n=5) | CON (n=7) | COS-low (n=7) | COS-medium (n=7) | COS-high (n=7) |
| Body weight, kg | | | | | | | | |
| Day 0 | 9.56 ± 0.122 | 9.56 ± 0.123 | 9.56 ± 0.121 | 9.56 ± 0.124 | 9.23 ± 0.173 | 9.23 ± 0.172 | 9.23 ± 0.172 | 9.24 ± 0.173 |
| Day 14 | 14.23 ± 0.315 | 13.29 ± 0.370 | 13.74 ± 0.350 | 13.92 ± 0.472 | 13.30 ± 0.295 | 13.43 ± 0.139 | 13.46 ± 0.294 | 13.21 ± 0.277 |
| Day 42 | 35.65 ± 1.034 | 33.18 ± 1.029 | 33.30 ± 0.595 | 34.58 ± 1.648 | 32.82 ± 1.025 | 32.28 ± 0.763 | 33.30 ± 0.600 | 32.79 ± 1.023 |
| CV, % | | | | | | | | |
| Day 0 | 0.85 ± 0.181 | 0.83 ± 0.123 | 0.95 ± 0.126 | 0.87 ± 0.126 | 0.92 ± 0.114 | 1.05 ± 0.132 | 0.94 ± 0.099 | 0.74 ± 0.080 |
| Day 14 | 10.11 ± 0.991 | 7.35 ± 1.301 | 8.51 ± 2.455 | 9.36 ± 1.582 | 6.19 ± 0.639 | 6.80 ± 1.276 | 8.18 ± 1.090 | 7.79 ± 0.812 |
| Day 42 | 12.26 ± 2.436 | 9.79 ± 1.212 | 9.72 ± 2.313 | 12.42 ± 0.979 | 9.99 ± 2.491 | 6.58 ± 1.168 | 10.13 ± 1.681 | 8.89 ± 1.289 |
| ADG, g/piglet | | | | | | | | |
| Day 0 – 14 | 334 ± 18.8 | 266 ± 19.4 | 299 ± 18.2 | 311 ± 30.0 | 291 ± 18.2 | 300 ± 11.7 | 302 ± 12.6 | 284 ± 12.2 |
| Day 14 – 42 | 765 ± 26.0 | 711 ± 25.6 | 698 ± 15.3 | 738 ± 43.4 | 697 ± 28.7 | 673 ± 25.2 | 708 ± 13.3 | 699 ± 30.7 |
| Day 0 – 42 | 621 ± 23.1 | 562 ± 22.7 | 565 ± 11.9 | 596 ± 37.8 | 562 ± 23.8 | 549 ± 17.3 | 573 ± 12.0 | 561 ± 23.6 |
| ADFI, g/piglet | | | | | | | | |
| Day 0 – 14 | 448 ± 28.5 | 373 ± 15.8 | 427 ± 26.2 | 452 ± 36.2 | 374 ± 19.8 | 383 ± 10.7 | 394 ± 13.2 | 382 ± 18.4 |
| Day 14 – 42 | 1087 ± 71.2 | 1018 ± 57.4 | 980 ± 46.7 | 1052 ± 62.7 | 1044 ± 40.7 | 991 ± 27.6 | 1002 ± 37.2 | 991 ± 49.4 |
| Day 0 – 42 | 874 ± 56.2 | 803 ± 42.5 | 795 ± 38.3 | 852 ± 47.8 | 821 ± 32.5 | 788 ± 17.9 | 799 ± 27.2 | 788 ± 38.3 |
| FCR, g/g | | | | | | | | |
| Day 0 – 14 | 1.34 ± 0.023 | 1.42 ± 0.059 | 1.43 ± 0.041 | 1.46 ± 0.046 | 1.29 ± 0.023 | 1.28 ± 0.031 | 1.31 ± 0.038 | 1.35 ± 0.020 |
| Day 14 – 42 | 1.41 ± 0.048 | 1.43 ± 0.067 | 1.40 ± 0.056 | 1.44 ± 0.082 | 1.50 ± 0.032 | 1.48 ± 0.045 | 1.41 ± 0.048 | 1.42 ± 0.041 |
| Day 0 – 42 | 1.41 ± 0.041 | 1.43 ± 0.059 | 1.41 ± 0.050 | 1.44 ± 0.069 | 1.46 ± 0.026 | 1.44 ± 0.039 | 1.40 ± 0.044 | 1.41 ± 0.034 |
| Faecal consistency score | | | | | | | | |
| Day 0 – 14 | 5.76 ± 0.256 | 5.96 ± 0.160 | 5.92 ± 0.314 | 5.88 ± 0.280 | 6.11 ± 0.213 | 6.31 ± 0.278 | 6.47 ± 0.256 | 6.14 ± 0.253 |
| Day 14 – 42 | 6.39 ± 0.105 | 6.27 ± 0.08 | 6.27 ± 0.187 | 6.30 ± 0.178 | 6.52 ± 0.147 | 6.29 ± 0.140 | 6.45 ± 0.144 | 6.24 ± 0.209 |
| Day 0 – 42 | 6.08 ± 0.179 | 6.17 ± 0.059 | 6.10 ± 0.252 | 6.12 ± 0.174 | 6.33 ± 0.096 | 6.30 ± 0.146 | 6.41 ± 0.156 | 6.14 ± 0.220 |

***Supplementary Table S9***. *The effect of experimental diet on piglet performance between day 0-42 post-weaning, excluding outliers for the first SFR study (VOF-61).*

| **Treatment** | **1** | **2** | **3** | **4** | | | |
|---|---|---|---|---|---|---|---|
| Description | Negative control (NC) | NC + low dose | NC + medium dose | NC + high dose | SEM[1] | LSD[2] | *P* - value |
| Body weight, kg | | | | | | | |
| Day 0 | 9.37 | 9.37 | 9.37 | 9.37 | 0.002 | 0.007 | 0.39 |
| Day 14 | 13.69 | 13.37 | 13.58 | 13.51 | 0.159 | 0.456 | 0.56 |
| Day 42 | 34.00 | 32.65 | 33.30 | 33.53 | 0.610 | 1.755 | 0.48 |
| CV in body weight, % | | | | | | | |
| Day 0 | 0.89 | 0.96 | 0.94 | 0.80 | 0.060 | 0.174 | 0.24 |
| Day 14 | 7.83 | 7.03 | 8.32 | 8.44 | 0.921 | 2.650 | 0.70 |
| Day 42 | 10.93 | 7.92 | 9.96 | 10.36 | 1.250 | 3.595 | 0.37 |
| ADG, g/d/piglet | | | | | | | |
| Day 0 – 14 | 309 | 286 | 301 | 295 | 11.3 | 32.5 | 0.55 |
| Day 14 – 42 | 725 | 689 | 704 | 715 | 17.9 | 51.5 | 0.52 |
| Day 0 – 42 | 587 | 554 | 570 | 575 | 14.5 | 41.7 | 0.48 |
| ADFI, g/d/piglet | | | | | | | |
| Day 0 – 14 | 405 | 379 | 408 | 411 | 13.8 | 39.7 | 0.34 |
| Day 14 – 42 | 1062 | 1002 | 992 | 1017 | 24.6 | 70.6 | 0.22 |
| Day 0 – 42 | 843 | 794 | 798 | 815 | 19.1 | 55.0 | 0.28 |
| FCR, g/g | | | | | | | |
| Day 0 – 14 | 1.31[a] | 1.34[ab] | 1.36[ab] | 1.40[b] | 0.020 | 0.059 | **0.045** |
| Day 14 – 42 | 1.47 | 1.46 | 1.41 | 1.43 | 0.024 | 0.067 | 0.31 |
| Day 0 – 42 | 1.44 | 1.44 | 1.40 | 1.42 | 0.022 | 0.062 | 0.58 |
| Faecal consistency score[3] | | | | | | | |
| Day 0 – 7 | 6.00 | 6.19 | 6.14 | 6.14 | 0.173 | 0.499 | 0.88 |
| Day 7 – 14 | 6.00 | 6.22 | 6.22 | 5.86 | 0.175 | 0.504 | 0.40 |
| Day 0 – 14 | 5.97 | 6.17 | 6.18 | 6.03 | 0.159 | 0.459 | 0.74 |
| Day 14 – 42 | 6.47 | 6.28 | 6.38 | 6.26 | 0.095 | 0.274 | 0.41 |
| Day 0 – 42 | 6.23 | 6.24 | 6.28 | 6.13 | 0.112 | 0.321 | 0.82 |

***Supplementary Table S10***. *Overview of antibiotic treatments between day 0-42 post-weaning (PW). Trt = dietary treatment. Repl = replicate.*

| Piglet | Room | Pen | Repl | Trt | Batch | Day PW | Symptom | Product | Start | End | Duration |
|---|---|---|---|---|---|---|---|---|---|---|---|
| 4806 | 5 | 4 | 2 | 1 | 1 | 23 | Pneumonia | Cyclosol | 30-04-22 | 03-05-22 | 3 |
| 4966 | 20 | 3 | 7 | 1 | 1 | 8 | Meningitis | Cyclosol | 15-04-22 | 01-05-22 | 16 |
| 9990[1] | 5 | 5 | 10 | 1 | 2 | 14 | Meningitis | Cyclosol | 23-06-22 | 24-06-22 | 1 |
| 7804 | 20 | 2 | 11 | 2 | 2 | 6 | Arthritis | Cyclosol | 15-06-22 | 18-06-22 | 3 |
| 7823[2] | 20 | 2 | 11 | 2 | 2 | 18 | Arthritis | Cyclosol | 27-06-22 | 30-06-22 | 3 |
| 9830 | 20 | 5 | 12 | 2 | 2 | 2 | General weakness | Cyclosol | 11-06-22 | 17-06-22 | 6 |
| 4821 | 6 | 2 | 3 | 3 | 1 | 21 | Meningitis | Cyclosol | 28-04-22 | 01-05-22 | 3 |
| 4878 | 19 | 2 | 5 | 3 | 1 | 41 | Arthritis | Cyclosol | 18-05-22 | 19-05-22 | 1 |
| 8527 | 5 | 2 | 9 | 3 | 2 | 14 | Arthritis | Cyclosol | 23-06-22 | 26-06-22 | 3 |
| 9872 | 5 | 4 | 10 | 3 | 2 | 11 | Arthritis | Cyclosol | 20-06-22 | 23-06-22 | 3 |
| Unknown | 5 | 4 | 10 | 3 | 2 | 35 | Arthritis | Cyclosol | 14-07-22 | 17-07-22 | 3 |
| 9420 | 20 | 1 | 11 | 3 | 2 | 10 | Meningitis | Cyclosol | 19-06-22 | 28-06-22 | 9 |
| 9469[3] | 20 | 1 | 11 | 3 | 2 | 24 | Pneumonia | Cyclosol | 03-07-22 | 06-07-22 | 3 |
| 9469[3] | 20 | 1 | 11 | 3 | 2 | 36 | Pneumonia | Cyclosol | 15-07-22 | 15-07-22 | 1 |

[1] The piglet died on day 15 post-weaning.

[2] The piglet died on day 29 post-weaning.

[3] The piglet was treated for pneumonia twice and died on day 36 post-weaning.

***Supplementary Table S11.*** *Overview of the medical treatments per dietary treatment.*

| Treatment | 1<br>Negative control (NC) | 2<br>NC + LMW chitosan | 3<br>NC + MMW chitosan | Total (n) | Total (%) |
|---|---|---|---|---|---|
| n of animals | 96 | 96 | 96 | 288 | |
| Arthritis | 0 | 2 | 4 | 6 | 2.08 |
| Meningitis | 2 | 0 | 2 | 4 | 1.39 |
| Pneumonia | 1 | 0 | 1[1] | 2 | 0.69 |
| General weakness | 0 | 1 | 0 | 1 | 0.35 |
| Total (n) | 3 | 3 | 7 | 13 | 4.51 |
| Total (%) | 1.04 | 1.04 | 2.43 | 4.51 | |

[1] The piglet was treated twice for pneumonia.

***Supplementary Table S12***. *Overview of mortality between day 0-42 post-weaning (PW). Trt = dietary treatment.*

| Piglet | Sex | Room | Pen | Replicate | Trt | Weaning batch | Day PW[1] | Date | BW, kg | Reason |
|---|---|---|---|---|---|---|---|---|---|---|
| 5285 | Female | 19 | 4 | 6 | 1 | 1 | 15 | 22-04-22 | Unknown | Unknown |
| 9990 | Male | 5 | 5 | 10 | 1 | 2 | 15 | 24-06-22 | 6.7 | Meningitis |
| 5278 | Male | 20 | 2 | 7 | 2 | 1 | 13 | 20-04-22 | 9.6 | Pneumonia |
| 7823 | Male | 20 | 2 | 11 | 2 | 2 | 29 | 08-07-22 | 7.5 | Meningitis |
| 9469 | Male | 20 | 1 | 11 | 3 | 2 | 36 | 15-07-22 | 12.0 | Removed from pen due to meningitis |
| 9979 | Male | 20 | 4 | 12 | 3 | 2 | 28 | 07-07-22 | 12.2 | Meningitis |

[1]When calculating ADFI based on number of animal days (number of animals x number of days) feed intake of these animals was taken into account till the days they were excluded from the experiment.

***Supplementary Table S13***. *Descriptive statistics (mean ± SEM) of the effect of low and high molecular weight chitosan in phase 1 and 2 weaner diets compared to control diets (CON) on piglet performance in batch 1 and 2 of the weaner phase. Day represents day post-weaning.*

| | Batch 1 (n=24 pens) | | | Batch 2 (n=12 pens) | | |
|---|---|---|---|---|---|---|
| | CON (n=8) | LMW-C (n=8) | MMW-C (n=8) | CON (n=4) | LMW-C (n=8) | MMW-C (n=8) |
| Body weight, kg | | | | | | |
| Day 0 | 7.75 ± 0.273 | 7.75 ± 0.274 | 7.75 ± 0.273 | 7.46 ± 0.251 | 7.46 ± 0.254 | 7.46 ± 0.248 |
| Day 14 | 11.71 ± 0.397 | 11.61 ± 0.508 | 11.44 ± 0.431 | 10.63 ± 0.259 | 10.35 ± 0.117 | 10.59 ± 0.483 |
| Day 42 | 30.04 ± 0.912 | 30.48 ± 0.896 | 30.99 ± 0.650 | 29.59 ± 0.736 | 27.85 ± 0.885 | 28.10 ± 1.207 |
| CV, % | | | | | | |
| Day 0 | 1.64 ± 0.264 | 1.64 ± 0.262 | 1.58 ± 0.237 | 1.73 ± 0.440 | 1.70 ± 0.484 | 2.03 ± 0.519 |
| Day 14 | 10.01 ± 1.005 | 8.39 ± 1.047 | 9.34 ± 0.862 | 10.18 ± 2.356 | 8.67 ± 1.590 | 12.58 ± 2.302 |
| Day 42 | 12.50 ± 1.186 | 10.12 ± 0.602 | 11.20 ± 1.101 | 11.96 ± 1.209 | 9.91 ± 1.657 | 14.55 ± 1.181 |
| ADG, g/piglet | | | | | | |
| Day 0 – 14 | 283 ± 19.5 | 276 ± 19.6 | 263 ± 19.5 | 227 ± 3.8 | 207 ± 26.4 | 223 ± 17.2 |
| Day 14 – 42 | 655 ± 25.5 | 674 ± 23.7 | 698 ± 17.0 | 677 ± 24.4 | 625 ± 28.2 | 625 ± 26.6 |
| Day 0 – 42 | 531 ± 17.8 | 541 ± 17.2 | 553 ± 10.8 | 527 ± 16.6 | 485 ± 26.6 | 491 ± 23.3 |
| ADFI, g/piglet | | | | | | |
| Day 0 – 14 | 293 ± 14.0 | 297 ± 12.9 | 300 ± 13.7 | 255 ± 12.2 | 233 ± 18.3 | 247 ± 17.3 |
| Day 14 – 42 | 915 ± 51.7 | 945 ± 47.1 | 949 ± 29.3 | 888 ± 44.7 | 847 ± 31.0 | 849 ± 30.0 |
| Day 0 – 42 | 708 ± 37.5 | 729 ± 33.7 | 733 ± 21.0 | 677 ± 33.7 | 642 ± 26.3 | 648 ± 25.1 |
| FCR, g/g | | | | | | |
| Day 0 – 14 | 1.06 ± 0.052 | 1.09 ± 0.034 | 1.17 ± 0.060 | 1.12 ± 0.043 | 1.15 ± 0.062 | 1.11 ± 0.012 |
| Day 14 – 42 | 1.39 ± 0.045 | 1.40 ± 0.050 | 1.36 ± 0.033 | 1.31 ± 0.041 | 1.36 ± 0.029 | 1.36 ± 0.018 |
| Day 0 – 42 | 1.33 ± 0.038 | 1.35 ± 0.034 | 1.32 ± 0.025 | 1.28 ± 0.038 | 1.33 ± 0.031 | 1.32 ± 0.017 |
| Faecal consistency score | | | | | | |
| Day 0 – 14 | 6.29 ± 0.082 | 6.42 ± 0.141 | 6.40 ± 0.113 | 5.88 ± 0.172 | 5.58 ± 0.108 | 5.75 ± 0.221 |
| Day 14 – 42 | 6.80 ± 0.082 | 6.83 ± 0.082 | 6.78 ± 0.154 | 6.56 ± 0.120 | 6.38 ± 0.051 | 6.38 ± 0 |
| Day 0 – 42 | 6.58 ± 0.055 | 6.65 ± 0.076 | 6.61 ± 0.103 | 6.26 ± 0.118 | 6.04 ± 0.036 | 6.11 ± 0.095 |

***Supplementary Table S14*** *- The effect of experimental diet on piglet performance between day 0-42 post-weaning, excluding outliers for the second in vivo study at SFR (VOB-62). Data from which outliers were removed are marked by an orange background colour.*

| **Treatment** | **1** | **2** | **3** | | | |
|---|---|---|---|---|---|---|
| Description | Negative control (NC) | NC + LMW chitosan | NC + MMW chitosan | SEM[1] | LSD[2] | *P* - value |
| Body weight, kg | | | | | | |
| Day 0 | 7.65 | 7.65 | 7.65 | 0.002 | 0.005 | 0.80 |
| Day 14 | 11.4 | 11.2 | 11.2 | 0.20 | 0.58 | 0.76 |
| Day 42 | 29.9 | 29.6 | 30.0 | 0.57 | 1.68 | 0.87 |
| CV in body weight, % | | | | | | |
| Day 0 | 1.67 | 1.66 | 1.73 | 0.065 | 0.191 | 0.72 |
| Day 14 | 10.1 | 8.5 | 10.4 | 0.90 | 2.65 | 0.30 |
| Day 42 | 12.3 | 10.1 | 12.3 | 0.89 | 2.62 | 0.15 |
| ADG, g/piglet | | | | | | |
| Day 0 – 14 | 264 | 253 | 250 | 14.2 | 41.7 | 0.76 |
| Day 14 – 42 | 662 | 657 | 674 | 19.1 | 55.9 | 0.82 |
| Day 0 – 42 | 530 | 523 | 533 | 13.7 | 40.1 | 0.87 |
| ADFI, g/piglet | | | | | | |
| Day 0 – 14 | 281 | 275 | 283 | 9.9 | 29.1 | 0.87 |
| Day 14 – 42 | 906 | 912 | 915 | 23.5 | 69.0 | 0.96 |
| Day 0 – 42 | 697 | 700 | 704 | 17.6 | 51.5 | 0.97 |
| FCR, g/g | | | | | | |
| Day 0 – 14 | 1.08 | 1.11 | 1.15 | 0.037 | 0.108 | 0.43 |
| Day 14 – 42 | 1.37 | 1.39 | 1.36 | 0.021 | 0.061 | 0.60 |
| Day 0 – 42 | 1.31 | 1.34 | 1.32 | 0.017 | 0.050 | 0.58 |
| Faecal consistency score[3] | | | | | | |
| Day 0 – 7 | 6.03 | 6.11 | 6.11 | 0.116 | 0.341 | 0.85 |
| Day 7 – 14 | 6.28 | 6.17 | 6.25 | 0.126 | 0.371 | 0.82 |
| Day 0 – 14 | 6.15 | 6.14 | 6.18 | 0.074 | 0.218 | 0.93 |
| Day 14 – 42 | 6.72 | 6.68 | 6.64 | 0.076 | 0.223 | 0.75 |
| Day 0 – 42 | 6.46 | 6.45 | 6.44 | 0.058 | 0.170 | 0.90 |

[1] SEM is standard error of means.

[2] LSD is least significant difference at $\alpha<0.05$ (Fisher's LSD method).

[3] Faecal consistency was registered on a 8-point scale from severe water thin diarrhoea to hard, dry and lumpy faeces. Faecal score 6 was considered the optimal faecal score. Faecal scoring took place three times a week during day 0-14 post-weaning and twice a week between day 14-42 post-weaning and was averaged per experimental period.

***Supplementary Table S15*** - Effect of experimental diet on piglet performance between day 0-42 post-weaning (including outliers).

| **Treatment** | **1** | **2** | **3** | **4** | | | ***P* - value** | |
|---|---|---|---|---|---|---|---|---|
| Description | Negative control (NC) | NC + low dose | NC + medium dose | NC + high dose | SEM[1] | Trt | Linear | Quadratic |
| | | | | Body weight, kg | | | | |
| Day 0 | 9.37 | 9.37 | 9.37 | 9.37 | 0.002 | 0.39 | 0.18 | 0.37 |
| Day 14 | 13.69 | 13.37 | 13.58 | 13.51 | 0.159 | 0.56 | 0.64 | 0.45 |
| Day 42 | 34.00 | 32.65 | 33.30 | 33.53 | 0.610 | 0.48 | 0.78 | 0.21 |
| | | | | CV in body weight, % | | | | |
| Day 0 | 0.89 | 0.96 | 0.94 | 0.80 | 0.060 | 0.24 | 0.29 | *0.09* |
| Day 14 | 7.83 | 7.03 | 8.32 | 8.44 | 0.921 | 0.70 | 0.46 | 0.63 |
| Day 42 | 10.93 | 7.92 | 9.96 | 10.36 | 1.080 | 0.37 | 0.96 | 0.18 |
| | | | | ADG, g/d/piglet | | | | |
| Day 0 – 14 | 309 | 286 | 301 | 295 | 11.3 | 0.55 | 0.62 | 0.46 |
| Day 14 – 42 | 725 | 689 | 704 | 715 | 17.9 | 0.52 | 0.86 | 0.20 |
| Day 0 – 42 | 587 | 554 | 570 | 575 | 14.5 | 0.48 | 0.78 | 0.21 |
| | | | | ADFI, g/d/piglet | | | | |
| Day 0 – 14 | 405 | 379 | 408 | 411 | 13.8 | 0.34 | 0.45 | 0.29 |
| Day 14 – 42 | 1062 | 1002 | 992 | 1017 | 24.6 | 0.22 | 0.20 | 0.10 |
| Day 0 – 42 | 843 | 794 | 798 | 815 | 19.1 | 0.28 | 0.36 | 0.10 |
| | | | | FCR, g/g | | | | |
| Day 0 – 14 | 1.31[a] | 1.34[ab] | 1.36[ab] | 1.40[b] | 0.020 | **<0.05** | **<0.01** | 0.79 |
| Day 14 – 42 | 1.47 | 1.46 | 1.41 | 1.43 | 0.024 | 0.31 | 0.13 | 0.69 |
| Day 0 – 42 | 1.44 | 1.44 | 1.40 | 1.42 | 0.022 | 0.58 | 0.37 | 0.64 |
| | | | | Faecal consistency[2] | | | | |
| Day 0 – 14 | 5.97 | 6.17 | 6.18 | 6.03 | 0.159 | 0.74 | 0.77 | 0.29 |
| Day 14 – 42 | 6.47 | 6.28 | 6.38 | 6.26 | 0.095 | 0.41 | 0.24 | 0.68 |
| Day 0 – 42 | 6.23 | 6.24 | 6.28 | 6.13 | 0.112 | 0.82 | 0.61 | 0.48 |

Different superscripts [(ab)] within a row indicate a significant difference between treatments (P<0.05).

[1] SEM is standard error of mean.
[2] Faecal consistency was registered on an 8-point scale from severe water thin diarrhoea to hard, dry and lumpy faeces. Faecal score 6 was considered the optimal faecal score. Faecal scoring took place three times a week during day 0 – 14 post-weaning and twice a week between day 14 – 42 post-weaning and was averaged per experimental period.

***Supplementary Table S16*** - Effect of experimental diet on piglet performance between day 0-42 post-weaning (excluding outliers).

| **Treatment** | **1** | **2** | **3** | **4** | | | ***P* - value** | |
|---|---|---|---|---|---|---|---|---|
| Description | Negative control (NC) | NC + low dose | NC + medium dose | NC + high dose | SEM[1] | Trt | Linear | Quadratic |
| Body weight, kg | | | | | | | | |
| Day 0 | 9.37 | 9.37 | 9.37 | 9.37 | 0.002 | 0.39 | 0.18 | 0.37 |
| Day 14 | 13.55 | 13.37 | 13.58 | 13.51 | 0.144 | 0.75 | 0.90 | 0.72 |
| Day 42 | 33.49 | 32.65 | 33.30 | 32.95 | 0.481 | 0.63 | 0.66 | 0.62 |
| CV in body weight, % | | | | | | | | |
| Day 0 | 0.89 | 0.96 | 0.94 | 0.80 | 0.060 | 0.24 | 0.29 | *0.09* |
| Day 14 | 7.83 | 7.03 | 8.32 | 9.30 | 0.828 | 0.29 | 0.14 | 0.30 |
| Day 42 | 9.62 | 7.92 | 9.96 | 10.36 | 1.080 | 0.41 | 0.39 | 0.34 |
| ADG, g/d/piglet | | | | | | | | |

| | | | | | | | | |
|---|---|---|---|---|---|---|---|---|
| Day 0 – 14 | 299 | 286 | 301 | 295 | 10.3 | 0.75 | 0.92 | 0.74 |
| Day 14 – 42 | 724 | 689 | 704 | 696 | 15.6 | 0.42 | 0.33 | 0.39 |
| Day 0 – 42 | 587 | 554 | 570 | 561 | 13.1 | 0.35 | 0.30 | 0.37 |
| ADFI, g/d/piglet | | | | | | | | |
| Day 0 – 14 | 395 | 379 | 408 | 411 | 13.0 | 0.29 | 0.19 | 0.46 |
| Day 14 – 42 | 1044 | 1002 | 992 | 998 | 21.5 | 0.33 | 0.14 | 0.28 |
| Day 0 – 42 | 832 | 794 | 798 | 800 | 17.3 | 0.41 | 0.25 | 0.26 |
| FCR, g/g | | | | | | | | |
| Day 0 – 14 | 1.33 | 1.34 | 1.36 | 1.40 | 0.020 | *0.09* | **0.02** | 0.55 |
| Day 14 – 42 | 1.44 | 1.46 | 1.41 | 1.44 | 0.021 | 0.40 | 0.54 | 0.81 |
| Day 0 – 42 | 1.42 | 1.44 | 1.40 | 1.43 | 0.019 | 0.54 | 0.97 | 0.79 |
| Faecal consistency[2] | | | | | | | | |
| Day 0 – 14 | 5.97 | 6.17 | 6.18 | 6.03 | 0.159 | 0.74 | 0.77 | 0.29 |
| Day 14 – 42 | 6.47 | 6.28 | 6.38 | 6.26 | 0.095 | 0.41 | 0.24 | 0.68 |
| Day 0 – 42 | 6.23 | 6.24 | 6.28 | 6.13 | 0.112 | 0.82 | 0.61 | 0.48 |

[1] SEM is standard error of mean.

[2] Faecal consistency was registered on an 8-point scale from severe water thin diarrhoea to hard, dry and lumpy faeces. Faecal score 6 was considered the optimal faecal score. Faecal scoring took place three times a week during day 0 – 14 post-weaning and twice a week between day 14 – 42 post-weaning and was averaged per experimental period.
